# Transient protein structure guides surface diffusion pathways for electron transport in membrane supercomplexes

**DOI:** 10.1101/2024.09.30.615558

**Authors:** Chun Kit Chan, Jonathan Nguyen, Corey F. Hryc, Chitrak Gupta, Kevin Redding, William Dowhan, Matthew L. Baker, Alberto Perez, Eugenia Mileykovskaya, Abhishek Singharoy

## Abstract

The biological significance of protein supercomplexes have remained contentious, particularly how they tune the shuttling of charge-carrier redox proteins across cell membranes during biological energy conversion. We employ multiscale modeling and single particle cryo-electron microscopy (cryo-EM) to determine the mechanisms of diffusive electron transfer in mitochondrial supercomplexes, composed of respiratory complexes III and IV (CIII and CIV). Using a combination of bioinformatic and entropy maximization tools, we model an ensemble of structures representing the conformational space of CIII’s disordered QCR6 ‘hinge’ within the yeast CIII_2_CIV_2_ supercomplex. Molecular and Brownian Dynamics simulations of the entire supercomplex reveal a mechanism for electrostatic coupling between these negatively charged hinge conformations, and binding and directional diffusion of the redox proteins on the mitochondrial membrane, which is simulated over the millisecond timescale. Anionic lipids reinforce this conformationally-coupled recognition of the supercomplex by retaining a pool of the redox proteins in the vicinity of the membrane when the hinges are of a critical length. Cryo-EM models reveal a large-scale rearrangement of the ΔQCR6 supercomplex, which retains a surprisingly robust electrostatic environment for recognition of the redox protein, despite compromise in the supercomplex’s negative charge, still enabling a surface-mediated electron transfer in this CIII_2_CIV_2_ variant. Altogether, the evolutionary need of confining electron carriers on the surface of bioenergetic membranes is found to give rise to a refolding-guided diffusion model of the redox proteins, which improves the energy conversion efficiency within the supercomplex by nearly 30%.

## Introduction

Cellular bioenergetic pathways employ small diffusible redox proteins for the transport of single electrons between neighboring, as well as widely separated, integral-membrane enzymatic complexes. The movement of free electrons between solvated parts of the bioenergetic membranes is achieved only by special redox proteins, not by small molecules as the latter are converted through transfer of single electrons to chemical radicals that cannot be turned loose in the cell. Cytochrome *c* (cyt. *c*) is one such single electron carrier protein capable of protecting electrons from forming reactive chemical species. Cyt. *c* belongs to an exemplary family of redox proteins that participate in the mitochondrial electron transport chain, shuttling electrons, usually only over a short distance [1], between the donor cytochrome *bc*_1_ complex (denoted respiratory Complex III or CIII) and acceptor cytochrome *c* oxidase (respiratory Complex IV or CIV). This electron current through the respiratory chain drives proton transport across the membrane (primarily via the respiratory Complex I or CI), generating an electrochemical gradient of ∼200 meV that is ubiquitously employed to synthesize ATP and power all aerobic cells [2].

The role of physical proximity of the charge donor and acceptor complexes in tuning the transport efficiency of the carrier proteins, and hence overall energy turnover of the cell has remained elusive over the past two decades now. Native gel electrophoresis together with cryo-tomography studies of bioenergetic membranes in mammals, yeast and plants revealed that the electron-transport chain components needed for ATP production are organized in a dynamic equilibrium between physically dispersed and closely packed states [3,4]. These clusters of the interacting respiratory complexes with variable stoichiometries are denoted as *mitochondrial supercomplexes* [5]. A general formula of the supercomplexes is written as CI*_x_*CIII*_y_*CIV*_z_*, where *x* → 0, 1; *y* → 2; *z* → 1, 2. For the special case of *x* → 0, a supercomplex is reduced to CIII*_y_*CIV*_z_* that we study here [5,6]. The fraction of CIII and CIV in the free *vs.* associated forms determines the redox state of the overall cyt. *c* pool, which is expected to tune the mechanisms of charge transfer in response to changes in environmental conditions and cellular redox-signaling pathways [7,8]. However, little is known about the detailed mechanism of such regulation.

Consequently, the biological relevance of supercomplexes has been questioned. A number of views have transpired. The generally agreed upon finding is that protein association reduces the probability of electron leakage, leading to a controlled formation of reactive oxygen species [7,9]. Also, the supercomplexes can avoid random aggregation of the proteins in the membrane [10,11]. However, the issue that has generated an exuberant discussion is whether the respiratory supercomplexes increase the efficiency of cellular energy conversion by reducing the physical distance between the individual complexes [12,13]. Central to addressing this puzzle is the definition of a supercomplex’s efficiency in terms of thermodynamic and kinetic parameters. Enhanced thermodynamic efficiency would entail an increase in the yield of ATP from the supercomplex-rich electron transport chain. But, despite several biophysical and biochemical studies, such enhanced ATP yields have not been observed [3 ,14–16].

Alternatively, a combination of low-resolution electron microscopy data together with theoretical rate models have suggested kinetic advantages of supercomplex formation towards expediting cyt. *c* turnover between CIII and CIV if three-dimensional diffusion is the dominant mechanism of transport [3,4,11,16–19]. Even this interpretation becomes convoluted with the possibility of a two-dimensional cyt. *c* diffusion that can uniquely leverage the densely packed supercomplex environment [4], and potentially compete with the three-dimensional diffusion (**Fig. 1**A-B). In particular, the detailed molecular interactions underpinning the two- and three-dimensional diffusion pathways of cyt. *c* are yet unresolved.

**Fig. 1:**
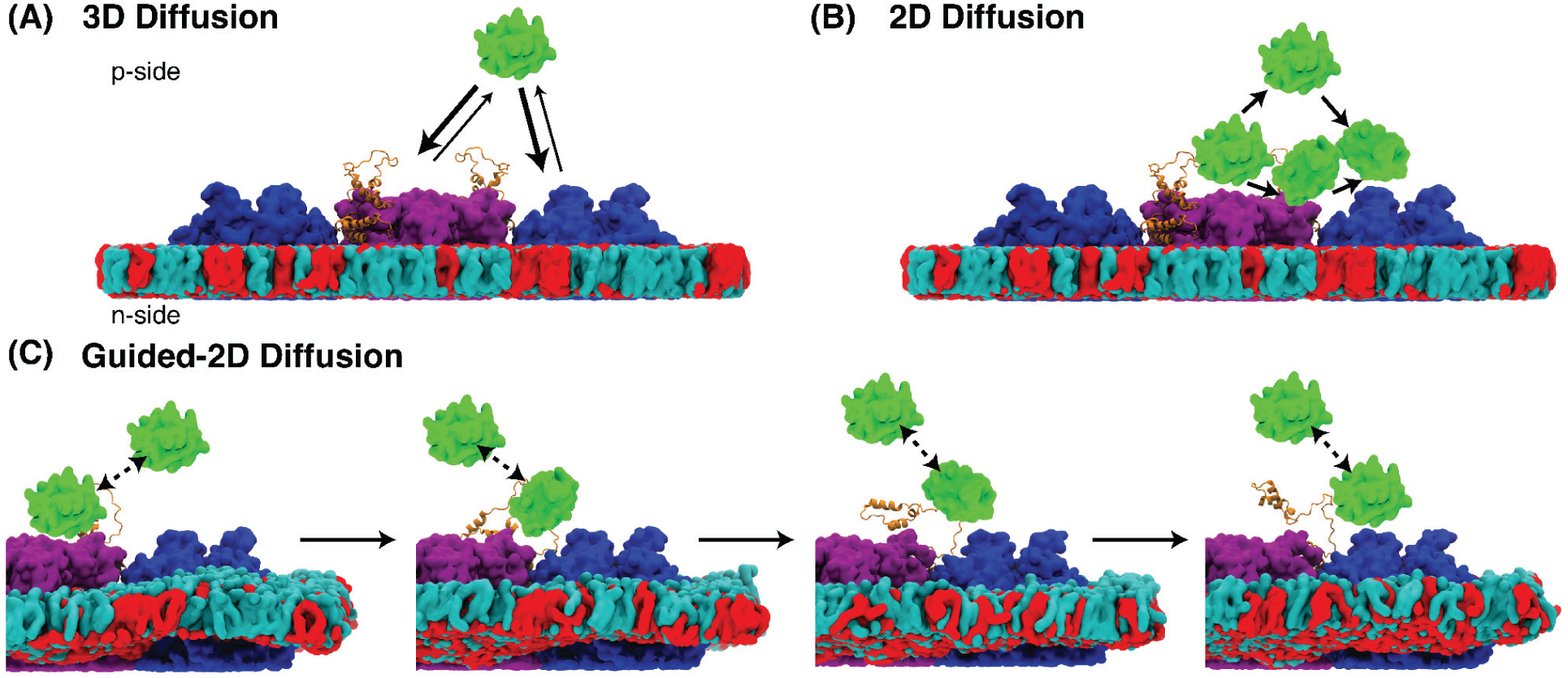
Possible mechanisms of cyt. *c* diffusion across a supercomplex. **(A)** Cyt. *c* (green) undergoes 3D diffusion, where the protein exchanges between the bulk solution and the surface to transfer charges between complex III (purple) and complex IV (blue) [59]. **(B)** Cyt. *c* undergoes surface or 2D diffusion, where, as suggested in [1], the protein stays in the vicinity of the supercomplex during its translocation between complex III and IV. **(C)** Cyt. *c* can undergo guided diffusion, where the protein’s translocation is directed by the conformation of the disordered domain of QCR6 (orange), a hinge-like subunit of CIII. In figures below, the anionic mitochondria membrane is represented by our model membrane with anionic lipids (here, cardiolipin molecules) in red and charge-neutral lipids (here, POPC) in cyan.

Two understudied molecular components that play a major role in the kinetics of diffusive cyt. *c* transport are **(1)** the anionic lipids e.g. cardiolipins in the membrane, and **(2)** the highly negatively charged QCR6 hinge of either an isolated or supercomplex-bound CIII. While the anionic lipids are already known to enhance the stability of mitochondrial complexes [20–24], their organization within the yeast CIII_2_CIV_2_ supercomplex is revealed recently in our published cryo-EM models (PDB: 8E7S) [25], and also in more recent models [23,26]. Cardiolipins offer a reconfigurable interaction network between individual complexes, which allow association of the complexes into supercomplex structures that may be required to regulate cyt. *c* dynamics in response to varying physiological conditions [20]. Hence, lower cardiolipin levels are associated with reduced formation of supercomplexes in a number of neurodegenerative diseases, ischemia followed by reperfusion, induction of apoptosis, heart failure, obesity and aging [24,27].

QCR6, on the other hand, bears structural similarity to the twin CX_9_C proteins, having two canonical α-helices joined by only one disulfide bond into a ‘hinge’ shape. The N-terminal region of QCR6 is conformationally transient and has no reported structure thus far, even within the yeast supercomplex model at 3.2 Å resolution [25]. It contributes to functional roles in preserving the heme environment of cyt. *c*_1_ inside CIII and promotes interaction with cyt. *c* [28,29]. Consistent with this function, cells lacking QCR6 block maturation of cyt. *c*_1_, attenuating CIII catalytic activity and growth [28]. Thus, despite not being a requisite for supercomplex formation, QCR6 directly contributes to the mechanism of cyt. *c* turnover. Recent findings and the identification of disease-associated mutations in specific CX_9_C motif-carrying proteins have highlighted members of this family of proteins as potential therapeutic targets in a variety of human disorders [30].

Here, we seek to resolve the volume-dependent (3D) *vs.* surface-dependent (2D) mechanisms of cyt. *c* transport in the mitochondrial environment by modeling their hinge-mediated diffusion across the CIII*_y_*CIV*_z_*-type supercomplexes under varying QCR6 composition and dynamics, anionic lipids concentration in the membrane, and association architectures. To this end, we employ a combination of cryo-electron microscopy (Cryo-EM) density maps with protein folding methods (Modeling Employing Limited Data or MELD), all atom and accelerated molecular dynamics (MD) simulations of the entire CIII_2_CIV_2_ construct [31,32]. This interplay between membrane interactions, and transient order and hinge-like movement of QCR6 is further interrogated with continuum electrostatics and Brownian Dynamics (BD) simulations [33]. These approximately millisecond-scale computations offer a working model of how the supercomplex architecture tunes cyt. *c* movements between CIII to CIV from random 3D to 2D diffusion onto folding/unfolding-guided 2D diffusion (**Fig. 1**C). Integration of the simulated data into a master equation framework quantifies the kinetic advantage of energy conversion with surface-guided electron transport in a supercomplex environment over the volume-guided mechanism in freely dispersed complexes. We further investigate how supercomplexes reorganize the surface electrostatics potentials for charge-carrier recognition and transport even when QCR6 is deleted, hence maintaining a fairly robust environment for 2D diffusion dynamics to dominate biological energy conversion. Finally, our models offer insights on differences between conducting charge transport with soluble carriers in higher organisms over utilizing their membrane-anchored counterparts in more primitive bioenergetic membranes.

## Results

In what follows, first our cryo-EM model of the CIII_2_CIV_2_ supercomplex is augmented with an ensemble refinement of the QCR6 structure. Second, the dynamics of QCR6 is probed under different electrostatic environments as it undergoes conformational transitions to control the transport of cyt. *c* from bulk solution to CIII, and thereafter between CIII and CIV. Third, the interplay between supercomplex stoichiometry, membrane compositions and QCR6 dynamics is explored to determine the probability of 3D *vs.* 2D diffusion of cyt. *c*. Finally, in view of the existing structural, biochemical and computational data, a rate-kinetic model of soluble protein transport is proposed to outline advantages of energy transfer in confined spaces.

### The QCR6 domain of yeast Complex III can exist in transiently folded conformations

There are no known structures of the complete QCR6 domain of CIII, it displays a diversity of lengths across species with yeast engendering one of the longest sequences (**<u>Fig. SI1.1</u>**) [34]. This unusual property of the yeast CIII prompted our ensemble refinement of the QCR6’s N-terminal hinge region, starting from the cryo-EM supercomplex model from Saccharomyces cerevisiae. The cryo-EM model of CIII_2_CIV_2_ used for this study was reported in the PDB: 8E7S, which is resolved at 3.2 Å and builds upon previously resolved CIII and supercomplexes (PDB: 6YMX; 1KB9; 3CX5; 6Q9E; 6GIQ). As in all the models, after purification, the yeast supercomplex has CIII bound to one or two CIV in a triangular trimeric or linear tetrameric shape, yet with comparable CIII-CIV interfaces. In the 8E7S model, these interfaces are found to be stabilized by anionic lipid molecules, with the density maps revealing ten distinct cardiolipin binding sites that were identified using POPG lipids.

To build a complete QCR6 structure within the supercomplex, first, a reduced cyt. *c* was docked on the p-side of the CIII_2_CIV_2_ supercomplex by leveraging existing X-ray crystallography data on the cyt. *c*-docked CIII structures (see Methods: **<u>Initial model preparation</u>**) [35]. Thereafter, the MELD-guided MD simulations (see Methods: **<u>MELD</u>**) revealed an ensemble of structures for the extended QCR6 N-terminal in the presence of the supercomplex-bound cyt. *c* (**Movie 1**). Illustrated in **Fig. 2A**, the most probable folded state determined at room temperature, is composed of five helices with a total of 73 residues. Two of these helices are stabilized by positive charges on cyt. *c*, and are connected to the other three, which are proximal to the membrane via a 40-residue long loop.

**Fig. 2:**
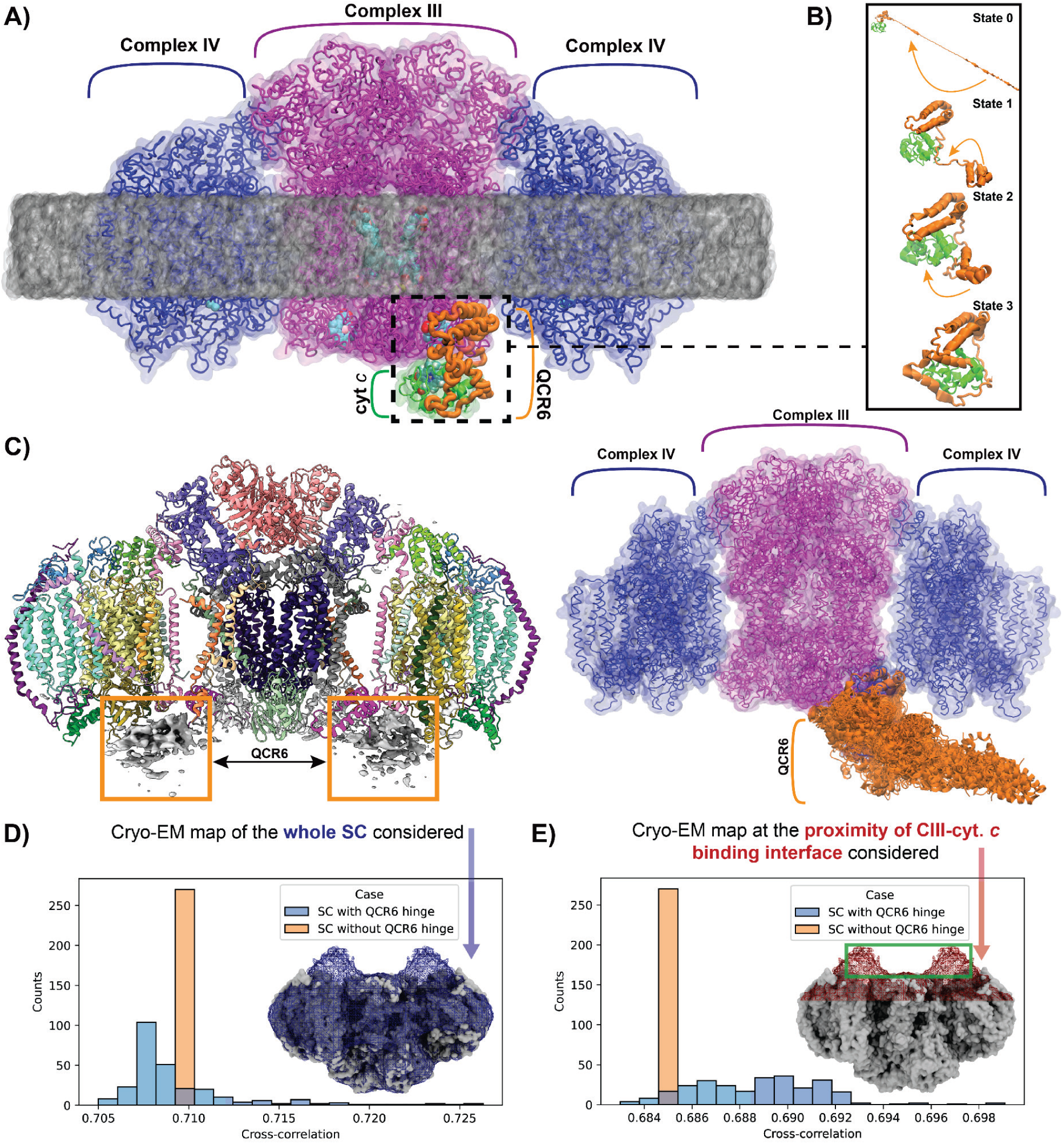
Simulated structural model of QCR6 domain in CIII. **(A)** An all-atom model of the yeast supercomplex composed of electron donor CIII dimer (purple) associated with two copies of electron acceptor CIV (blue) and one carrier cyt. *c* (green) embedded in a POPC model membrane. **(B)** A starting model for the QCR6 N-terminal region is constructed by folding its first 76 residues using MELD-accelerated MD simulations in the presence of cyt. *c* and soluble residues from CIII that are within 14 Å of the cyt. *c*. Four snapshots (denoted states 0–3) representing the local interaction energy minima (see **<u>SI1.4</u>**<u>B</u>) along the protein folding simulation are illustrated, demonstrating the flexibility of the QCR6 hinge [31]. **(C)** (Left) Reconstruction of the supercomplex using non-uniform refinement shows strong density features (white map) located beneath the QCR6 (magenta chain). (Right) Supercomplex with an overlay of the disordered QCR6 hinge, encompassing models with an end-to-end distance of ∼115Å or shorter (see states between 2 and 3 in (B)) occupies a similar space in MELD-accelerated MD simulation snapshots as that observed in the refined density map. The impact of including our modeled and simulated QCR6 (with the acidic hinge region) to the partially resolved supercomplex was accessed by measuring the cross-correlation coefficient structure and 2 different cryo-EM maps. **(D)** Correlation between the supercomplex with/without QCR6 hinge region and the full cryo-EM map (shaded blue) was measured. Because of the flexible nature of the QCR6 hinge region, a slight drop in correlation is observed upon including the QCR6 hinge region. **(E)** Correlation coefficient is recomputed with/without QCR6 hinge region and the fraction of cyro-EM map around the CIII-cyt. *c* binding interface (shaded red) is measured. The inclusion of QCR6 hinge region noticeably improves the resulting ccc as the QCR6 hinge region fills the map region where not a single atom has been resolved (highlighted by a green rectangle)

To verify this predicted motif, we performed an Alphafold2 modeling of the QCR6 sequence both in the presence as well as absence of the bound cyt. *c*. In the absence of cyt. *c* there was minimal structure remaining in this highly negatively charged sequence, including the long disordered loop with –CE_9_C– residues. However, in the presence of cyt. *c*, the confidently resolved parts of the Alphafold2 solution, namely with a confidence threshold of pLDDT > 90, were found to be within an RMSD of 2-6 Å of the converged MELD model (**<u>Fig. SI1.2</u>**). This analysis brings to light a transient nature of the QCR6 structure, particularly in the yeast-specific extended loop, which folds due to the presence of complementary cyt. *c* interfaces. Very recently, there is evidence of such cyt. *c* - induced QCR6 folding from cryo-EM studies. [36]. Furthermore, k_D_ estimates of the QCR6-updated CIII-cyt. *c* (determined from the PRODIGY server) is roughly 0.2 µM (**<u>Fig. SI1.3</u>**), very much in the range of values of ∼0.3 µM determined from plasma resonance microscopy [37]. In the absence of QCR6 in our model, this value decreases to ∼0.4 µM in our previous work, reinforcing the importance of this unresolved region in cyt. *c* binding to CIII [38].

Going beyond the static Alphafold2 models, our MELD simulations further visit at least three partially folded low-energy conformations during the QCR6 folding (**Fig. 2B**). These intermediates elucidate the conformational space that QCR6 can thermally access at room temperature by varying its helical content (estimated in **<u>Fig. SI1.5</u>**). Within the folding intermediates seen by MELD, end-to-end distances of the extended QCR6 loop varies between 35 Å to 26 Å (denoted State 3), starting from a fully extended loop of length 175 Å (State 0, **Fig. 2B**). By comparing this stretch to the separation between the heme group of electron donor CIII and the copper group of electron acceptor CIV, which ranges up to 115 Å from the cryo-EM models of the supercomplex, we infer that a fully stretched QCR6 can potentially interact with CIV, and most certainly interact with the upper (p-side) leaflet of the mitochondrial membrane. Further processing of the cyt. *c*-free supercomplex data (EMD-27940) with a nonuniform refinement (Methods: **<u>Non-uniform refinement of WT supercomplex</u>**) brought to light delocalized unresolved density features grazing the surface of CIII and CIV, which cannot be interpreted by the structured subunits (**Fig. 2C**). Interestingly, a cross correlation analysis reveals that inclusion of the folded QCR6 N-terminal hinge from the MELD simulations improves match between the structural ensemble and the unresolved density features (**Fig. 2D-E**). Together with outcomes of the MELD simulation, now, we take this unresolved density for an ensemble representation of the dynamic QCR6 N-terminal.

### Direct QCR6 interactions with anionic lipids regulate cyt. *c* movement between electron donor and acceptor complexes

The variance of QCR6 conformations cannot be accessed using brute-force MD, even with microsecond-long simulations. This is due to the longer timescales over which its folding-unfolding transitions and associated changes in the cyt. *c* structure occurs. Following an initial thermalization with 1 µs of conventional MD of the entire membrane-embedded CIII_2_CIV_2_ supercomplex, three copies of 500 ns long Steered MD (SMD) simulation were performed (see Methods: **<u>MD and SMD</u>**). Notably, the membrane is composed of POPC, and POPG lipids in the 8E7S model were replaced with cardiolipin (C18:2) molecules. During these SMD simulations, the biasing potential only acted on the distance between the center of mass of the alpha carbons of cyt. *c* and those of residues (TYR146, ASP150, GLU151, ASP162, ASP164) describing a putative cyt. *c* binding site on the CIV surface [39]. The QCR6 hinge (i.e. the flexible domain of QCR6) was, thus, capable of relaxing in any direction other than that of the steering. Such relaxation allowed the QCR6 to stretch and concomitantly interact with the membrane leaflet as well as with the cyt. *c*, which was translocating from CIII→CIV (**Fig. 3**).

**Fig. 3:**
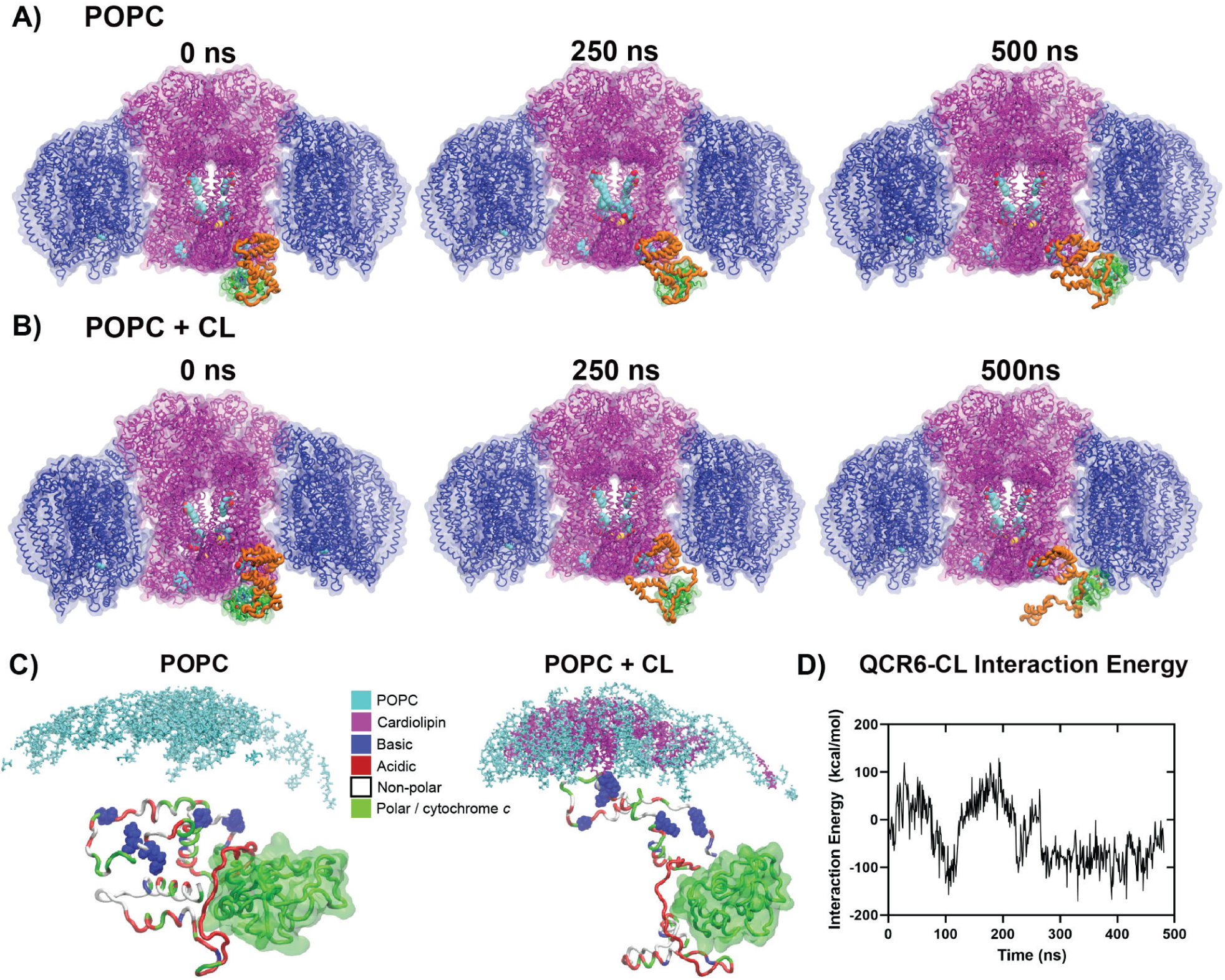
Transient QCR6 conformations enable direct interactions with cardiolipins. Structural snapshots from 500 ns of SMD simulation of cyt. *c* movements from CIII → CIV in the absence **(A)** vs. presence of cardiolipin **(B)**. Movement of cyt. *c* is guided by unfolding of the QCR6 hinge (Movie 2), which is further verified using GaMD simulations (**<u>Fig. SI1.5</u>**). Hemes are shown in VdW representation inside CIII in (A) and (B). **(C)** The basic residues (Arg104 and Lys106) on QCR6 hinge closest to the membrane surface form salt bridges with the anionic cardiolipins, while QCR6 N-terminal simultaneously aiding in cyt. *c* transport. **(D)** Favorable QCR6-CL interaction energies that emerge half way along the transport pathway, ∼250 ns, and remain stable till the cyt. *c* reaches CIV.

Following translocation from CIII, binding of the cyt. *c* to the Cu_A_ pocket (or hotspot) of CIV was confirmed using dissociation constant (k_D_) computations of the cyt. *c-*CIV complex models resulting from SMD simulations (**<u>Fig. S1.3</u>**). The computed affinities are ∼0.1 μM in the range reported in the 0.01 - 1 μM range surface plasmon resonance measurements [40–41]. Such systematic underestimates of absolute protein affinities have also been noted in the past [42]. Despite this discrepancy, the CIV binding energies are found to be at least two-fold stronger than those from cyt. *c-*CIII. This trend remains in line with the observation that cyt. *c* turnover at CIV (and not CIII where k_D_ is almost twice weaker) offers one of the energy bottlenecks of electron transport [1,19]. Such stronger affinity of a reduced cyt. *c* for an oxidized CIV over oxidized CIII seen in our computations, further supports the forward directionality of electron flow in the ATP synthesis direction, and avoids backward leakage.

Remarkably, the cardiolipin headgroups in the membrane form an electrostatic interaction with the Arg104 and Lys106 residues of the folded QCR6 domain. The interaction saturates around -100 kcal/mol (**Fig. 3**). So, these interactions tether the QCR6 hinge to the membrane while the cyt. *c* approaches the Cu_A_ binding pocket of CIV. The SMD simulations were repeated in the absence of the CL in the membrane. Now, the average work required to transport cyt. *c* from CIII to CIV increased by 25-30 kcal/mol (**<u>Fig. SI2.1</u>**). Hence, the electrostatic coupling between the QCR6 and the anionic lipids of the membrane cuts down the cost of cyt. *c* transport across the supercomplex. A sequence alignment analysis further reveals that the Arg104 residues, which are found to interact with cardiolipin headgroups, are conserved across five different species, notwithstanding the heterogeneous length of the hinged region (**<u>Fig. SI1.1</u>**).

Conservation of the arginine residue further suggests that QCR6-assisted transport of cyt. *c* between the donor and the acceptor complexes can be ubiquitous to the electron transport chain, potentially beyond the yeast membrane. Also, the cardiolipin tails were already resolved in the supercomplex structure, highlighting the importance of protein-lipid interactions in offering stability [13,22]. Our simulation together with this structural information now suggests that the membrane does not just offer stability to the supercomplex, the anionic lipids directly interact with transient conformations of CIII’s disorder regions, namely the QCR6 loop to energetically favor cyt. *c* transport across the supercomplex surface.

### Surface-guided diffusion model for cyt. *c* transport

Gaussian accelerated MD (GaMD) simulations were performed to explore the secondary structures of the extended QCR6 during the cyt *c* transport from CIII to CIV. 400 ns of GaMD simulations, each starting from the SMD snapshots of **Fig. 3** revealed that, in the absence of the cyt *c*, the QCR6 remains in a conformational pre-equilibrium between folded and unfolded states, revealing two distinct population of helicities (**<u>Fig. SI1.5</u>**). In the presence of cyt. *c*, the QCR6 remains more tightly packed in the vicinity of CIII, but unwinds as it approaches CIV, akin to the free form. Such a dynamical pre-equilibrium, together with an overall high negative charge of QCR6’s N-terminal domain (−37 e), can underpin the disorder and radiation damage that has kept the piece of the protein unresolved via cryoEM.

Almost 90% of the mitochondrial electron transfer proteins remain in supercomplex states within yeast, as opposed to only 35% of complexation propensity in mammals [1,43–44]. The longer QCR6 of the yeast CIII (**<u>Fig. SI1.1</u>**) offers a clear functional advantage in the densely packed supercomplex environments. A 2D movement of the charge carrier cyt. *c* was suggested based on low-resolution structural evidence [1,4]. Our model illustrates that such movement represents not just surface diffusion of the carrier, but explicit carrier-anionic lipid interactions mediated by the QCR6. Hence, the cardiolipins contribute on one hand to the stability of the supercomplexes, as highlighted in the cryo-EM structure, but more importantly, directly interact with transient conformations of CIII’s disordered regions to energetically favor cyt. *c* displacement across the supercomplex surface (**Movie 2**). This interplay between protein-lipid and protein-protein interactions complements the utility of a relatively longer QCR6 in yeast supercomplex. The results, however, do not imply that the QCR6 enforces the cyt. *c* to act like a structural component of the supercomplex with any particular stoichiometry. Rather, the membrane interaction with the stretchable acidic region of QCR6 keeps the transporting cyt. *c* in the vicinity of both the donor and the acceptor, with exchanges of the cyt. *c* still possible between the surface and the bulk solvent (**<u>Fig. SI3.1</u>**).

### Supercomplex recognition and surface localization of the cyt. *c* pool is reinforced by collective electrostatics of QCR6 and anionic lipids

It is now established that the supercomplex formation has minimal bearing on the rate-limiting step of the mitochondrial electron transfer chain [1,4]. Our results of **Fig. 3** and **<u>Fig. SI1.3</u>** are in line with such observation, in that we find the QCR6 in supercomplexes to only improve long-range interactions. Whereas, the rate-determining step associated with cyt. *c* or *c*_2_ binding is dominated by the entropy bottlenecks imposed via the short-range interactions, namely via a highly constrained half-ring distribution of complementary hydrophobic residues [34,45]. Such short-range interactions remain similar between the supercomplex and individually dispersed systems. The mechanism for molecular recognition, nonetheless, plays a major role to ensure that the cyt. *c* pool associates with the supercomplex and not unproductively interacts with the membrane surface, particularly the anionic lipids. So, we seek how QCR6-augmented and QCR6-removed (or ΔQCR6) supercomplex environments promote the feasibility of productive charge transfer.

#### Role of QCR6 electrostatics

Atomic Resolution BD simulations were performed to simulate the diffusive dynamics of cyt. *c* in the vicinity of the supercomplex at different stages of its CIII → CIV transport (**Fig. 4**). Starting models for the BD simulations were extracted from the energetically favorable conformations of the SMD simulation (see Methods: <u>BD</u>). The BD simulations offer a broader range of cyt. *c*-bound CIII and CIV models than SMD, which reveals a unimodal distribution of affinities for the former, and a bimodal distribution of relatively loosely bound models for the later (**<u>Fig. SI1.3</u>**). This result is in line with the presence of separate long-range recognition and electron-transfer complexes in CIV, the former being weaker with a k_D_ of 0.4-0.7 μM and the latter with a stronger k_D_ of 0.1-0.2 μM, while only a single productive complex is observed in CIII [40–41].

**Fig. 4:**
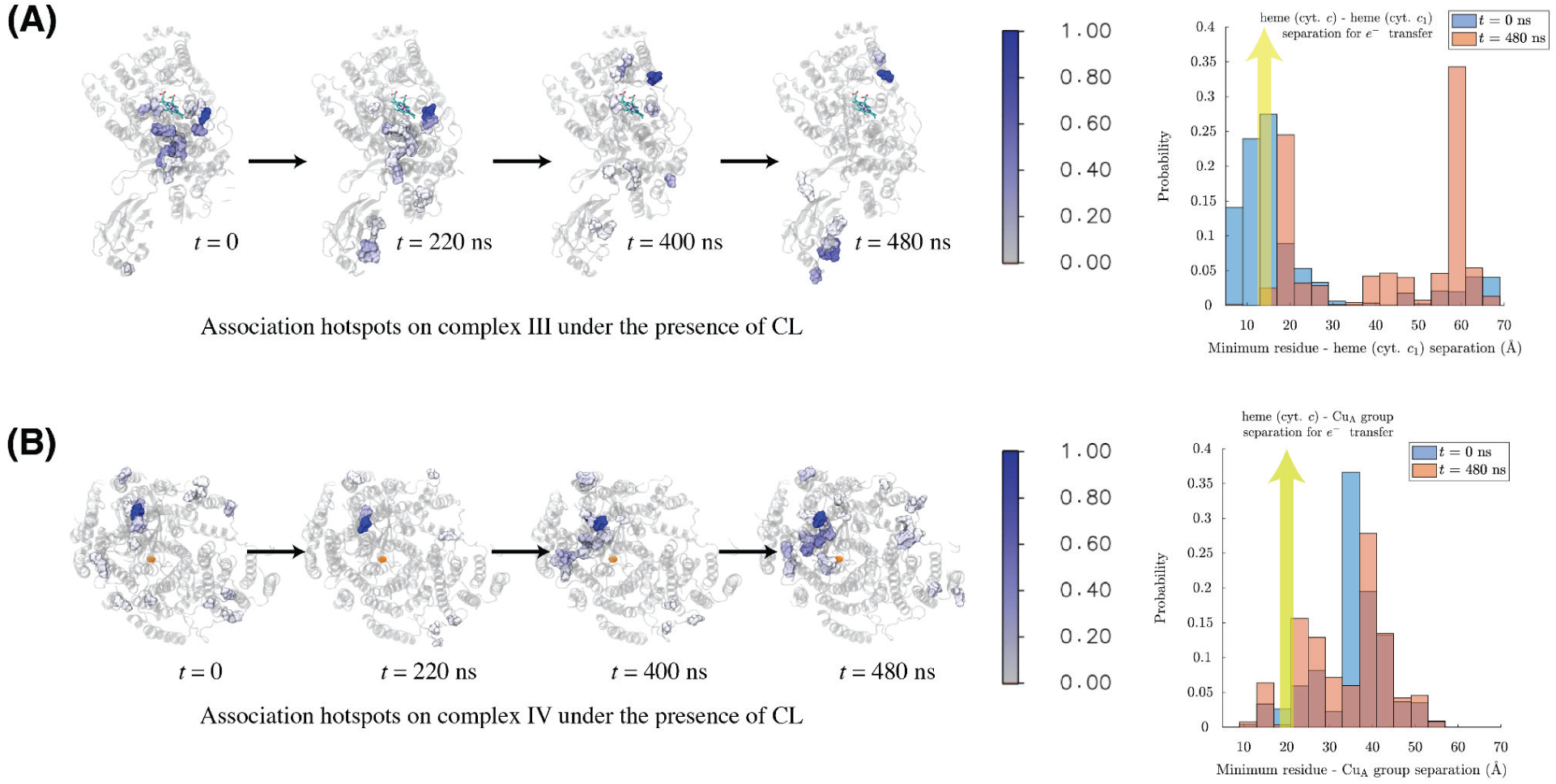
Conformations of QCR6 regulate diffusive cyt. *c* association on CIII and IV hotspots. (A) Primary binding sites for cyt. *c* associations on CIII, in the presence of cardiolipins. Residues of CIII are colored according to the colorbar on the right, where the scale indicates percentage occupancies. The percentage occupancy of a residue is the ratio of its number of cyt. *c* contacts (i.e. the number of events when cyt. *c* is within 6 Å of the residue) with the maximum number of cyt. *c* - CIII contacts (cyt. *c* is within 6 Å of all the CIII surface residues) observed during simulations. So, when the occupancy of a residue with occupancy 1,the most frequent cyt. *c* - interacting residue throughout the molecular trajectory. (B) Similar molecular representation highlighting binding to the CuA group, the cofactor in CIV for receiving electrons from cyt. *c*, serves as the reference for distance measurements. (C-D) Distributions of the separation from each cyt. *c* - interacting residue, highlighted in the corresponding molecular image, to the respective cofactors for electron transfer are plotted. The data shown here for each residue is weighted by its percentage occupancy, where a higher percentage occupancy contributes a higher weight (see Methods: <u>Analyzes details</u>). As the SMD snapshots evolve from *t* = 0 ns to 480 ns, BD revealed a decrease in the cyt. *c* occupancy closer to the CIII interface (distance of cyt. *c* heme to the heme *c*1 of CIII, ≤14 Å, indicated by the yellow arrow), and growth in the vicinity of CIV (distance of cyt. *c* heme to the CuA group of CIV, ≤ 20 Å [39], indicated by the yellow arrow).

Illustrated in **Figs. 5**A-C and **<u>SI4.1</u>**, we find that the cyt. *c* grazing on the surface of the supercomplex via at least three different pathways. The cyt. *c* - QCR6 interactions are stronger than that of its membrane affinity. This is facilitated by the additional negative charge on the QCR6 N-terminal that complements the seven Lys residues on cyt. *c* surface. Consequently, the number of cyt. *c*’s associations/Å^2^ surface area of the supercomplex (Method: <u>Computations of binding propensity per unit area</u>) is 2-3 fold more than that of the cyt. *c*-membrane association, leaving on ∼10% of the cyt.*c* in the bulk (**Fig. 5**D-E) for exchanging with the surface-associated ones. As the QCR6 unfolds and stretches towards CIV, a cloud of cyt. *c* (quantified as occupancy, Method: <u>occupancy calculation</u>) moves away from the hotspot residues of CIII, which are close to the protein’s cofactor (heme), towards the Cu_A_ group of CIV **(Fig. 4)**. Consequently, cyt. *c* switches from being localized around CIII surface to being localized around CIV (**<u>Fig. SI4.2</u>**). This finding on surface localization of cyt. *c* is also in agreement with the kinetic experiments [4,46], wherein exclusive 3D diffusion is excluded as the most probable mode of cyt. *c* transfer in the electron transport chain. Worth noting, the directional bias in cyt. *c* transfer is affected when the anionic lipids are removed (**<u>Fig. SI4.3</u>**). Now, the hotspot residues for cyt. *c* occupancy are more evenly distributed between CIII and CIV, despite the presence of QCR6, creating ambiguity about the site of electron acceptance. So, the dependence of directional cyt.*c* occupancy on membrane composition alludes to a possible cooperativity between QCR6 activity and anionic lipids for sustaining the carrier protein’s surface accessibility, which will be further dissected below.

**Fig. 5:**
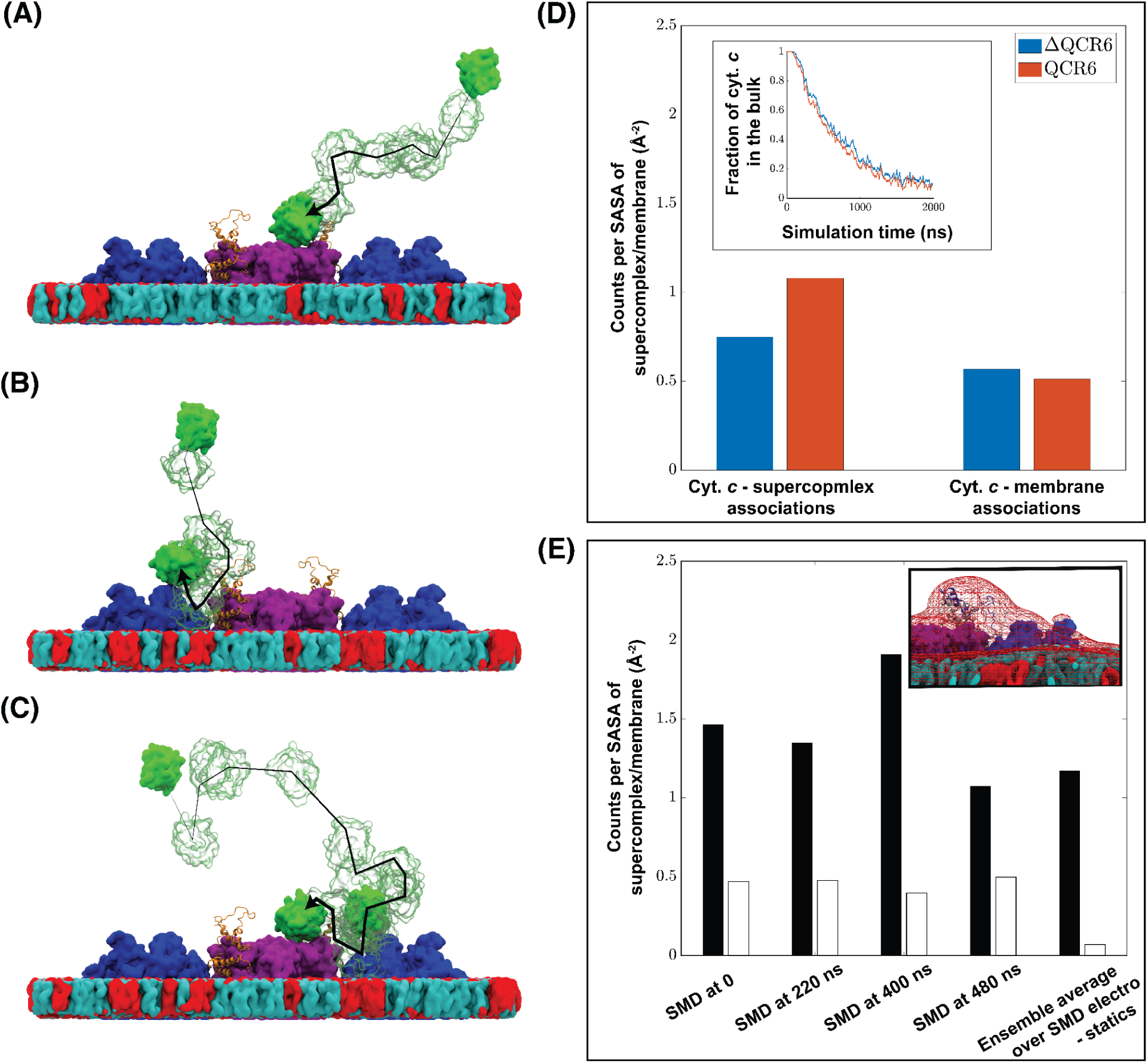
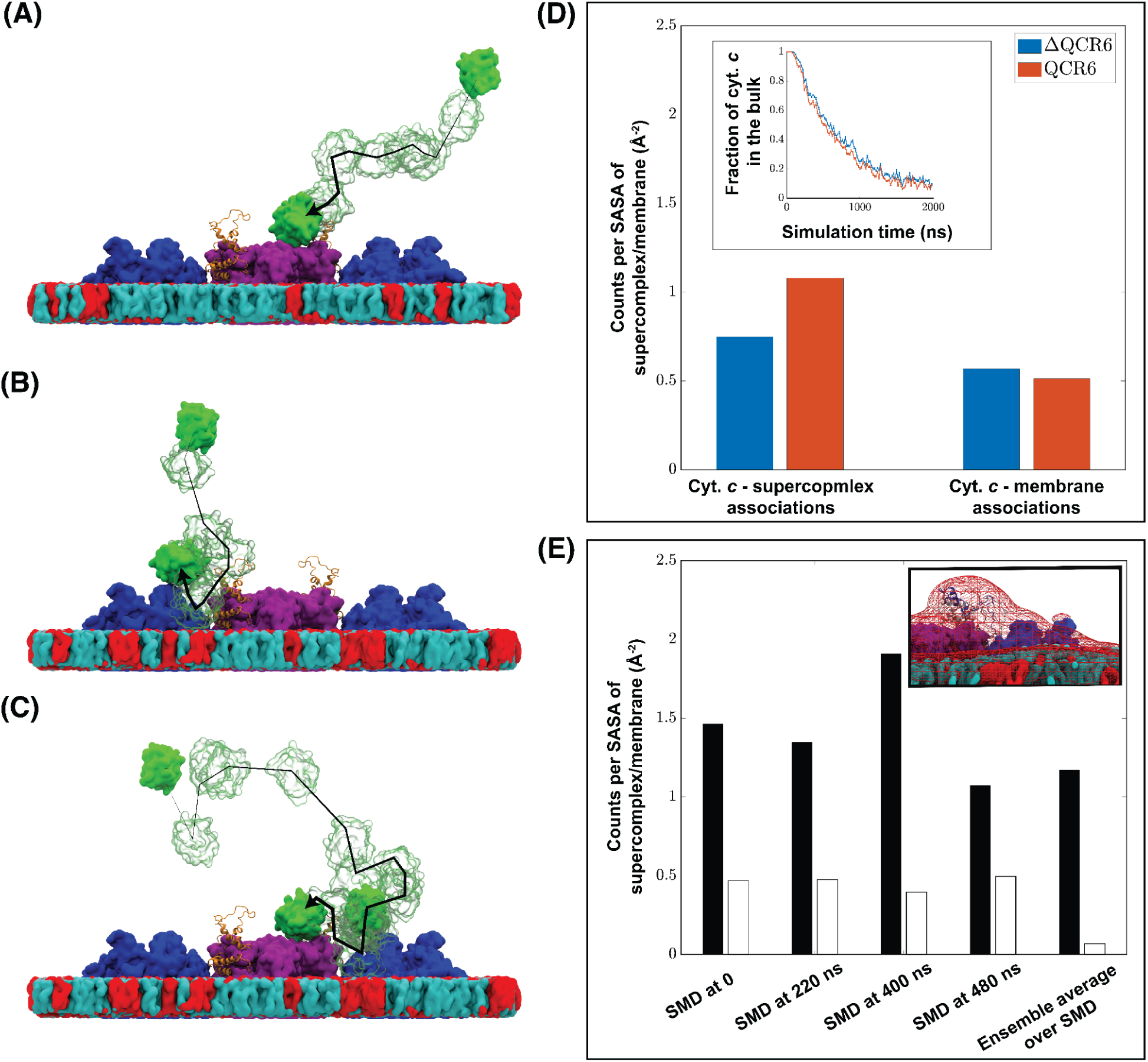
QCR6 promotes cyt. *c* - supercomplex association via multiple diffusion pathways. Cyt. *c* (green) diffusion follows multiple pathways during BD simulations: **(A)** diffusing entirely in the bulk solvent and arriving at the supercomplex; **(B)** diffusing to the membrane then to the supercomplex; and **(C)** encountering the supercomplex then translocating between CIV and CIII through subsequent diffusion over either the membrane or QCR6 or both. These different diffusion pathways are indicated in **(A)**, **(B)**, and **(C)** by black arrows tracing snapshots of cyt. *c* (green, transparent) captured in corresponding BD simulations. Color-code for CIII (purple) and CIV (blue) are maintained, while POPC and cardiolipin are indicated in cyan and red. The number of cyt. *c*’s associations with the supercomplex and that with the cardiolipin-containing membrane, normalized to their respective solvent accessible surface area (SASA), are shown in **(D)** for the complete supercomplex model, namely the WT and is labeled QCR6 here, and the one without QCR6s’ disordered hinge domains, labeled ΔQCR6. The fraction of cyt. *c* remaining in the bulk solution along the simulation time is given in the inset. The number of cyt. *c* associations with the supercomplex and that with the membrane for the supercomplex at different time points of the SMD simulation, as well as for a thermally averaged electrostatic maps from all of these QCR6 conformations called “Ensemble average electrostatic”, are shown in **(E)**, where data for cyt. *c* - supercomplex associations are shown by black bars and those for cyt. *c* - membrane associations are given in white bars. A molecular representation of this cyt. *c* - membrane association average model is given in the inset (details at **<u>Fig.</u> <u>SI4.1</u>**).

The ΔQCR6 reduces the CIII-CIV recognition surface of the cyt. *c* from 1.1 cyt.*c*/Å^2^ of surface in the wild-type supercomplex to 0.7 cyt.*c*/Å^2^. In ΔQCR6, marginally more cyt. *c* interacts with the membrane in the vicinity of the supercomplex (from 0.5 cyt.*c*/Å^2^ to 0.6 cyt.*c*/Å^2^ **Fig. 5**D), while very little change happens to the bulk cyt. *c* concentration (**Fig. 5**D<u>-inset</u>). Noting that the surface area of the membrane is 7.9-fold larger than the supercomplex model, even the minor increase in the probability is adequate to shift a significant population of the cyt. *c* towards the weakly interacting membranes. Also, cyt. *c*’s residence on the supercomplex at regions close to to the electron-transferring cofactors of CIII and CIV for forming productive complexes (**Fig. 4**) becomes shorter-lived in ΔQCR6 relative to WT (**<u>Fig. SI4.4</u>**<u>A</u>). Thus, productive cyt. *c*-supercomplex interaction is compromised in the ΔQCR6 models. On the contrary, cyt. *c*’ s residence time on other parts of the supercomplex as well as on the membrane are similar between ΔQCR6 and WT (**<u>Fig. SI4.4</u>**<u>B</u> and <u>C</u>).

Rate matrices determined from the BD simulation following partition of the space into four regions (CIII, CIV, membrane, and the bulk in **<u>Fig. SI3.2</u>**<u>A</u>) reveals that the transition of cyt. *c* from CIII to CIV occurs noticeably differently between ΔQCR6 and WT. ΔQCR6 is able to achieve an initial population transfer rate from bulk to the surface that is around 2-fold higher than that for WT. Yet, ΔQCR6 reduces the transfer of the cyt. *c* population between the donor and acceptor complexes, resulting in a lower equilibrium cyt. *c* population on CIV (**<u>Fig. SI3.2</u>**<u>B</u>). As a result, the effective transfer rate of cyt. *c* from CIII to CIV for ΔQCR6 (0.63µs^-1^) is lesser than that of WT (0.96µs^-1^) (**<u>Fig. SI3.2</u>**<u>C</u>). Altogether, the combined propensity of protein and membrane interactions is conserved between the WT and ΔQCR6 models, but the number of productive interactions goes down in the latter, highlighting the importance of QCR6 for charge carrier binding and transport.

#### Cooperativity of QCR6 and Membrane electrostatics

Next, we focus on the role of anionic lipids in cyt. *c* recognition of the supercomplex. In the presence of anionic cardiolipins, the binding propensity of cyt. *c* per unit area of the membrane, ∼0.5 cyt.*c*/Å^2^, is only around 45% relative to that of the QCR6 binding propensity of 1.1 cyt.*c*/Å^2^ (**Fig. 5**D). So, for a given surface segment, cyt. *c* are much more localized around the supercomplex than being scattered over the membrane. Nonetheless, this relatively weaker cyt. *c*-membrane interaction still prevents the cyt. *c* pool from being lost into the bulk e.g. leaving only ∼10% of the cyt. *c* in the bulk (**Fig. 5**D<u>-inset</u>). When the anionic lipids are removed, despite a robust rate of cyt. *c* approaching the surface from the bulk (**<u>Fig. SI4.6</u>**), the bulk cyt. *c* concentration increases to 20-50% (**<u>Fig.</u> <u>SI4.7</u>**), dissipating the surface pool of carriers. Though the association of the remaining surface-bound cyt. *c* improves, 2.1 cyt.*c*/Å^2^ for the supercomplex and only 0.2 cyt.*c*/Å^2^ for the membrane (**<u>Fig. SI4.8</u>**) and have comparable residence times as in the anionic membrane (**<u>Fig. SI4.9</u>**), their occupancy at CIV sites is considerably reduced, as most cyt. *c* molecules are in the bulk (**Figs. 4** *vs.* **<u>SI4.3</u>**). Therefore, instead of competing with QCR6, the anionic cardiolipins complement the environment created by this hinge domain to enable cyt.*c*’s surface diffusion directed from CIII to CIV [47].

An artificial system is created, denoted *QCR6-trunc*, where the sequence of the QCR6 domain is partially truncated from Met1 to Glu46, representing a length comparable to the four higher organisms (see sequence alignment of **<u>Fig. SI1.1</u>**). Here also, similar to ΔQCR6 and QCR6, we find that the addition of anionic lipids to the membrane improves recognition of the supercomplex by cyt. *c* on the supercomplex, even with the remaining QCR6 (**<u>Fig. SI4.9</u>**). Our comparative analysis of WT, ΔQCR6 and QCR6-trunc BD simulations verify that fields from anionic lipids and those from the heterogeneous QCR6 conformations mutually cooperate to keep the cyt. *c* pool in the vicinity of the supercomplex during electron transport. Consequently, QCR6 acts as a recognition element within the supercomplex for the cyt. *c* to identify the mitochondrial complexes within the crowded cellular milieu. Despite not modifying the mechanism of binding, supercomplexes enrich the cyt. *c* pool and increases the probability of productive binding.

### QCR6 deletion potentially rigidifies supercomplex

QCR6 is not mandatory for supercomplex assembly, and yet our computational model highlighted its functional role for carrier protein recognition and diffusion [48]. So, the CIII-CIV composite is reassembled without the QCR6 to probe how it recovers from the loss of this recognition element. The resulting ΔQCR6 assembly was analyzed with cryo-EM, and a rigid-body refinement was performed with the relatively lower resolution dataset at 4.2 Å (see: Method: **<u>Cryo-EM data/details for ΔQCR6</u>**, **<u>Fig. SI5.1</u>**<u>A</u>). This model, obtained with around 81000 particles, shows a higher local resolution than that of the wild type CIII_2_CIV_2_ supercomplexes with data truncated at identical global resolution (**<u>Fig. SI5.1</u>**<u>B</u>). Interestingly, plants have a much smaller QCR6 that lacks the negatively charged N-terminal **<u>(Fig. SI1.1)</u>**. Only 28,000 particles are required to achieve a reconstruction at 3.2 Å resolution (PDB: 7JRP) [49], which is indicative of a reduced flexibility when QCR6 is small. These systematic differences in map resolution between QCR6-containing and deleted supercomplexes indicate the ΔQCR6 assembly to be less flexible than the wild type. A reason for this gained stability is that the removal of the whole QCR6 reduces charge-charge repulsion among subunits of the supercomplex at the p-side, particularly in the absence of cyt. *c*. Also, NADH-dehydrogenase or Complex I offers additional stability to the CIII_2_CIV_2_ supercomplex in higher organisms that are missing in the yeast, making the assembly more flexible. So, we find that the loss of recognition of the ΔQCR6 supercomplex is made up by a gain in stability, revealing the molecular origins of QCR6’s inaptness as a key ingredient for supercomplex assembly.

Normally, supercomplexes remain in an equilibrium between CIII_2_CIV_2_ and CIII_2_CIV stoichiometries, both containing similar CIII-CIV interfaces (**Fig. 6A-B**) [50]. The QCR6 deletion enables alternate packing distinct from the WT CIII_2_CIV/CIII_2_CIV_2_ linear to triangular conformations. The removal of QCR6 drastically reduced the surface electronegativity of the CIII_2_CIV supercomplex in the original wild-type arrangement of CIII and CIV (**Fig. 6**C). Rearrangement of the CIV into a triangular conformation, reducing the inter-complex distance from 73 Å to 66.0Å, assembles the ΔQCR6-CIII_2_CIV supercomplex into a more compact form (**Fig. 6 <u>D-E</u>**). This compact ΔQCR6 supercomplex regains the electrostatic potential over the extended CIII_2_CIV form. Hence, binding sites of the rigidified ΔQCR6-CIII_2_CIV supercomplex still remain viable for cyt. *c* binding, but less so relative to the QCR6-containing WT CIII_2_CIV_2_ supercomplex (**Fig. 6**). Supporting this computation, it is also biochemically shown that the ΔQCR6 mutants remain functionally active for transporting cyt. *c* limited by the levels of cytochrome *c*_1_ in CIII [51].

**Fig. 6:**
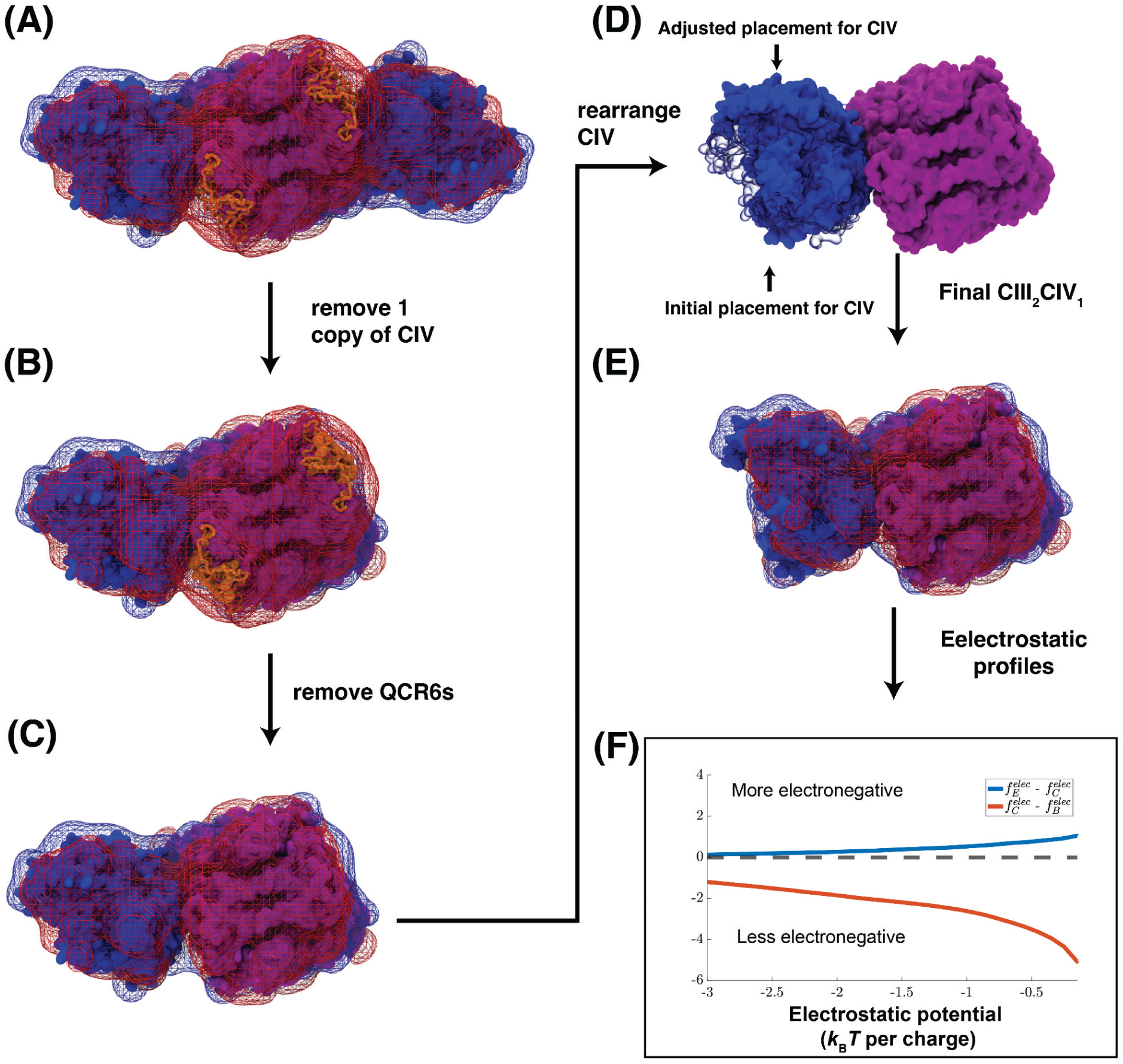
Electrostatic reorganization from CIII_2_CIV_2_ to CIII_2_CIV_1_ supercomplex upon QCR6 deletion. The electrostatic potential map of CIII_2_CIV_2_ and CIII_2_CIV_1_ complex in the WT and ΔQCR6 supercomplex, where electrostatic surfaces at a potential of 0.5*k_B_T* per charge and a potential of -0.5*k_B_T* per charge are given in blue and red, respectively. The impact of structural changes from CIII_2_CIV_2_ → CIII_2_CIV_1_ on the electrostatic properties is described stepwise **(A)** by first removing a single copy of CIV **(B)** then deleting QCR6 **(C)**, followed by adjustment on the remaining CIV’s position based on the cryo-EM information in **<u>Fig. SI2.6</u> (D** and **E)**. Visual inspection of the four iso-surfaces show negligible differences in the vicinity of CIII and CIV. The cumulative density function (CDF) for negative electrostatic potential values, denoted as *f^elec^*, was determined (see Methods: <u>Analyzes details</u>). CDFs for structures in panels (B), (C), and (E) are denoted as *f^elec^_B_*, *f^elec^_C_*, and *f^elec^_E_*, respectively. The pairwise difference between these distributions, presented in **(F)**, show that deletion of QCR6 expectedly reduces the strength of the electronegativity as from (B) to (C). The reorganization of the CIII_2_CIV structure (E) makes the assembly more electronegative than the CIV-deleted CIII_2_CIV_2_ structure in (C).

To quantify the role of QCR6 in supercomplex function, a rate-kinetic model is used. For a given stoichiometry of the donor and acceptor membrane proteins undergoing electron-exchange via carrier proteins, we have derived that the rate of ATP/sec generated from a given input pool of energy is inversely proportional to the turnover rate of the slower process [52] (See <u>Method: Estimating the ATP production</u> <u>rate</u>) - in the current case, the turnover rate of CIV. The rate matrix of **<u>Fig. SI3.2</u>** implicates this process as the timescale of CIII to CIV cyt. *c* transfer, which drops from 0.96 µs^-1^ in the native to 0.63 µs^-1^ in the ΔQCR6 supercomplex. Noting that respiration is ∼50% efficient and at any instant mitochondria generates 4 Watt of power (so it assimilates λ = 8 Watt), the 1.5-fold drop in CIV turnover translates to about 30% drop in the ATP production rate in ΔQCR6, which saturates to a drop of a maximum 40% even at unphysically energy intake [53] (**Fig. 7A**). Conversely, at very low energy intake values e.g. in aging cells, the native and ΔQCR6 supercomplex shows comparable efficiency, offering no general advantage to the surface guided charge transfer mechanism in the clustered environment. Recently, it has been argued that within crowded cellular environments the salinity is much lower (20mM) than the 150 mM determined from osmosis experiments [54]. This lower salinity is expected to reduce the screening effect and further increase the kinetic gap between the native and the ΔQCR6 constructs. The kinetic effect is expected to be dramatic if multiple cyt. *c* bind to the supercomplex (as we see in **Movie 3**) [36]. Indeed, the cyt. *c* activity dropped by ∼50% in ΔQCR6 [28], leading to petite phenotypes. Our simulations now reveal a potential molecular mechanism for this kinetic loss stemming from imprecise targeting of cyt. *c* within the supercomplex environment.

**Fig. 7:**
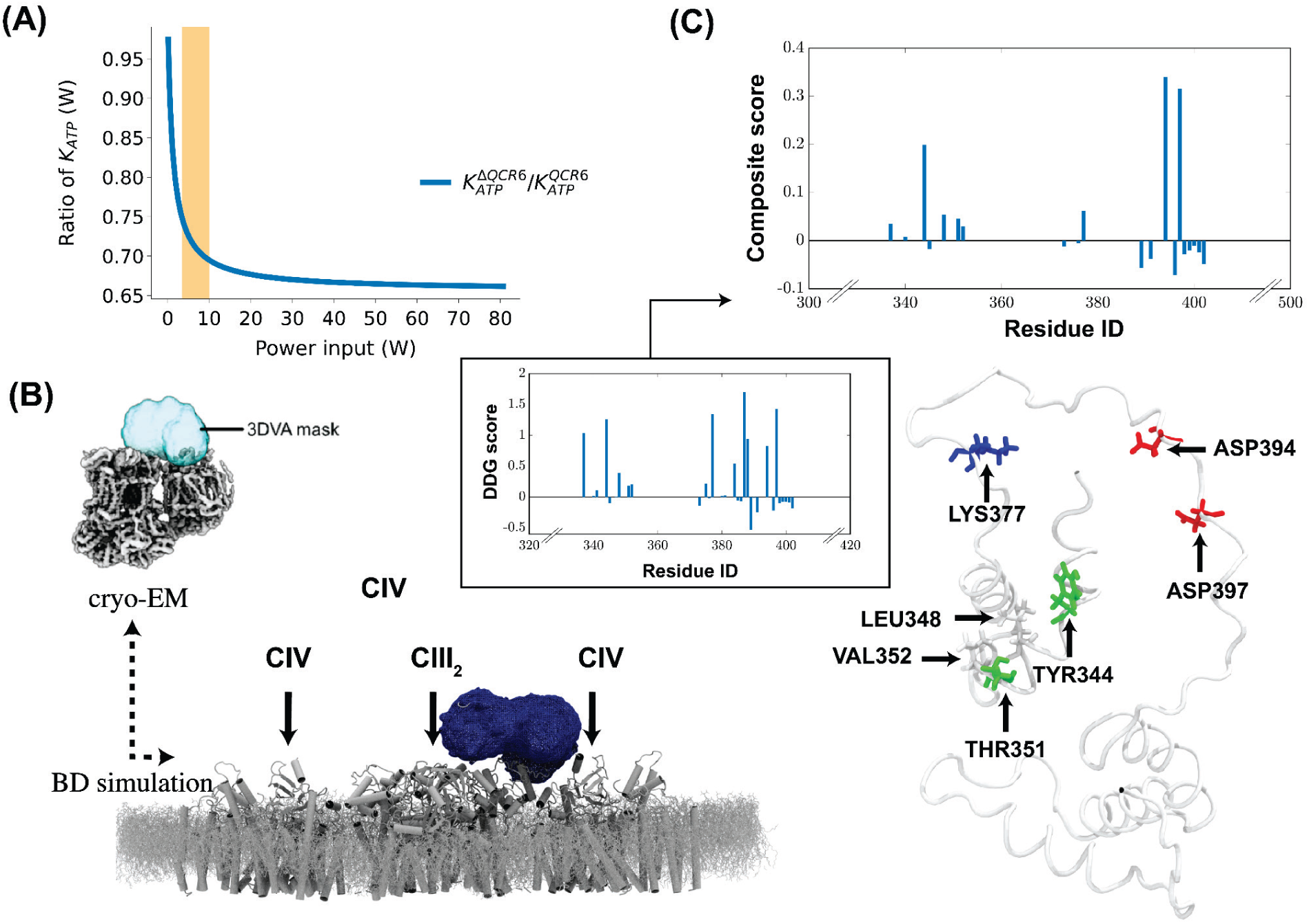
Diffusion profile of cyt. *c* and proposed mutations. (A) The rate of ATP production, measured in the number of ATP molecules generated per second, computed across a range of power input rate to the system, here yeast (See <u>Method: Estimating the ATP</u> <u>^p^roduction rat^e^</u>). The computed ATP production rate (*K_ATP_*) reveals a noticeable decline at the high power input domain upon the removal of the QCR6 hinge region. The range of input power (3.4W to 10W) for which a percentage drop of such removal starts to go beyond 25% to at least 30% is highlighted in orange. (B) Simulated density of cyt. *c* diffusing between CIII and CIV in BD simulations is shown as a contour surface of an occupancy map (isovalue = 0.5). For the same region, density of cyt. *c* observed by cryo-EM experiment and analyzed by 3-dimensional variational analysis (3DVA) is shown in inset - Figure adapted from [4] with permission from the authors. A cross-correlation analysis for this simulated diffusion profile of cyt. *c* and our experimentally obtained cryo-EM density map for SC is available in **Fig. <u>SI6.4</u>**. In (D), ΔΔG scores for individual residues of QCR6, which indicate the respective capability to stabilize QCR6 upon being mutated to ALA, are combined with respective residues’ percentage occupancies (**<u>Fig. SI6.1</u>**) to give a composite score for each residue. The top 6 residues with the highest composite scores are highlighted in the molecular image of QCR6, with acidic residues in red, basic residues in blue, polar residues are in green, and hydrophobic residues in white.

### A working model of cyt. *c* transport within the yeast CIII_2_CIV_2_ supercomplex

A general picture of diffusive electron transport within the supercomplex environment arises: first, following the accelerated MD results, the reduced CIII within the supercomplex remains in equilibrium between folded and unfolded QCR6 conformations. Second, according to the BD simulations, recognition and binding of an oxidized cyt. *c* to CIII populates its folded QCR6 state. Third, following from our past investigations and dissociation constant measurements, electron transfer from CIII into cyt. *c* weakens the interface. Fourth, it is inferred from the steered MD simulations that the reduced cyt. *c* is dislodged from CIII, and leveraging the electrostatic environment uniquely created by the anionic lipids and the extending QCR6 hinge within the supercomplex assembly, cyt. *c* is transported to the oxidized CIV. Finally, since residual interactions of the stretched QCR6 with the CIV-bound cyt. *c* is negligible to those of either CIII-bound or CIV-bound cyt. *c*, it is expected that the QCR6 hinge will disengage with cyt. *c* returning to its pre-equilibrium prior to the latter’s release from CIV into the medium, resetting the entire cycle (**Fig. 1**).

An occupancy map of cyt. *c* on the surface of the supercomplex (**Fig. 7**B), created by averaging over all cyt. *c*’s BD poses that make contact with either CIII or CIV (**Movie 3**), matches qualitatively with the mask of cyt. *c* movements created from three dimensional variational analysis of the cryo-EM density by Brizenski et al (**Fig. 7**B (inset)) [1]. This agreement between our simulated occupancy maps and the processed experimental data suggests that the grazing of cyt. *c* on the supercomplex seen in our model can be interpreted using a combination of surface diffusion as well as QCR6-induced conformational changes.

Finally, using Rosetta alanine scanning [55], we propose a list of mutations on the QCR6 region that will reduce its interaction with the cyt. *c* proteins, and decrease the conformationally coupled movements, relegating the charge transport primarily to diffusion. We primarily targeted the interface between the N-terminal hinge and cyt. *c* and chose residues with the top six most positive binding energies (ΔΔG) (**Figs. 7C** and **<u>SI6.1</u>**), followed by being weighted with their respective frequency of interacting with cyt. *c*. BD simulations of a hypothetical joint mutant brought to light the coupling between QCR6 conformations and diffusivity of the cyt. *c* proteins. The mutations collectively reduced the negative charge and weakened the attractive field in the vicinity of QCR6 (**<u>Fig. SI6.2</u>**<u>A</u>). Consequently, now, the diffusion of cyt. *c* leverage more of the bulk solution by spreading away from the supercomplex (**<u>Fig. SI6.2</u>**<u>C-D</u>) with less localization around the binding interface of CIII and CIV.

## Discussion

Capitalizing on cryo-EM structures, we report all-atom models and multiscale simulation of an entire respiratory supercomplex. The cumulative simulation time is 6 µs of MD and 14.08 ms (0.64 ms × 22) of BD simulations (**Table 1**). On one hand these computations push the envelope of modern exascale hardware such as on the Summit and Frontier supercomputers to achieve massively parallel biophysical computations, while on the other, it offers detailed insights on unresolved parts of the supercomplex and how they potentially contribute to membrane-wide electron transport. We found an electrostatic cooperativity between the negatively charged and flexible QCR6 loop of respiratory CIII and the anionic lipids in the membrane, which facilitates recognition and surface-bound transport of the cyt. *c* carrier proteins. Instead of competing for binding to cyt. *c*, the electrostatic potential from these two components reinforce each other to keep the electron carriers in the vicinity of the membrane. This finding is reinforced by the fact that about 8-fold higher concentrations of complex IV from cardiolipin-free mitochondria had to be used to obtain rates comparable with those in wild-type mitochondria [56].

Unfolding or folding dynamics of the QCR6, specifically in yeast, prompts charge transport towards or away from CIII, supporting a directional CIII → CIV electron transfer model. Removal of either one of these recognition elements from the membrane model makes the transport of cyt. *c* more three-dimensional, compromises the number or residence time of productive binding to the electron donors or acceptor complexes, or reorganizes the supercomplex to recover for the loss in surface potential. Electron transfer from CIII to CIV by two-dimensional diffusion of cyt. *c* along the supercomplex surface resembles a “substrate channeling” model, which has been criticized based on the finding of that cyt. *c* diffusion in *S. cerevisiae* is unrestricted [57]. However, our QCR6-guided model of surface diffusion of cyt. *c* is not in conflict with this finding because it elucidates electrostatic interactions between cyt. *c* and the supercomplex surface, and cyt. *c* remains in equilibrium with the cyt. *c* pool during the entire electron-transfer process. So, when the QCR6 is removed, the ATP activity drops, but still saturates to 60-70% of the original; the bulk cyt. *c*, though slow, still shuttles the electron to enable this reduced activity.

In primitive organisms such as the heliobacteria, the cyt. *c* remain tethered to the membrane by employing covalent linkers of sequence presented in **<u>Fig. SI6.3</u>**. These linkers diffuse with the membrane, and keep the charge carriers in the vicinity of the membrane. Consequently, the need to create a locally negative charge environment is partially avoided for keeping the cyt. *c* confined. Supercomplexes are yet to be observed in these species. The N-terminal domain of the mitochondrial QCR6 unit has a highly dissimilar sequence with the covalent tethers. This suggests that confinement of the charge carrier on the membrane surface is ubiquitous within multiple organisms, but the mechanism has evolved. In the absence of QCR6, covalent linkages are required to confine charge carriers. QCR6 improves the probability of cyt. *c* recognition either in isolated CIII or as part of the supercomplex. It potentially transports cyt. *c* by directly interacting with the membrane, as can be seen in the yeast membrane. When it is removed, the supercomplex reorganizes to still mainitin an optimal environment for confining the carriers that can still exchange between the bulk and the surface. In our studies, the electrostatic confinement of charge carriers is relevant to yeast supercomplexes that preclude Complex I, and similarly for purple bacteria where electron transfer occurs between light-harvesting reaction center complexes and CIII [52]. Plants and mammals possess much shorter QCR6 domains, but it has been surmised that presence of Complex I together with anionic lipids stabilizes the supercomplex assembly [49,58], which still creates a local electrostatic environment to keep cyt. *c* in the vicinity of the donor and acceptor integral-membrane complexes. Hence, the need for a longer yeast-like QCR6 is avoided in higher eukaryotes.

So, QCR6, despite not being an essential component for supercomplex stability, remains a key element of CIII composition, recognition and function, that is possibly over-utilized in yeast as our simulations exhibit. Any modification of QCR6 compromise is expected to impact the catalytic activity of CIII, which is indeed the case. Also, if QCR6 is removed, supercomplex formation is a tangible way of receiving cyt. *c*_1_ and hence CIII activity. We have now simulated the mechanism of this recovery of activity by describing the reorganization of the electrostatic environment, reinforcing that confinement of the charge carrier is an evolutionary requirement. Taken together, there can be three mechanisms to ensure confinement of charge carriers: covalent linkers, isolated complexes or supercomplexes. QCR6 on one hand, acts as a linker for guided surface diffusion when participating in the yeast supercomplex without driving complex formation, while on the other hand, still maintains its functional properties seen in CIII acting as a recognition element.

## Methods

The entire modeling workflow is summarized in Fig. SI7.1.

### Initial model construction

Though the flexible domain of QCR6 has thus far been unable to be resolved using experimental methods, the sequence is known (Fig. SI1.1). Five predicted structures for the flexible domain of QCR6 were generated through QUARK [60]. Using VMD, these structures were aligned with the resolved region of QCR6 in a Complex III with reduced cytochrome *c* bound (PDB: 3CX5). However, when aligning the QCR6, there were steric clashes between the flexible domain and Complex III that were unable to be resolved through conventional molecular dynamics. So, after some minimization of the model, the following MELD simulations were performed.

### MELD

The Modeling Employing Limited Data (MELD) approach is useful to integrate sources of ambiguous and noisy information about the molecular system. In particular, we were interested in modeling QCR6, which is a subunit of complex III (CIII), and its interaction with the remaining CIII in the presence of a CIII-bound cytochrome *c* (cyt.*c*).

QCR6 is a small subunit of 147 residues. Among these residues, residues 74 to 147 have a well defined structure and are experimentally resolved. In contrast, residues 1 to 73 constitute a highly acidic region that is highly flexible and remains unsolved in both the crystal and cryo-EM density map. This flexible linker region is most likely unstructured or intrinsically disordered and is thus well posed for MELD, which is capable of modeling such flexibility while incorporating information from the resolved region. It should be noted that this flexible linker region, though interacting with CIII domains embedded in the membrane, is found outside the membrane.

For computational efficiency, we first constructed an initial model with CIII (PDB: 8E7S) domains (without QCR6) that were within 8Å of QCR6 (chain Q). In this model, we excluded other proteins in the CIII_2_CIV_2_ supercomplex with which QCR6 might interact (chain F or W). We capped incomplete subunits (chains) of our initial model with ACE and NME residues. Then, we generated preliminary QCR6 models by QUARK. As all QUARK models had residues 1 to 73 of QCR6 overlapping with our starting model, which contained CIII, we removed residues 1 to 73 from these quark models and modeled residues 1 to 73 as a non-overlapping, extended chain outside our starting model. Lastly, we attached this extended chain to our initial model with the resolved part of QCR6 (residues 74 to 147) at its C-termini. This model with an extended chain of QCR6 formed the final form of our initial model.

We next modeled the information to guide the simulations. We chose two lists of residues from our initial model. List 1 consisted of residues {236 318 320 237 331 330 304 481 184 223 220 286 287} from cyt. *c* and list 2 consisted of all residues in the flexible region of QCR6 (residues 1 to 73) (**<u>Fig. SI7.2</u>**). We combinatorially created all possible contacts between the two lists (collection). Within the collection, we grouped restraints such that each CIII residue (without QCR6) would have a contact (pairwise alpha carbon distances less than 8Å) to any residue of the flexible region. We satisfied one restraint in each group, and overall we satisfied 60% of the 13 groups in the collection. We further used cartesian restrains on residues belonging to CIII. Other aspects of the simulation were as in previous MELD simulations [31], using the ff14SB side force field, a 4.5fs timestep, and with the H,T-REMD protocol. The H,T-REMD protocol used 24 550 ns long replicas that expanded a temperature range of 300K to 450K with the data (QCR6-CIII contacts) enforced strongly at low replica index and with force constants vanished at the highest replica index.

### MD and Steered MD (SMD)

From the MELD results, interaction energies between cyt.*c* and Complex III were calculated using the NAMD Energy Plugin in VMD. Four conformational states were identified, with the first being the unfolded flexible domain of QCR6 (state 0 in **Fig. 2**), the last fully folded (state 3), and two intermediate states (states 1 and 2). A model from state 3, which had the lowest interaction energy was used to manually incorporate into the full CIII_2_CIV_2_ supercomplex. This model was set up in two membranes in CHARMM-GUI [61]: one with just POPC as a reference, and one with POPC + Cardiolipin (CL) to mimic the supercomplex environment. In particular, the POPG lipids binding to the supercomplex in the original 8E7S model were replaced with CL. The system was parameterized with CHARMM36 force fields and CHARMM-GUI was also used to solvate and ionize both systems. The system was thermalized under NPT conditions and explicit solvent molecular dynamics simulations were done using NAMD 2.14 (double check what version was on OLCF’s Summit) for 1 µs [62].

After explicit solvent molecular dynamics, SMD was used to probe the dynamics of QCR6 as a hinge when cyt. *c* moves from CIII to CIV. A distance collective variable was used, where one group was defined as the center of mass of alpha carbons of cyt. *c* and the second group defined five residues around the CIV binding pocket (TYR146, ASP150, GLU151, ASP162, ASP164) and the force constant used was 0.05 kcal/mol. Though no specific residues were identified in previous studies, a predicted binding site on CIV was used [28], where similar residues were used as targets for the binding site (find residues used). Following SMD, interaction energies were calculated between cyt. *c*-CIII and cytochrome *c*-CIV, using the NAMD Energy plugin in VMD. Dissociation constants (K_d_) were calculated using the PRODIGY web-server [63].

### Brownian Dynamics (BD)

BD simulations had been used to sample cyt. *c*’s diffusion over the yeast CIII_2_CIV_2_ supercomplex through an in-house software, the Atomic-Resolution Brownian Dynamics (ARBD) [64]. The starting models are extracted at times 0 ns, 220 ns, 400 ns, and 480 ns from the SMD simulations, which provided local energy minima of the cyt. *c*-QCR6 interaction energy profile of **Fig. 4**. The underlying mechanism for and the setup of BD simulations using ARBD had been discussed in detail previously [47,65,66]. Here, following this previous approach, we will outline our setup of BD simulations below.

First, we classified our biomolecules into 2 categories, namely the diffusing particle and the diffusion environment. In this project, cyt. *c* is our diffusing particle and it was dynamic during BD simulations. Its atoms were clustered into 4 representative atom types as in [47]. The particle distribution for each of these representative atom types and the charge distribution for the whole cyt. *c* were determined by VMD [67]. Our cyt. *c* was then represented by these distributions, each stored as a volumetric data grid. On the other hand, each of the different membrane systems, with the supercomplex embedded, was our diffusion environment. It was static during BD simulations, and its electrostatic potential and its potential of mean force (PMF) for each of the representative atom types of cyt. *c* were computed by Adaptive Poisson-Boltzman Solver [68,69] and Implicit ligand sampling [70], respectively. Then, to mimic the fluctuating nature of atoms’ positions, a 3D-Gaussian filter with a half width of 1.5Å, which resembles the RMSF of atoms belonging to the protein-membrane interface, was convoluted with each of these potentials. Finally, during BD simulations of each membrane system, the different volumetric data grids for cyt. *c* interacted with either the electrostatic potential or the corresponding PMFs of the membrane system. This interaction was converted to forces on cyt. *c*, as being prescribed in [47]

After the volumetric data grids for cyt. *c* as well as the electrostatic potentials and PMFs for each membrane were computed, BD simulations were launched. For each replica of BD simulation, the membrane midplane of the membrane system was located at *z* = 0, and one cyt. *c* was initially positioned above the supercomplex at a height of 100Å, which corresponded to *z* ∼ 150Å. The *x* and *y* coordinates for our cyt. *c* were randomized.

For each membrane system, there were 320 replicas of 2-microseconds long BD simulations. In the current study, we employed 21 different membrane systems, each differed from another in terms of either the lipid composition of the membrane or the conformation of QCR6 or both. The total simulation time for BD simulations in the current study was 1.12 milliseconds.

### Gaussian accelerated Molecular Dynamics (GaMD)

GaMD simulations [71] were performed at 3 time points for QCR6 with and without cytochrome *c*. All GaMD simulations were performed using AMBER 20. The proteins were parameterized using AMBER ff14SB force field, solvated in a TIP3P water box of dimension 20 Å, and K+ ions were added for neutralization. This system was minimized, heated to 300K under NVT conditions, and equilibrated at 300K under NPT conditions for 2 ns. Following this system preparation, GaMD equilibration was performed for 20 ns, including 2 ns of conventional MD. GaMD production was performed for 500 ns.

### Non-uniform refinement of WT supercomplex

As described in [25], 493,055 particle images of the wildtype supercomplex were reconstructed using standard single particle cryo-EM image processing routines in CryoSparc (v3.2) resulting in a final density map of the supercomplex at 3.2Å resolution [72]. Using this same dataset, 3D variability analysis was carried out to identify heterogeneity in this data set. Of the 10 modes identified in the 3D variability analysis, the five modes containing the largest number of particles, a total of 85,455 particles, were selected for a heterogenous refinement and reconstruction. This refinement generated 2 C2-symmetry reconstructions: class 0, containing 19,447 particles, reconstructed to 9.3Å and class 1, containing 66,008 particles, reconstructed to 6.6Å resolution. Particles in class 1 were further processed imposing C1 symmetry and using non-uniform refinement, which were then reconstructed to 4.3Å resolution. The wildtype model from Hryc et al (PDB: 8E7S) was then fit to this 4.3Å resolution density map using Chimera (v1.16), from which unmodeled density could be observed near the termini of QCR6.

### ΔQCR6 strain construction and purification

Mitochondria isolation and Supercomplex purification from ΔQCR6 strain were performed as described previously for the wild type strain [25].

#### Mitochondria isolation

Cells were harvested at OD_600_2-3 by centrifugation (4700 x g for 10 min at 4 °C), washed with ice-cold 1X Tris-buffered saline. The pellet was resuspended in 100 mM TrisSO4 (pH 9.4) and 10mM dithiothreitol (2 ml/g of cell pellet), incubated at 30 °C for 20 min with shaking, centrifuged at 750 x g at 4 °C for 10 min, washed with 1 M sorbitol. The cell wall was digested by zymolase-20T at 3mg/g weight of cells in 1 M sorbitol and 20mM Potassium phosphate buffer (pH 7.5) for 90 min at 37 °C. The spheroplasts were washed with 1M Sorbitol (2ml/g of cells) at 4 °C and disrupted with a Dounce homogenizer in the 10 mM Tris-HCl buffer (pH 7.2) with 0.5M sorbitol, 0.02% bovine serum albumin, 1/1000 volume of protease inhibitor cocktail set III (Calbiochem Millipore) and 1mM phenylmethylsulfonyl fluoride. The lysed cells were centrifuged at 1700 x g at 4 °C for 5 min, and the supernatant was centrifuged at 8500 x g at 4 °C for 20 min. The pellet was suspended in 1ml of 10mM Mops (pH 7.2) buffer with 250 mM sucrose and 1mM EDTA and loaded onto a 15–60% sucrose gradient in 15 mM Tris-HCl (pH 7.4) and 20mM KCl that was centrifuged at 111,000 x g at 4 °C for 90min. The mitochondrial layer was aspirated, flash frozen with liquid nitrogen and stored at −80 °C.

#### Supercomplex purification

The Supercomplex from ΔQCR6 strain was purified by the method we previously developed for WT [25]. Isolated mitochondria (12 mg of protein) were suspended in the 1.5 ml of lysis buffer containing 2% (w/v) digitonin, 50mM potassium acetate, 10% glycerol, 1:50 volume protease inhibitor cocktail, 1.5 mM phenylmethylsulfonyl fluoride and 30mM HEPES-KOH, (pH 7.4) for 1h at 4 °C with gentle shaking. After incubation the lysate was centrifuged at 4 °C for 20 min at 90,700 x g. Supernatant was incubated with magnetized Cobalt Beads (Dynabeads TALON catalog #101.02D, Invitrogen) for 45 min at 4 °C with shaking to remove FoF1-ATPase. After incubation the beads were removed using a magnetic separator. The supernatant (1.5 ml) was layered onto two equal 8 ml sucrose gradients (0.75M to 1.5M sucrose in 15mM Tris-HCl (pH 7.2), 20mM KCl, and 0.4% digitonin) and centrifuged at 4 °C for 22 hours at 111,000 x g. Fractions (80–100 μL) from the gradients were analyzed by BN-PAGE and western Blot with antibodies to yeast CIII and CIV as described previously [25]. Fractions containing the purified SC were combined and used for cryo-EM. Protein concentrations were determined using the BCA protein assay kit (Thermo Scientific) according to manufacturer’s instructions.

### Cryo-EM data/details for ΔQCR6

Data was collected with a Titan Krios microscope (Thermo Fisher Scientific) operated at 300kV. A post-GIF K2 Summit direct electron detector (Gatan), operating in counting mode was used at a nominal magnification of 130,000× (pixel size of 1.07Å) for image collection, and an energy slit with a width of 20eV was used during data collection. A total dose of 56e^-^/Å^2^ fractionated over 35 frames; nominal defocus range set from −1.5μm to −4μm. Here, 7,750 micrograph movies were collected of ΔQCR6.

The ΔQCR6 complex was processed with CryoSPARC [72]. Patch motion was used for frame alignment and exposure weighting with default parameters. Patch CTF was used to generate CTF parameters. Blob picker was initially used on a subset of images (200) to select a subset of particles, which were then used to generate a low-resolution template. This template was then used to train the Topaz particle picker [73]. A total of 369,297 particles were selected. Multiple rounds of 2D classification then followed, narrowing the dataset to 81,260 particles. An initial refinement was completed using C1 symmetry, which resulted in a 7.77Å density map. A local refinement routine focusing on a CIII dimer and a CIV monomer then followed (using a mask to reduce the detergent band). This resulted in a 4.21Å structure.

## Acknowledgements

We acknowledge a CAREER award from NSF (MCB-1942763). This work is also supported by the National Defense Education Program (NDEP) program under grant number HQ0034-21-S-F001, and DOE grants DE-SC0022956 and DE-SC0010575. The simulation work used resources of the Oak Ridge Leadership Computing Facility’s (OLCF) Frontier Supercomputer, which was awarded through the ASCR Leadership Computing Challenge (ALCC). OLCF is a DOE Office of Science User Facility supported under Contract DE-AC05-00OR22725. We further acknowledge John Rubinstein and Peter Brzezinski for permitting us to use their 3DVA analyzed density data in our analysis.

## Notes

### Competing Interest Statement

The authors have declared no competing interest.

## References

1. Brzezinski, P., Moe, A. and Ädelroth, P. (2021) ‘Structure and mechanism of respiratory III–IV supercomplexes in bioenergetic membranes’, Chemical Reviews, 121(15), pp. 9644–9673. doi:10.1021/acs.chemrev.1c00140.

2. Martin, D.R. and Matyushov, D.V. (2017) ‘Electron-transfer chain in respiratory complex I’, Scientific Reports, 7(1). doi:10.1038/s41598-017-05779-y.

3. Moe, A., Trani, J.D., Rubinstein, J.L.. (2021) ‘Cryo-EM structure and kinetics reveal electron transfer by 2D diffusion of cytochrome c in the yeast III-IV respiratory supercomplex’, Proceedings of the National Academy of Sciences, 118(11). doi:10.1073/pnas.2021157118.

4. Jha P, Wang X, Auwerx J. Analysis of Mitochondrial Respiratory Chain Supercomplexes Using Blue Native Polyacrylamide Gel Electrophoresis (BN-PAGE). Curr Protoc Mouse Biol. 2016 Mar 1;6(1):1–14. doi: 10.1002/9780470942390.mo150182.

5. Schagger, H. (2000) ‘Supercomplexes in the respiratory chains of yeast and mammalian mitochondria’, The EMBO Journal, 19(8), pp. 1777–1783. doi:10.1093/emboj/19.8.1777.

6. Mileykovskaya, E.; Penczek, P. A.; Fang, J.; Mallampalli, V. K. P. S.; Sparagna, G. C.; Dowhan, W. Arrangement of the Respiratory Chain Complexes in Saccharomyces Cerevisiae Supercomplex III2IV2 Revealed by Single Particle Cryo-Electron Microscopy. Journal of Biological Chemistry 2012, 287 (27), 23095–23103. DOI:10.1074/jbc.m112.367888.

7. Maranzana, E.; Barbero, G.; Falasca, A. I.; Lenaz, G.; Genova, M. L. Mitochondrial Respiratory Supercomplex Association Limits Production of Reactive Oxygen Species from Complex I. Antioxidants & Redox Signaling 2013, 19 (13), 1469–1480. DOI:10.1089/ars.2012.4845.

8. Rydström Lundin C.; Von Ballmoos C.; Ott M.; Ädelroth P.; Brzezinski P. Regulatory role of the respiratory supercomplex factors in Saccharomyces cerevisiae. Proc. Natl. Acad. Sci. U. S. A. 2016, 113, E4476–E4485. doi:10.1073/pnas.1601196113.

9. Javadov, S.; Jang, S.; Chapa-Dubocq, X. R.; Khuchua, Z.; Camara, A. K. Mitochondrial Respiratory Supercomplexes in Mammalian Cells: Structural versus Functional Role. Journal of Molecular Medicine 2020, 99 (1), 57–73. DOI:10.1007/s00109-020-02004-8.

10. Novack, G. V.; Galeano, P.; Castaño, E. M.; Morelli, L. Mitochondrial Supercomplexes: Physiological Organization and Dysregulation in Age-Related Neurodegenerative Disorders. Frontiers in Endocrinology 2020, 11. DOI:10.3389/fendo.2020.00600.

11. Blaza J. N.; Serreli R.; Jones A. J. Y.; Mohammed K.; Hirst J. Kinetic evidence against partitioning of the ubiquinone pool and the catalytic relevance of respiratory-chain supercomplexes. Proc. Natl. Acad. Sci. U. S. A. 2014, 111, 15735–15740. 10.1073/pnas.1413855111.

12. Berndtsson, J.; Kohler, A.; Rathore, S.; Marin-Buera, L.; Dawitz, H.; Diessl, J.; Kohler, V.; Barrientos, A.; Büttner, S.; Fontanesi, F.; Ott, M. Respiratory Supercomplexes Enhance Electron Transport by Decreasing Cytochrome *c* Diffusion Distance. EMBO reports 2020, 21 (12). DOI:10.15252/embr.202051015.

13. Kao, W.-C.; Ortmann de Percin Northumberland, C.; Cheng, T. C.; Ortiz, J.; Durand, A.; von Loeffelholz, O.; Schilling, O.; Biniossek, M. L.; Klaholz, B. P.; Hunte, C. Structural Basis for Safe and Efficient Energy Conversion in a Respiratory Supercomplex. Nature Communications 2022, 13 (1). DOI:10.1038/s41467-022-28179-x.

14. Wikström, M.; Springett, R. Thermodynamic Efficiency, Reversibility, and Degree of Coupling in Energy Conservation by the Mitochondrial Respiratory Chain. Communications Biology 2020, 3 (1). DOI:10.1038/s42003-020-01192-w.

15. Genova, M. L.; Lenaz, G. Functional Role of Mitochondrial Respiratory Supercomplexes. Biochimica et Biophysica Acta (BBA) - Bioenergetics 2014, 1837 (4), 427–443. DOI:10.1016/j.bbabio.2013.11.002.

16. Hackenbrock, C. R.; Chazotte, B.; Gupte, S. S. The Random Collision Model and a Critical Assessment of Diffusion and Collision in Mitochondrial Electron Transport. Journal of Bioenergetics and Biomembranes 1986, 18 (5), 331–368. DOI:10.1007/bf00743010.

17. Pfeiffer, K.; Gohil, V.; Stuart, R. A.; Hunte, C.; Brandt, U.; Greenberg, M. L.; Schägger, H. Cardiolipin Stabilizes Respiratory Chain Supercomplexes. Journal of Biological Chemistry 2003, 278 (52), 52873–52880. DOI:10.1074/jbc.m308366200.

18. Stuchebrukhov, A., Schäfer, J., Berg, J., & Brzezinski, P. (2020). Kinetic advantage of forming respiratory supercomplexes. Biochimica et biophysica acta. Bioenergetics, 1861(7), 148193. 10.1016/j.bbabio.2020.148193

19. Pérez-Mejías, G. Guerra-Castellano, A., Díaz-Quintana, A., De la Rosa, M.A., Díaz-Moreno, I.. (2019) ‘Cytochrome c: Surfing off of the mitochondrial membrane on the tops of complexes III and IV’, Computational and Structural Biotechnology Journal, 17, pp. 654– 660. doi:10.1016/j.csbj.2019.05.002.

20. Bergstrom, C., Beales, P.A., Lv, Y., Vanderlick, K.T., Groves, J.T. Cytochrome c causes pore formation in cardiolipin-containing membranes. Biochemistry 2012, 110 (16) 6269–6274 doi:10.1073/pnas.1303819110

21. Böttinger, L.; Horvath, S. E.; Kleinschroth, T.; Hunte, C.; Daum, G.; Pfanner, N.; Becker, T. Phosphatidylethanolamine and Cardiolipin Differentially Affect the Stability of Mitochondrial Respiratory Chain Supercomplexes. Journal of Molecular Biology 2012, 423 (5), 677–686. DOI:10.1016/j.jmb.2012.09.001.

22. Mileykovskaya, E.; Dowhan, W. Cardiolipin-Dependent Formation of Mitochondrial Respiratory Supercomplexes. Chemistry and Physics of Lipids 2014, 179, 42–48. DOI:10.1016/j.chemphyslip.2013.10.012.

23. Wú, F., Mühleip, A., Gruhl, T., Sheiner, L., Maréchal, A., Amunts, A. (2024). Structure of the II2-III2-IV2 mitochondrial supercomplex from the parasite *Perkinsus marinus*. bioRxiv. doi: 10.1101/2024.05.25.595893

24. Chicco, A. J.; Sparagna, G. C. Role of Cardiolipin Alterations in Mitochondrial Dysfunction and Disease. American Journal of Physiology-Cell Physiology 2007, 292 (1). DOI:10.1152/ajpcell.00243.2006.

25. Hryc, C.F. Mallampalli, V.K.P.S., Bovshik, E.I., Azinas, S., Fan, G., Serysheva, I.I., Sparagna, G.C., Baker, M.L., Mileykovskaya, E., Dowhan, W. (2023) ‘Structural insights into cardiolipin replacement by phosphatidylglycerol in a cardiolipin-lacking yeast respiratory supercomplex’, Nature Communications, 14(1). doi:10.1038/s41467-023-38441-5.

26. Hartley, A. M., Lukoyanova, N., Zhang, Y., Cabrera-Orefice, A., Arnold, S., Meunier, B., Pinotsis, N., & Maréchal, A. (2019). Structure of yeast cytochrome c oxidase in a supercomplex with cytochrome bc1. Nature structural & molecular biology, 26(1), 78–83. 10.1038/s41594-018-0172-z

27. Gómez, L. A.; Hagen, T. M. Age-Related Decline in Mitochondrial Bioenergetics: Does Supercomplex Destabilization Determine Lower Oxidative Capacity and Higher Superoxide Production? Seminars in Cell & Developmental Biology 2012, 23 (7), 758– 767. DOI:10.1016/j.semcdb.2012.04.002.

28. Yang, M.; Trumpower, B. L. Deletion of QCR6, the Gene Encoding Subunit Six of the Mitochondrial Cytochrome Bc1 Complex, Blocks Maturation of Cytochrome C1, and Causes Temperature-Sensitive Petite Growth in Saccharomyces Cerevisiae. Journal of Biological Chemistry 1994, 269 (2), 1270–1275. DOI:10.1016/s0021-9258(17)42253-8.

29. Nakai, M.; Endo, T.; Hase, T.; Tanaka, Y.; Trumpower, B. L.; Ishiwatari, H.; Asada, A.; Bogaki, M.; Matsubara, H. Acidic Regions of Cytochrome C1 Are Essential for Ubiquinol-Cytochrome c Reductase Activity in Yeast Cells Lacking the Acidic Qcr6 Protein1. The Journal of Biochemistry 1993, 114 (6), 919–925. DOI:10.1093/oxfordjournals.jbchem.a124277.

30. Modjtahedi, N.; Tokatlidis, K.; Dessen, P.; Kroemer, G. Mitochondrial Proteins Containing Coiled-Coil-Helix-Coiled-Coil-Helix (CHCH) Domains in Health and Disease. Trends in Biochemical Sciences 2016, 41 (3), 245–260. DOI:10.1016/j.tibs.2015.12.004.

31. MacCallum, J.L., Perez, A. and Dill, K.A. (2015) ‘Determining protein structures by combining semireliable data with atomistic physical models by Bayesian inference’, Proceedings of the National Academy of Sciences, 112(22), pp. 6985–6990. doi:10.1073/pnas.1506788112.

32. Shekhar, M., Terashi, G., Gupta, C., Sarkar, D., Debussche, G., Sisco, N.J., Nguyen, J., Mondal, A., Vant, J., Fromme, P., Van Horn, W.D., Tajkhorshid, E., Kihara, D., Dill, K., Perez, A., Singharoy, A. (2021), CryoFold: Determining protein structures and data-guided ensembles from cryo-EM density maps. Matter, 4 (10), 3195–3216. doi:10.1016/j.matt.2021.09.004.

33. C Maffeo, B Luan, A Aksimentiev, End-to-end attraction of duplex DNA. Nucleic Acids Res 40, 3812–3821 (2012).

34. Hunte, C. Koepke, J., Lange, C., Roßmanith, T., Michel, Hartmut. (2000) ‘Structure at 2.3 Å resolution of the cytochrome bc1 complex from the yeast saccharomyces cerevisiae co-crystallized with an antibody FV fragment’, Structure, 8(6), pp. 669–684. doi:10.1016/s0969-2126(00)00152-0.

35. Solmaz, S.R.N. and Hunte, C. (2008) ‘Structure of complex III with bound cytochrome c in reduced state and definition of a minimal core interface for electron transfer’, Journal of Biological Chemistry, 283(25), pp. 17542–17549. doi:10.1074/jbc.m710126200.

36. Lobez, A.P., Wu, F., Di Trani, J.M., Rubinstein, J.L., Oliveberg, M., Brzezinski, P., Moe, A.. Electron transfer in the respiratory chain at low salinity. Nat Commun 15, 8241 (2024). 10.1038/s41467-024-52475-3

37. Devanathan S, Salamon Z, Tollin G, Fitch JC, Meyer TE, Berry EA, Cusanovich MA. Plasmon waveguide resonance spectroscopic evidence for differential binding of oxidized and reduced Rhodobacter capsulatus cytochrome c2 to the cytochrome bc1 complex mediated by the conformation of the Rieske iron-sulfur protein. Biochemistry. 2007 Jun 19;46(24):7138–45. doi: 10.1021/bi602649u.

38. Singharoy A, Barragan AM, Thangapandian S, Tajkhorshid E, Schulten K. Binding Site Recognition and Docking Dynamics of a Single Electron Transport Protein: Cytochrome c2. J Am Chem Soc. 2016 Sep 21;138(37):12077–89. doi: 10.1021/jacs.6b01193.

39. Shimada, S. Shinzawa-Itoh, K., Baba, J., Aoe, S., Shimada, A., Yamashita, E., Kang, Jiyoung., Tateno, M., Yoshikawa, S., Tsukihara, T.. (2016) ‘Complex structure of cytochrome *c*–cytochrome *c* oxidase reveals a novel protein–protein interaction mode’, The EMBO Journal, 36(3), pp. 291–300. doi:10.15252/embj.201695021.

40. Wilms, J., Veerman, E.C.I., König, B.W., Dekker, H.L., van Gelder, B.F. (1981) ‘Ionic strength effects on cytochrome aa3 kinetics’, Biochimica et Biophysica Acta (BBA) - Bioenergetics, 635(1), pp. 13–24. doi:10.1016/0005-2728(81)90003-7.

41. Ferguson-Miller, S., Brautigan, D.L. and Margoliash, E. (1976) ‘Correlation of the kinetics of electron transfer activity of various eukaryotic cytochromes C with binding to mitochondrial cytochrome c oxidase.’, Journal of Biological Chemistry, 251(4), pp. 1104– 1115. doi:10.1016/s0021-9258(17)33807-3.

42. Malatesta, F., Antonini, G., Nicoletti, F., Giuffrè, A., D’Itri, E., Sarti, P., Brunori, M. (1996) ‘Probing the high-affinity site of beef heart cytochrome *c* oxidase by cross-linking’, Biochemical Journal, 315(3), pp. 909–916. doi:10.1042/bj3150909.

43. Lobo-Jarne, T. and Ugalde, C. (2018) ‘Respiratory chain supercomplexes: Structures, function and biogenesis’, Seminars in Cell & Developmental Biology, 76, pp. 179–190. doi:10.1016/j.semcdb.2017.07.021.

44. Heinemeyer, J., Braun, H-P., Boekema, E.J., Kouril, R.. (2007) ‘A structural model of the cytochrome c reductase/oxidase Supercomplex from yeast mitochondria’, Journal of Biological Chemistry, 282(16), pp. 12240–12248. doi:10.1074/jbc.m610545200.

45. Abresch, E.C. Gong, X-M., Paddock, M.L., Okamura, M.Y.. (2009) ‘Electron transfer from cytochrome *c*2 to the reaction center: A transition state model for ionic strength effects due to neutral mutations’, Biochemistry, 48(48), pp. 11390–11398. doi:10.1021/bi901332t.

46. Zheng, W., Chai, P., Zhu, J. et al. High-resolution in situ structures of mammalian respiratory supercomplexes. Nature 631, 232–239 (2024). doi:10.1038/s41586-024-07488-9.

47. C. K. Chan, A. Singharoy, and E. Tajkhorshid, “Anionic Lipids Confine Cytochrome *c*2 to the Surface of Bioenergetic Membranes without Compromising Its Interaction with Redox Partners.” Biochemistry, 2022 Mar 1;61(5):385–397.

48. M. Yang, B.L. Trumpower (1994) Deletion of QCR6, the gene encoding subunit six of the mitochondrial cytochrome bc1 complex, blocks maturation of cytochrome c1, and causes temperature-sensitive petite growth in Saccharomyces cerevisiae. Journal of Biological Chemistry.

49. Maria Maldonado, Fei Guo, James A Letts (2021) Atomic structures of respiratory complex III2, complex IV, and supercomplex III2-IV from vascular plants. eLife 2021 10.7554/eLife.62047

50. Vercellino, I. and Sazanov, L.A. (2021) ‘Structure and assembly of the mammalian mitochondrial supercomplex CIII2CIV’, Nature, 598(7880), pp. 364–367. doi:10.1038/s41586-021-03927-z.

51. Smith PM, Fox JL, Winge DR. Biogenesis of the cytochrome bc(1) complex and role of assembly factors. Biochim Biophys Acta. 2012 Feb;1817(2):276–86. doi: 10.1016/j.bbabio.2011.11.009. Epub 2011 Nov 22. Corrected and republished in: Biochim Biophys Acta. 2012 Jun;1817(6):872-82. doi: 10.1016/j.bbabio.2012.03.003.

52. A. Singharoy, C. Maffeo, K. H. Delgado-Magnero, et al., “Atoms to phenotypes: Molecular design principles of cellular energy metabolism,” Cell, vol. 179, no. 5, 1098–1111.e23, 2019.

53. Pizzorno J. Mitochondria-Fundamental to Life and Health. Integr Med (Encinitas). 2014 Apr;13(2):8–15. PMID: 26770084; PMCID: PMC4684129.

54. Wennerström, H.; Vallina Estrada, E.; Danielsson, J.; Oliveberg, M. Colloidal stability of the living cell. Proc. Natl. Acad. Sci. U. S. A. 2020, 117, 10113−10121

55. Kortemme, T., Kim, D.E. and Baker, D. (2004) ‘Computational alanine scanning of protein-protein interfaces’, Science’s STKE, 2004(219). doi:10.1126/stke.2192004pl2.

56. Kathy Pfeiffer, Vishal Gohil, Rosemary A. Stuart, Carola Hunte, Ulrich Brandt, Miriam L. Greenberg, Hermann Schägger (2003). Cardiolipin Stabilizes Respiratory Chain Supercomplexes. 278 (52) 52873–52880. DOI: 10.1074/jbc.M308366200

57. Trouillard, M.; Meunier, B.; Rappaport, F. Questioning the functional relevance of mitochondrial supercomplexes by time-resolved analysis of the respiratory chain. Proc. Natl. Acad. Sci. U. S. A. 2011, 108, E1027– E1034, DOI: 10.1073/pnas.1109510108

58. Klusch, N. Dreimann, M., Senkler, J., Rugen, N., Kühlbrandt, W., Braun, H-P. (2022) ‘Cryo-EM structure of the respiratory I + III2 supercomplex from Arabidopsis thaliana at 2 Å resolution’, Nature Plants, 9(1), pp. 142–156. doi:10.1038/s41477-022-01308-6.

59. Volkov, A.N. and van Nuland, N.A. (2012) ‘Electron transfer interactome of cytochrome c’, PLoS Computational Biology, 8(12). doi:10.1371/journal.pcbi.1002807.

60. Xu, D. and Zhang, Y. (2012) ‘*ab initio* protein structure assembly using continuous structure fragments and optimized knowledge-based force field’, Proteins: Structure, Function, and Bioinformatics, 80(7), pp. 1715–1735. doi:10.1002/prot.24065.

61. Jo, S. Kim, T., Iyer, V.G., Im, W.. (2008) ‘Charmm-Gui: A Web-based graphical user interface for CHARMM’, Journal of Computational Chemistry, 29(11), pp. 1859–1865. doi:10.1002/jcc.20945.

62. James C. Phillips, David J. Hardy, Julio D. C. Maia, John E. Stone, João V. Ribeiro, Rafael C. Bernardi, Ronak Buch, Giacomo Fiorin, Jérôme Hénin, Wei Jiang, Ryan McGreevy, Marcelo C. R. Melo, Brian K. Radak, Robert D. Skeel, Abhishek Singharoy, Yi Wang, Benoît Roux, Aleksei Aksimentiev, Zaida Luthey-Schulten, Laxmikant V. Kalé, Klaus Schulten, Christophe Chipot, Emad Tajkhorshid; Scalable molecular dynamics on CPU and GPU architectures with NAMD. J. Chem. Phys. 28 July 2020; 153 (4): 044130. 10.1063/5.0014475

63. Xue, L.C. Rodrigues, J.P., Kastritis, P.L., Bonvin, A.M., Vangone, A.. (2016) ‘Prodigy: A web server for predicting the binding affinity of protein–protein complexes’, Bioinformatics, 32(23), pp. 3676–3678. doi:10.1093/bioinformatics/btw514.

64. Comer J, and Aksimentiev A (2012) Predicting the DNA sequence dependence of nanopore ion current using atomic-resolution Brownian dynamics. J. Phys. Chem C 116, 3376–3393.

65. A. T. Baker, R. J. Boyd, D. Sarkar, A. Teijeira-Crespo, C. K. Chan, E. Bates, K. Waraich, J. Vant, E. Wilson, C. D. Truong, M. Lipka-Lloyd, P. Fromme, J. Vermaas, D. Williams, L. Machiesky, M. Heurich, B. M. Nagalo, L. Coughlan, S. Umlauf, P. Chiu, P. J. Rizkallah, T. S. Cohen, A. L. Parker, A. Singharoy, M. J. Borad, ChAdOx1 interacts with CAR and PF4 with implications for thrombosis with thrombocytopenia syndrome. Science Advances, 2021, 7(49), eabl8213.

66. Mrinal Shekhar, Chitrak Gupta, Kano Suzuki, Chun Kit Chan, Takeshi Murata, and Abhishek Singharoy, “Revealing a Hidden Intermediate of Rotatory Catalysis with X-ray Crystallography and Molecular Simulations,” ACS Central Science 2022 8 (7), 915–925

67. W. Humphrey, A. Dalke, and K. Schulten, “VMD – Visual Molecular Dynamics,” J Mol Graph, vol. 14, no. 1, pp. 33–38, 1996.

68. E. Jurrus, D. Engel, K. Star, et al., “Improvements to the apbs biomolecular solvation software suite,” Protein Science, vol. 27, no. 1, pp. 112–128, 2018.

69. N. A. Baker, D. Sept, S. Joseph, M. J. Holst, and J. A. McCammon, “Electrostatics of nanosystems: Application to microtubules and the ribosome,” Proceedings of the National Academy of Sciences, vol. 98, no. 18, pp. 10 037–10 041, 2001.

70. J. Cohen, A. Arkhipov, R. Braun, and K. Schulten, “Imaging the migration pathways for O2, CO, NO, and Xe inside myoglobin,” Biophysical Journal, vol. 91, pp. 1844–1857, 2006.

71. Miao, Y., Feher, V.A. and McCammon, J.A. (2015) ‘Gaussian accelerated molecular dynamics: Unconstrained enhanced sampling and free energy calculation’, Journal of Chemical Theory and Computation, 11(8), pp. 3584–3595. doi:10.1021/acs.jctc.5b00436.

72. Punjani, A. Rubinstein, J.L., Fleet, D.J., Brubaker, M.A.. (2017) ‘CryoSPARC: Algorithms for rapid unsupervised cryo-em structure determination’, Nature Methods, 14(3), pp. 290–296. doi:10.1038/nmeth.4169.

73. Bepler, T., Morin, A., Rapp, M. et al. Positive-unlabeled convolutional neural networks for particle picking in cryo-electron micrographs. Nat Methods 16, 1153–1160 (2019). 10.1038/s41592-019-0575-8

74. Huang, J., MacKerell Jr, A.D. (2013) CHARMM36 all-atom additive protein force field: Validation based on comparison to NMR data. 34 (25) 2135–2145. doi:10.1002/jcc.23354

75. Melih Sener, Johan Strumpfer, Abhishek Singharoy, C Neil Hunter, Klaus Schulten (2016) Overall energy conversion efficiency of a photosynthetic vesicle. eLife 5:e09541.

