## Supplemental Information for "Transient protein structure guides surface diffusion pathways for electron transport in membrane supercomplexes"


SI Figures (ordered w.r.t their order of appearing in the main text)

- SI figures for Figure 1.
  - SI1.1 Sequence comparison for QCR6s from different species
  - SI1.2: Alphafold2 vs. MELD comparison of QCR6 folding.
  - SI1.3: CryoEM map and model with MD simulation snapshots
  - SI1.4: Dissociation constant (Kd) values between cyt. c and either CIII or CIV.
  - SI1.5: QCR6 helicities probed by GaMD
- SI figures for Figure 2.
  - SI2.1: Work calculations from SMD
- SI figures for Figure 3.
  - SI3.1 Transition matrix between different states of cyt. c under the presence of CL and 2 QCR6s.
  - SI3.2: Impacts of QCR6 on cyt. c equilibrium population distributions and transfer kinetics
- SI figures for Figure 4.
  - SI4.1: Electrostatic profiles in the vicinity of QCR6
  - SI4.2: QCR6 conformation regulates the association rates between cyt. c, CIII, and CIV.
  - SI4.3: Cyt. c associations on CIII and CIV in the absence of anionic lipids.
  - SI4.4: Residence time of cyt. c on different parts of the supercomplex-membrane system.
  - SI4.5: Fractions of cyt. c positioned in the vicinity of the supercomplex-membrane system along BD simulations.
  - SI4.6: The fraction of cyt. c remaining in the bulk solution along BD simulations without the presence of cardiolipins.
  - SI4.7: Association rate of cyt. c to the supercomplex in the absence of CL, with impacts from removing the flexible domain of QCR6.
  - SI4.8: Impacts of the presence of CL and QCR6 on the residence time of cyt. c on supercomplex associations.
  - SI4.9:Impacts of the presence of CL on the residence time of cyt. c - supercomplex associations regarding the QCR6 Trunc variant.
- SI figures for Figure 5
  - SI5.1: Mobility of the supercomplex upon removal of QCR6. (should go to SI for 4)
- SI figures for Figure 6
  - SI6.1: Hotspots for QCR6-cyt. c associations under the presence of CL, as being sampled by SMD.
  - SI6.2 Impact of proposed mutations on cyt. c’s diffusion profile in the proximity of the supercomplex.
  - SI6.3 Tethers for membrane localization of charge carrier proteins from primitive heliobacterial species do not show QCR-like $CX_9C$ sequence signatures.

SI6.4 Cross-correlations (CC) between simulated cyt. *c* surface diffusion map and experimental data.
SI figures for Method
SI7.1 Workflow for modeling, simulations and analysis
SI7.2 Modeling of QCR6 folds
SI7.3: Spatial distributions of spaces with weak electropositivity and spaces with weak electronegativity.

### SI figures for **Figure 1**.

#### SI1.1 Sequence comparison for QCR6s from different species.

The multi-sequence alignment (MSA), performed by CLUSTAL, demonstrated the unusually long sequence for yeast's QCR6. An Arginine residue that is conserved across multiple species, and is found to interact with cardiolipins in our current model, guiding the 2D diffusion of cyt. *c* is highlighted by a red box.

```
CLUSTAL O(1.2.4) multiple sequence alignment

SP|P07919|QCR6_HUMAN ----------------------------------------------MGLEDEQK--MLTE 12
SP|P00126|QCR6_BOVIN ----------------------------------------------MGLEDEQR--MLTG 12
SP|Q5M9I5|QCR6_RAT   ----------------------------------------------MGLEDERK--MLTG 12
SP|Q8SPH5|QCR6_MACFA ----------------------------------------------MGLEDERK--MLTE 12
SP|P00127|QCR6_YEAST MGMLELVGEYWEQLKITVVPVVAAAEDDDNEQHEEKAAEGEEKEEENGDEDEDEDEDEDD 60
                                                                    * *** .

SP|P07919|QCR6_HUMAN SGDPEEEEEEEEELVDPLTTVREQCEQLEKCVKARERLELCDERVSSR--------SHTE 64
SP|P00126|QCR6_BOVIN SGDPKEEEEEEEELVDPLTTVREQCEQLEKCVKARERLELCDERVSSR--------SQTE 64
SP|Q5M9I5|QCR6_RAT   SGDPKEE--EEEELVDPLTTVREHCEQLEKCVKARERLESCDRRVSSR--------SQTE 62
SP|Q8SPH5|QCR6_MACFA SGDPEKEEEEEEELVDPLTTVREQCEQLEKCVKARERLELCDKRVSSR--------SRTE 64
SP|P00127|QCR6_YEAST DDDDDEDEEEEEEVTDQLEDLREHFKNTEEGKALVHHYEECAERVKIQQQQPGYADLEHK 120
                     ..* .::  ****:.* *  :**: :: *:     .: * * .**. :         . :

SP|P07919|QCR6_HUMAN EDCTEELFDFLHARDHCVAHKLFNNLK 91
SP|P00126|QCR6_BOVIN EDCTEELLDFLHARDHCVAHKLFNSLK 91
SP|Q5M9I5|QCR6_RAT   EDCTEELFDFLHARDHCVAHKLFKSLK 89
SP|Q8SPH5|QCR6_MACFA EDCTEELLDFLHARDHCVAYKLFNNLK 91
SP|P00127|QCR6_YEAST EDCVEEFFHLQHYLDTATAPRLFDKLK 147
                     ***.**::.: *  * ..* :**..**
```

#### SI1.2: Alphafold2 vs. MELD comparison of QCR6 folding.

(A). Each of the five models generated using Alphafold2 (blue) are aligned with States 2 (orange) and 3 (magenta) from MELD. RMSD values between the Alphafold2 model and states 2 and 3 (S2 and S3, respectively) are listed below each model. (B) Shows sequence coverage of the structures that were used to create the final output model. The X-axis shows the position of the sequence (of our structure) and Y-axis shows how many of the database proteins had sequence coverage at the said position. and confidence values from the predicted local distance difference test (pLDDT). A pLDDT score of > 90 shows high modeling accuracy, while scores between 70–90 are considered generally modeled well. (C) 2D plot of the Predicted Aligned Error (PAE). The blue regions of QCR6 represent well-defined positions and orientations.

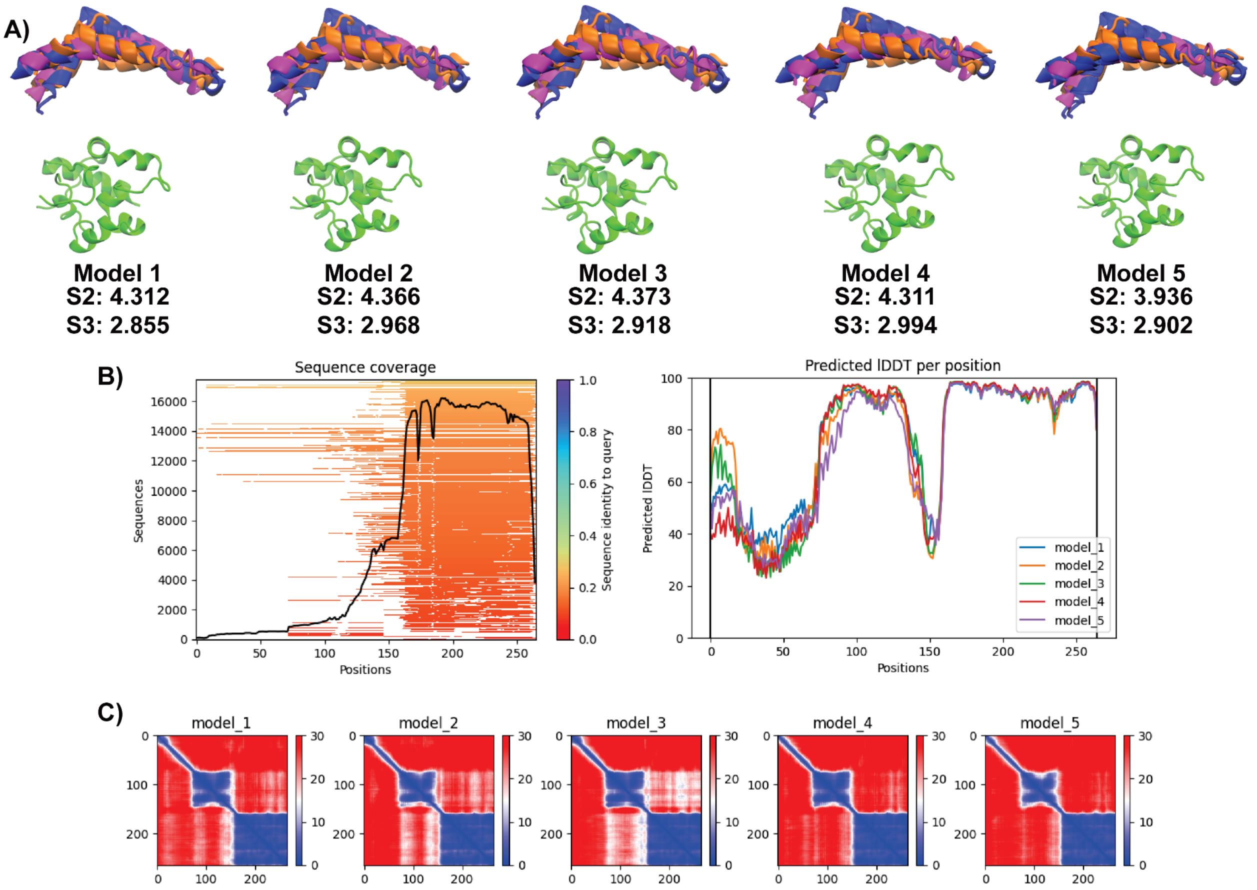


#### SI1.3 Dissociation constant ($K_d$) values between cyt. *c* and either CIII or CIV.

(A) During SMD, cytochrome c's interaction energy with CIII (also known as cytochrome $bc_1$) gradually weakens as cyt. *c* approaches CIV, wherein the carrier-protein interactions improve (B). (C) $K_d$ values computed using the Prodigy web-server with poses sampled from SMD simulations (denoted by _MD) and from BD simulations (denoted by _BD). (D) Shown are the highlighted 7 lysine residues (blue; from cyt. $c_2$) that are conserved across proteins of the cyt. *c* family and are used to establish strong associations with CIII's acidic (in red).

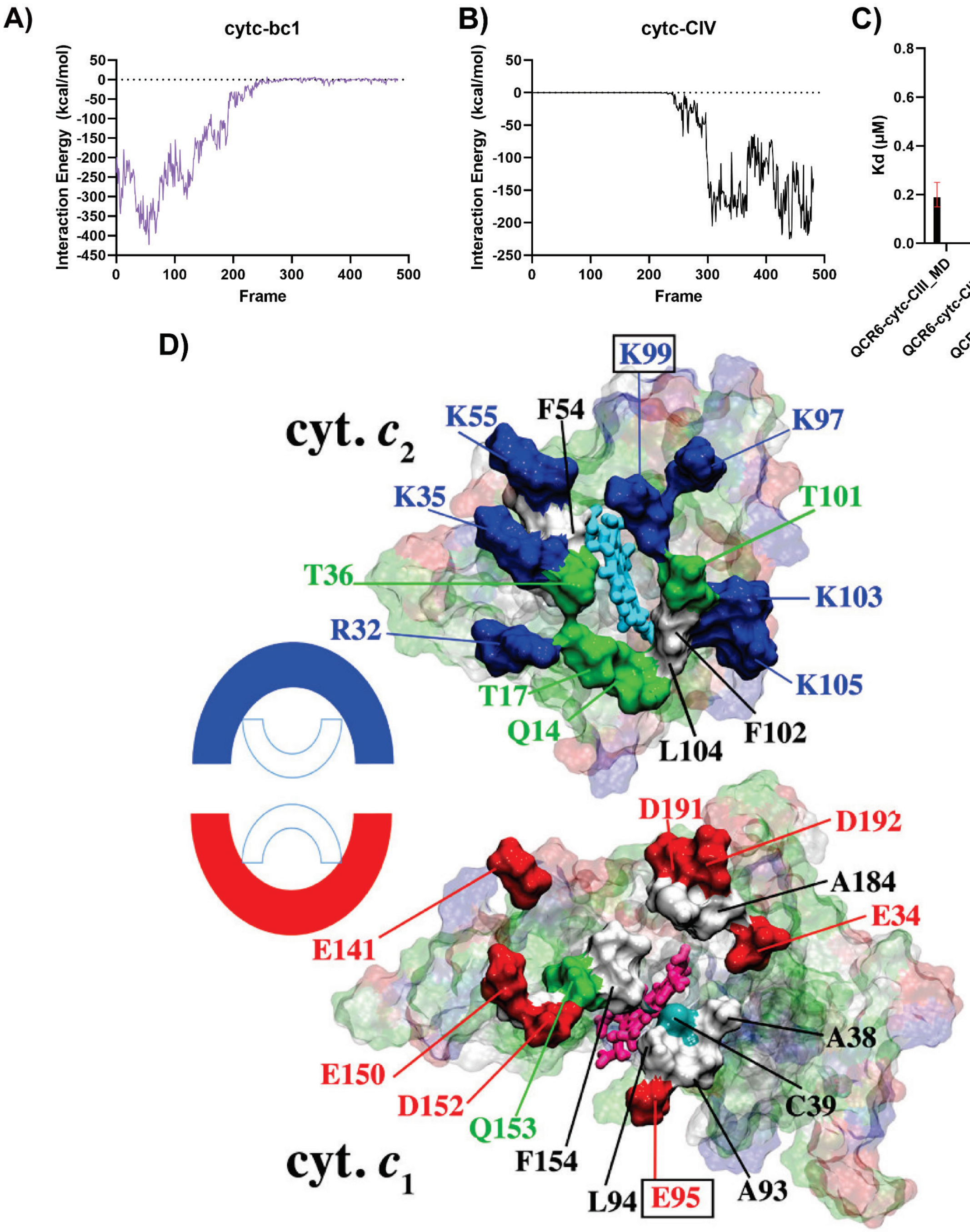
A)
cytc-bc1
Interaction Energy (kcal/mol)
Frame
B)
cytc-CIV
Interaction Energy (kcal/mol)
Frame
C)
Prodigy Kd
Kd (μM)
QCR6-cytc-CIII_MD
QCR6-cytc-CIII_BD
QCR6-cytc-CIV_MD
QCR6-cytc-CIV_BD
D)
cyt. c2
K99
K55
F54
K97
K35
T101
T36
K103
R32
K105
T17
F102
Q14
L104
D191
D192
A184
E34
E141
A38
E150
C39
D152
Q153
F154
A93
cyt. c1
L94
E95

#### SI1.4: Cryo-EM map and model with MD simulation snapshots

(A) A starting model for the QCR6 N-terminal region is constructed by folding its first 76 residues using MELD-accelerated MD simulations in the presence of cyt. *c* and soluble residues from CIII that are within 14 Å of the cyt. *c*. Four snapshots (denoted states 0–3) representing the local interaction energy minima along the protein folding simulation are illustrated, demonstrating the flexibility of the QCR6 hinge (B) Interaction energies between cyt. *c* and QCR6 plotted as a function of end-to-end distances, with states 0-3 highlighted, elucidating that the more compact states offer stronger interactions between cyt. *c* and QCR6.

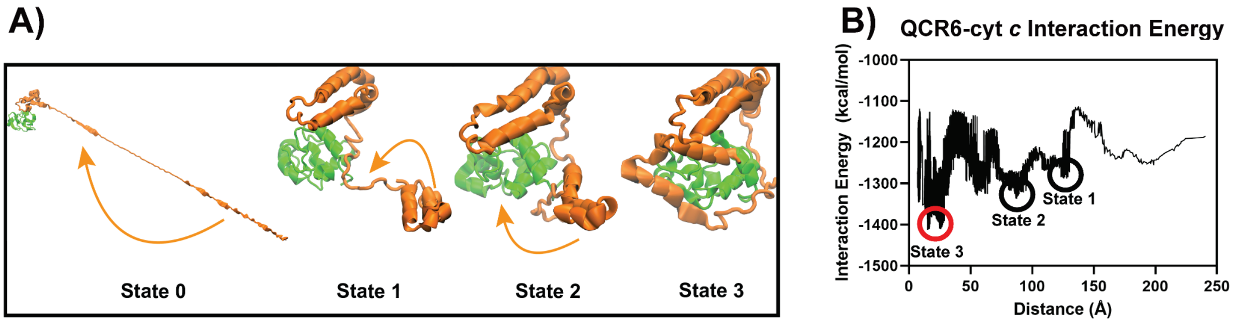


#### SI1.5 QCR6 helicities probed by GaMD

Helicity of QCR6 was probed by GaMD. Data from GaMD initiated with different snapshots (*t*=0, *t*=150 ns, *t*=480 ns) from SMD simulations (Fig. 3) were colored in blue, orange, and green, respectively. The data are presented here as distributions over the percentage (30% to 90%) of QCR6's content being structured as helices, monitored using the alpha collective variable in VMD. Distributions for GaMD performed with and without cyt. *c* are shown in (A) and (B). respectively.

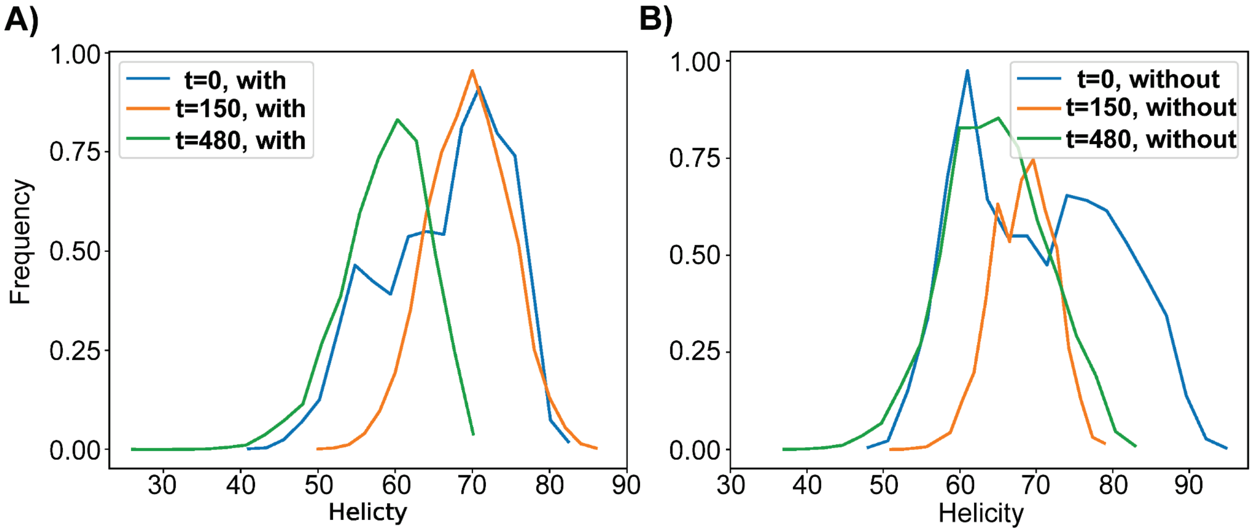

SI figures for **Figure 2**.

#### SI2.1 Work calculations from SMD

During the SMD replicates, the average work required as cytochrome c approaches CIV increases significantly between the POPC and POPC + CL models, showing that the presence of CL decreases the energy cost required for cytochrome c transport between CIII and CIV.

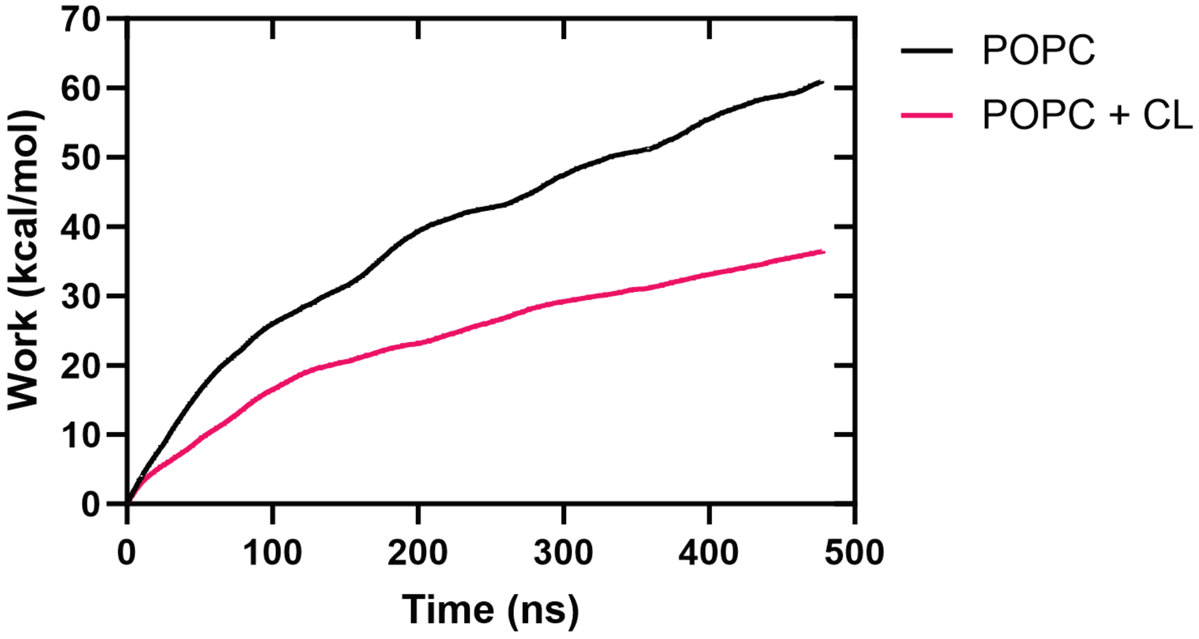

### SI figures for Figure 3.

#### SI3.1 Transition matrix between different states of cyt. *c* under the presence of CL and 2 QCR6s.

For any cyt. *c* in BD simulations, depending on its interaction with the supercomplex, it is classified to belong to 1 of the following 8 classes: **(1)** *c3m1* (associated with CIII of monomer 1 of the supercomplex); **(2)** *c3m2* (associated with CIII of monomer 2 of the supercomplex); **(3)** *c4m1* (associated with CIV of monomer 1 of the supercomplex); **(4)** *c4m2* (associated with CIV of monomer 2 of the supercomplex); **(5)** *qcm1* (associated with QCR6 of monomer 1 of the supercomplex); **(6)** *qcm2* (associated with QCR6 of monomer 2 of the supercomplex); **(7)** *Membrane* (associated with the membrane); **(8)** *Bulk* (remaining in the bulk). The frequency for each possible transition between these states, computed out of 31680 snapshots collected over 600μs of BD simulations WT supercomplex embedded in PC and CL-membrane (Table 1), is shown in the transition matrix below, with the color of each cell following the color scale on the right of the matrix.

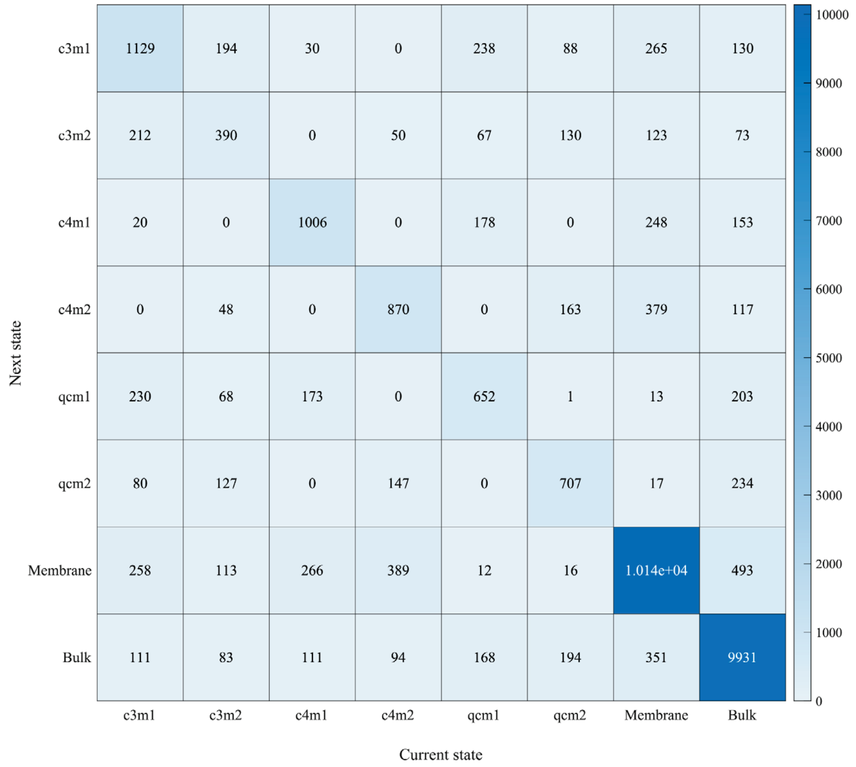


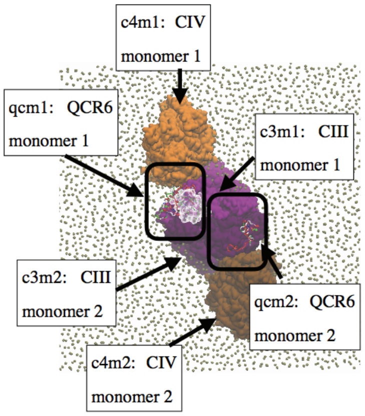

#### SI3.2: Impacts of QCR6 on cyt. *c* equilibrium population distributions and transfer kinetics

The rate matrices that summarize the transfer rates across 8 different cyt. *c* bound states used in analyzing BD simulations (See SI3.1) were computed (See Method: Rate matrix computations and population transfer simulations) for the wild type (WT) megacomplex, labeled QCR6, and its mutant (ΔQCR6). These matrices, together with differences across their elements, where the rate matrix for ΔQCR6 was subtracted from that for WT, are given in (A). The 8 different bound states were concatenated to give a high-level view on the impact of QCR6 on cyt. *c* transfer kinetics. As c3m1, c3m2, qcm1, and qcm2 represent individual components of the dimeric CIII, bound states on them were grouped as bound states on CIII. Similarly, bound states on c4m1 and c4m2, the 2 CIV monomers, were grouped as bound states on CIV. Under these concatenations, population transfers from CIII to CIV, and their loss to the membrane or the bulk, were simulated following the population transfers equations (See Method: Rate matrix computations and population transfer simulations). The simulation results (B) show that, with a population of cyt. *c* starting on CIII, WT achieves a higher level of equilibrium population for megacomplex-bound cyt. *c*, both for CIII and CIV. WT thus provides a higher success rate for cyt. *c* to be transferred from CIII to CIV. A zoom in to the 1st 0.5 μs cyt. *c* population transfer from CIII to CIV (C) shows that the mutant achieves a higher initial transfer rate than WT. For each of the mutant and WT, this process of initial population transfer was fitted by a linear fit (Linear fit 1) with the resulting slope being the estimate of the corresponding initial population transfer rate. As the cyt. *c* populations transferred to CIV encounter a significant decay in the case of the mutant but not WT. An effective transfer rate was also estimated by the slope (Linear fit 2) between the starting point at t = 0 μs and the time point where the cyt. *c* population first decay (mutant) /raise (WT) to 10% above the respective equilibrium cyt. *c* population.

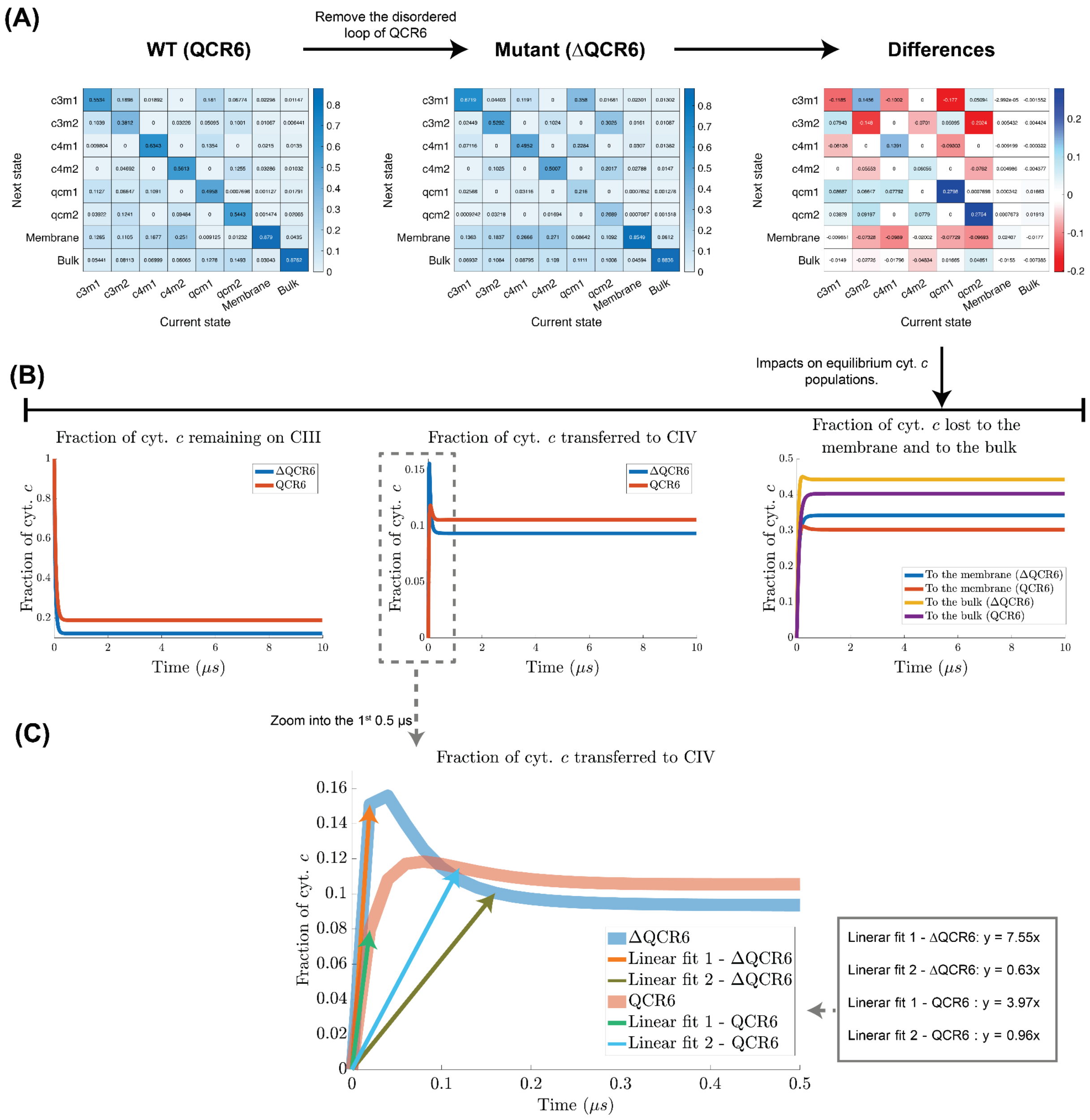

(A)
WT (QCR6)
Remove the disordered loop of QCR6
Mutant (ΔQCR6)
Differences
Next state
Current state
c3m1
c3m2
c4m1
c4m2
qcm1
qcm2
Membrane
Bulk
Impacts on equilibrium cyt. c populations.
(B)
Fraction of cyt. c remaining on CIII
Fraction of cyt. c transferred to CIV
Fraction of cyt. c lost to the membrane and to the bulk
Fraction of cyt. c
Time (μs)
ΔQCR6
QCR6
To the membrane (ΔQCR6)
To the membrane (QCR6)
To the bulk (ΔQCR6)
To the bulk (QCR6)
Zoom into the 1st 0.5 μs
(C)
Fraction of cyt. c transferred to CIV
Fraction of cyt. c
Time (μs)
ΔQCR6
Linear fit 1 - ΔQCR6
Linear fit 2 - ΔQCR6
QCR6
Linear fit 1 - QCR6
Linear fit 2 - QCR6
Linear fit 1 - ΔQCR6: y = 7.55x
Linear fit 2 - ΔQCR6: y = 0.63x
Linear fit 1 - QCR6 : y = 3.97x
Linear fit 2 - QCR6 : y = 0.96x

### SI figures for [Figure 4](#).

#### SI4.1: Electrostatic profiles in the vicinity of QCR6

Electrostatic profiles in the vicinity of QCR6 were determined using APBS (see Method: Brownian Dynamics (BD)) for different SMD snapshots used to source BD simulations. Snapshots corresponding to time points of 0, 220 ns, 400 ns, and 480 ns from SMD simulations, which both exhibit local minimums for interaction energy between cyt. *c* and QCR6 as well as are used to illustrate BD simulations results in **Fig. 4**, are shown below. Profiles computed, shown at the upper row, are visualized as contour surfaces of the electrostatic potential at 0.5k$_B$T (red) and -0.5k$_B$T (blue). The electronegative contour surfaces of these potentials, shown in red, pink, white, and blue, respectively, are overlaid on the electrostatic potential (black) of the ensemble average model. This model is an arithmetic average of the shown profiles and it mimics an environment with QCR6 in motion. All electrostatic profiles shown here, as well as, the averaged model, show that the surface of the supercomplex-membrane system is overall electronegative with a clearly protruding electronegative region established by the presence of QCR6. These regions of electronegativity are able to attract the electropositive side of cyt. *c*, potentially enhancing the chances for cyt. *c* to be found around the supercomplex.

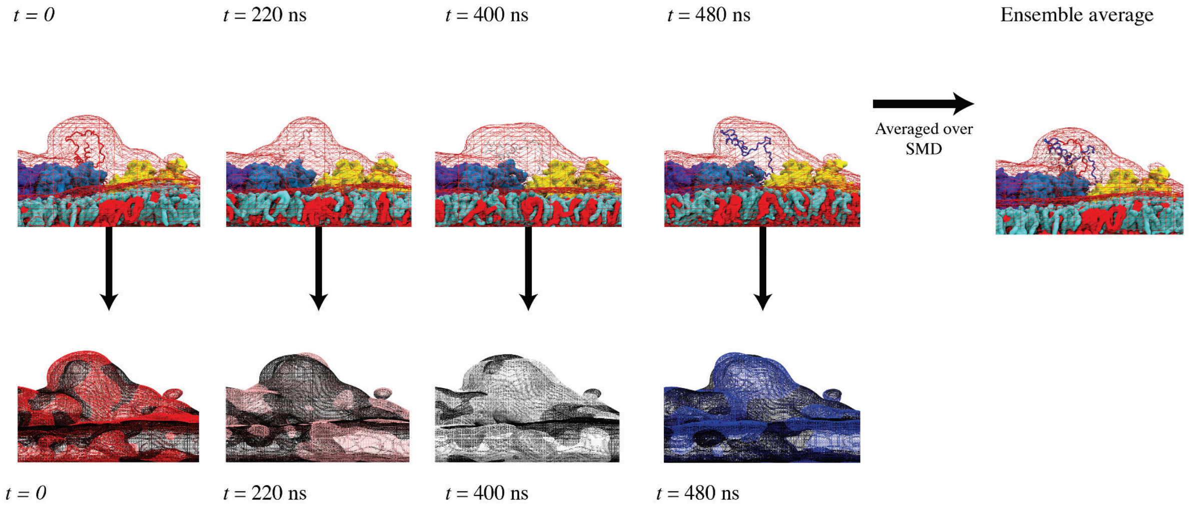


#### SI4.2: QCR6 conformation regulates the association rates between cyt. *c*, CIII, and CIV.

Shown are the individual fractions of cyt. *c*, out of those associated with the supercomplex, for CIIIs, CIVs, and QCR6s, as being probed by BD simulations at discrete stages (marked by “X” in each plot) extracted from SMD simulations under the presence of CL. Fractions for the QCR6 that interacts with the cyt. *c* undergoing SMD and its neighboring CIII and CIV are shown in (left), where CIV becomes substantially more attractive to cyt. *c* than CIII does as QCR6 fully unfolds towards CIV. Fractions for the QCR6 that do not interact with the cyt. *c* undergoing SMD and its neighboring CIII and CIV are given in (right).

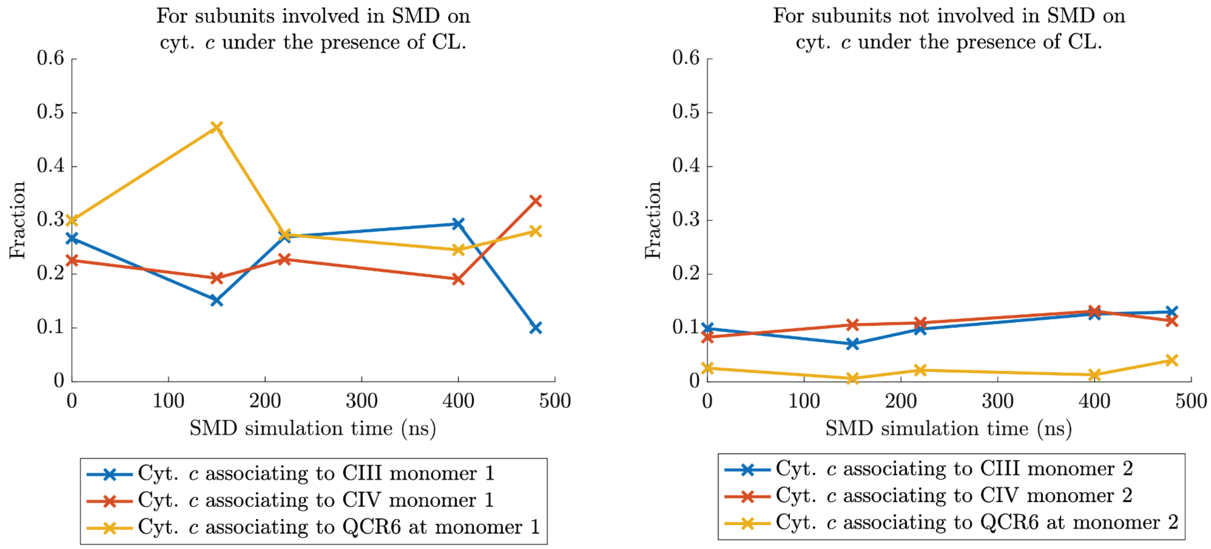


#### SI4.3: Cyt. *c* associations on CIII and CIV in the absence of anionic lipids.

In (A) and (B), hotspots for cyt. *c* associations on CIII and CIV in are shown, respectively. Notations used in each molecular image and in each histogram plot in (A) and (B) follow those in **Fig. 4**, residue colors reflect their frequency of interacting with cyt. *c*. Individual fractions of cyt. *c* associated with CIIIs, CIVs, and QCR6s are shown in (C) and (D). As a comparison and as to supplement details to **Fig. 4**, histogram plots in line with those shown in (A) and (B) are given in (E). Note, that the distribution of cyt. *c* associations to CIII changes minimally even when the QCR6 is displaced towards CIV. This is in stark contrast with cyt. *c*-CIII associations in the presence of cardiolipins. However, a growth in the vicinity of CIV is observed both in the presence and absence of the cardiolipins.

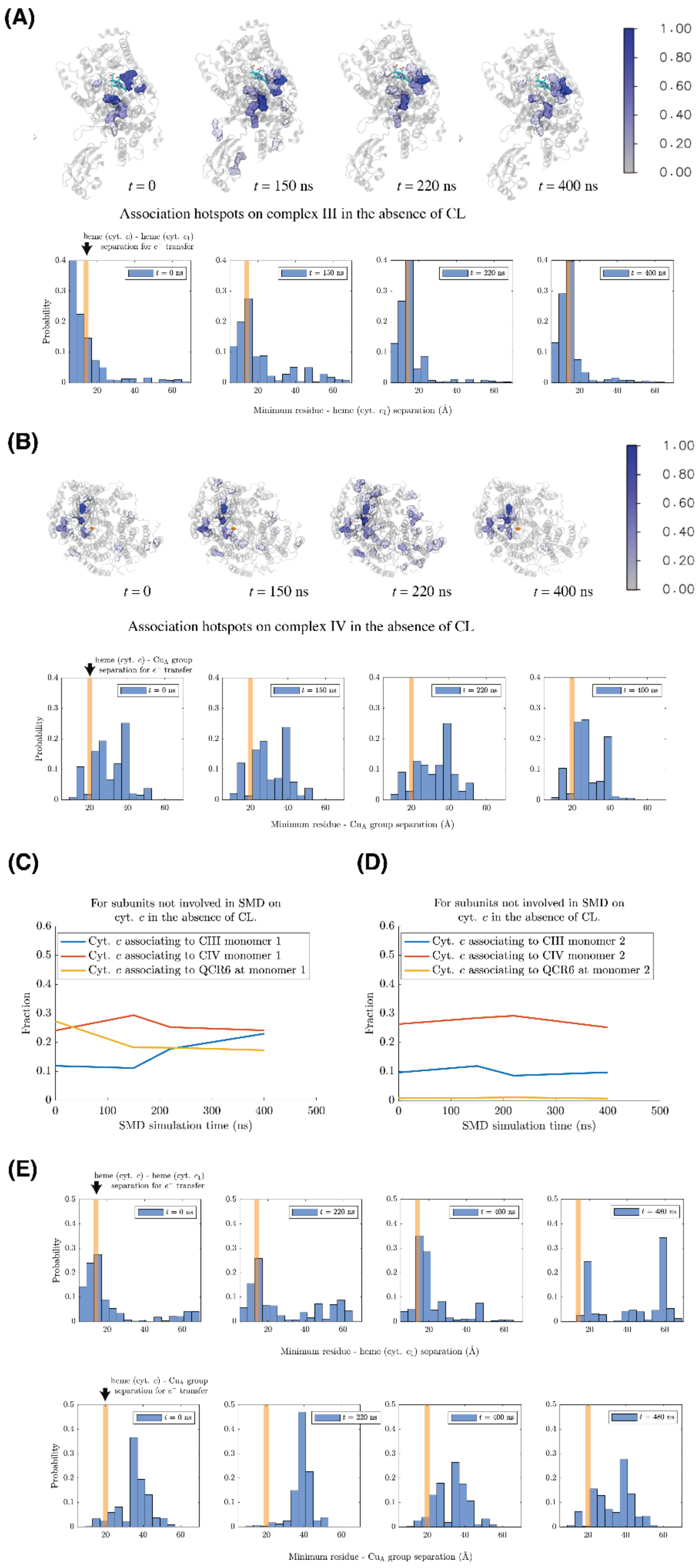
(A)
t = 0
t = 150 ns
t = 220 ns
t = 400 ns
1.00
0.80
0.60
0.40
0.20
0.00
Association hotspots on complex III in the absence of CL
heme (cyt. c) - heme (cyt. c1) separation for e− transfer
Probability
Minimum residue - heme (cyt. c1) separation (Å)
(B)
Association hotspots on complex IV in the absence of CL
heme (cyt. c) - CuA group separation for e− transfer
Minimum residue - CuA group separation (Å)
(C)
For subunits not involved in SMD on cyt. c in the absence of CL.
Cyt. c associating to CIII monomer 1
Cyt. c associating to CIV monomer 1
Cyt. c associating to QCR6 at monomer 1
Fraction
SMD simulation time (ns)
(D)
For subunits not involved in SMD on cyt. c in the absence of CL.
Cyt. c associating to CIII monomer 2
Cyt. c associating to CIV monomer 2
Cyt. c associating to QCR6 at monomer 2
Fraction
SMD simulation time (ns)
(E)
heme (cyt. c) - heme (cyt. c1) separation for e− transfer
Minimum residue - heme (cyt. c1) separation (Å)
heme (cyt. c) - CuA group separation for e− transfer
Minimum residue - CuA group separation (Å)

##### SI4.4: Residence time of cyt. *c* on different parts of the supercomplex-membrane system.

In the presence of CL, distributions of the residence time for cyt. *c* associated with the supercomplex are shown in (A). Population with a higher residence-time on the supercomplex is higher when the QCR6 is present. These positions are spots on the supercomplex where any heavy atom of cyt. *c* are separated from those of CIII for <6 Å if QCR6 is present. Distributions of the residence time for cyt. *c* associated with the supercomplex at positions far away from CIII (with an inter-heavy-atom-separation >6 Å) are shown in (B). In (C), distributions of the residence time of cyt. *c* on the cardiolipin-containing membrane are given with interface separation (measured between heavy atoms of the membrane and those of cyt. *c*) of < 6 Å.

**(A)**

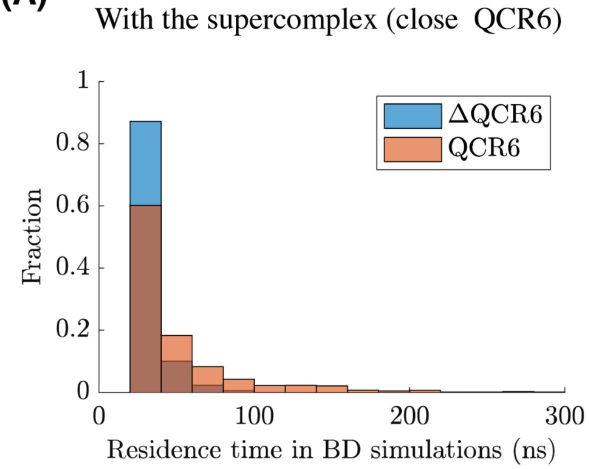


**(B)**

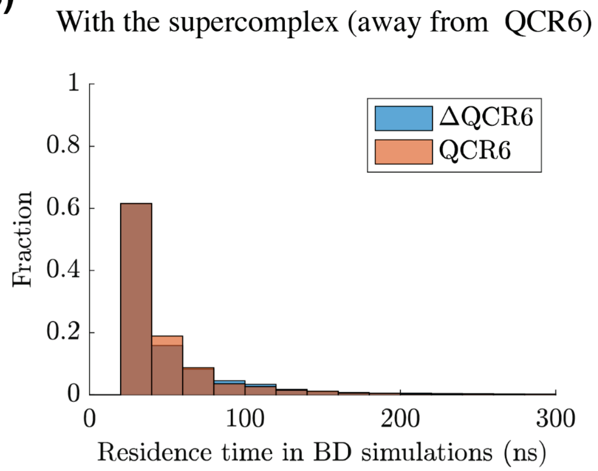


**(C)**

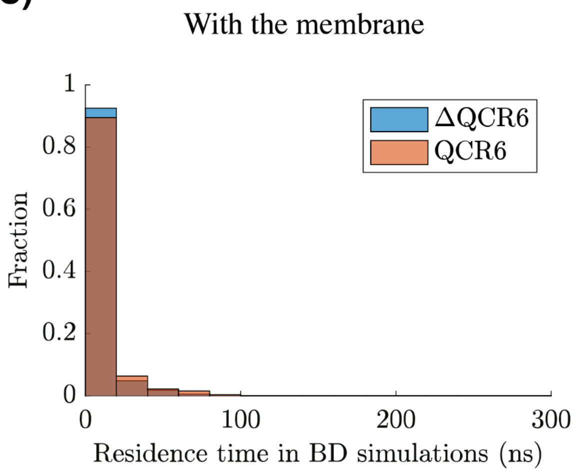

#### SI4.5: Fractions of cyt. *c* positioned in the vicinity of the supercomplex-membrane system along BD simulations.

The below shows the fraction of cyt. *c* positioned in the vicinity (within 6Å heavy-atom-to-heavy-atom separation) of either the supercomplex (SC) or the membrane embedding it. The data are presented for the WT (right) and the mutant (ΔQCR6, left), where the acidic hinge domain of QCR6 was truncated. For both cases, the presence of cardiolipin molecules (CL, blue) plays a critical role in drawing cyt. *c* towards the vicinity of either the SC or the membrane. Yet, upon the absence of CL (red) and compared with the mutant, WT retains its strong ability to attract cyt. *c* to either SC or the membrane by processing the hinge domain of QCR6.

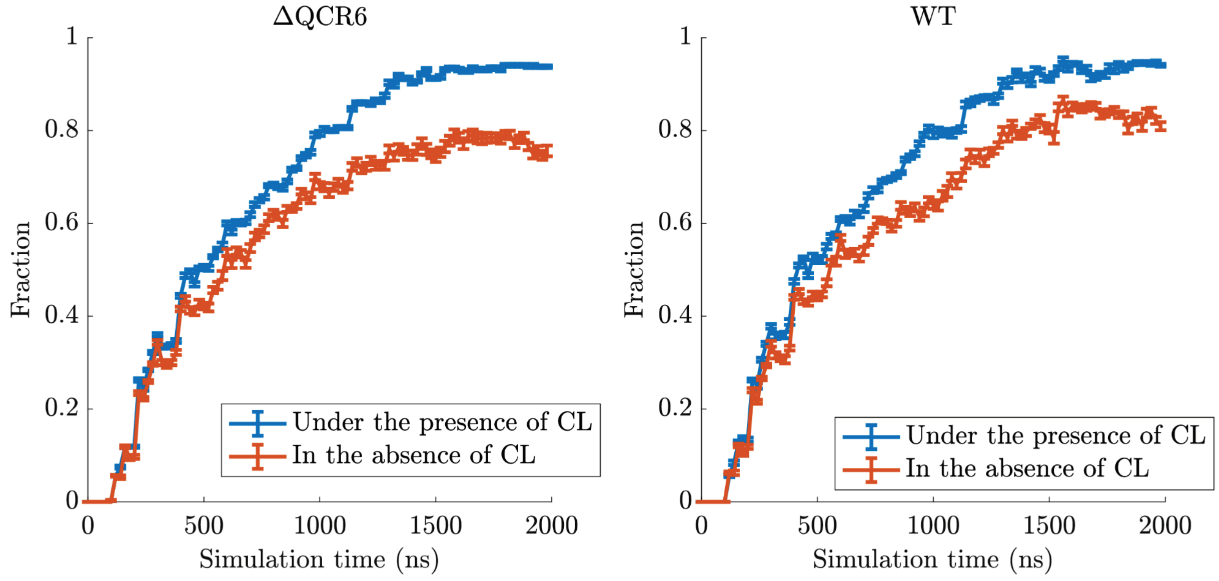


#### SI4.6: The fraction of cyt. *c* in the bulk solution along BD simulations without the presence of cardiolipins.

The below shows the fraction of cyt. *c* in the bulk solution in the absence of cardiolipins for both the case of a WT supercomplex (labeled QCR6) and that of the mutant (labeled ΔQCR6), where the hinge domain of QCR6 was removed. A cyt. *c* was considered to be in the bulk solution if it does not come close to either the supercomplex (SC) or the membrane with a heavy-atom-to-heavy-atom separation less than 6Å.

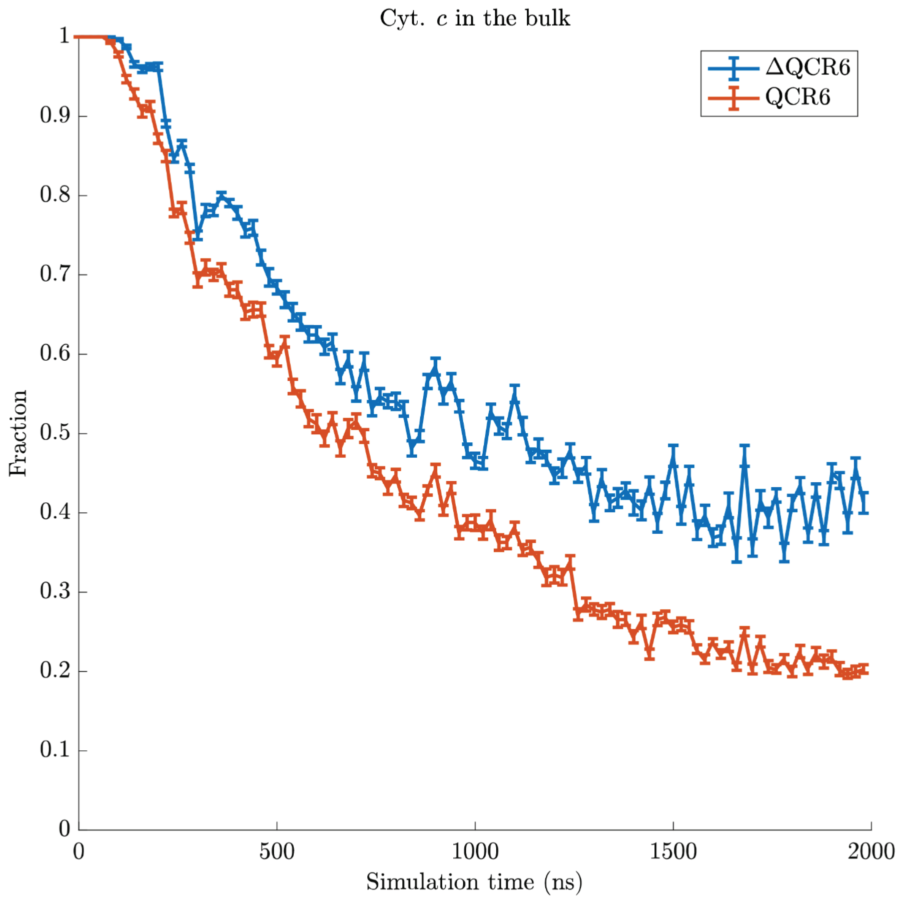


#### SI4.7: Association rate of cyt. *c* to the supercomplex in the absence of CL, with impacts from removing the flexible domain of QCR6.

The number of cyt. *c* associations to either the supercomplex or the membrane in the absence of cardiolipin molecules (CL) was computed and was normalized against the respective solvent accessible surface area (SASA). The resulting numbers of associations per SASA are shown below for both the WT (QCR6) and the mutant (ΔQCR6), where the hinge domain of QCR6 was removed.

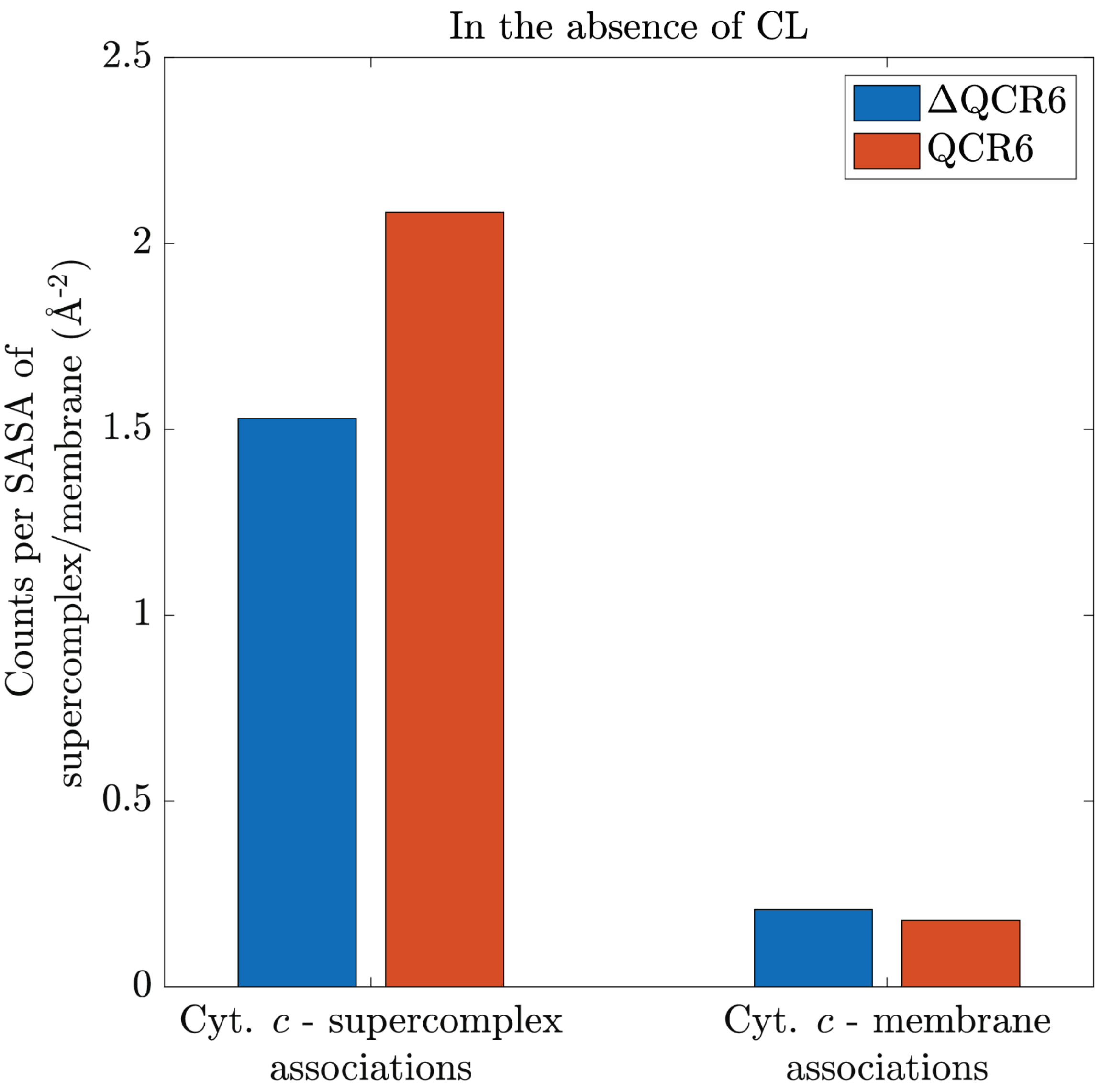


#### SI4.8: Impacts of the presence of CL and QCR6 on the residence time of cyt. *c* on whole-supercomplex associations

Data for all cyt. *c* associations to the supercomplex, including those with prior associations with the membrane lipids, are given in (A). Lots of transient associations, which correspond to cyt. *c* colliding the supercomplex, are observed in the presence of CL. Removing this transient behavior by checking on data for cyt. *c* associations to the supercomplex without any prior association with the membrane lipids (B) shows an improvement on the residence time under the presence of CL, where the improvement is clearly noticeable in the mutant (ΔQCR6) case and is relatively marginal for the WT (QCR6) case, suggesting the involvement of membrane-wise electronegativity in stabilizing cyt. *c* - supercomplex associations These data sets correspond to associations in which cyt. *c* approaching the supercomplex entirely through diffusing in the bulk or on the supercomplex. Therefore, the presence of anionic CL stabilizes cyt. *c*-supercomplex associations that already exist in the absence of CL , and supplements these associations with a pool of membrane-localized cyt. *c* that offers additional transient associations. It should be noted that these transient associations could lead to subsequent cyt. *c* diffusion on the supercomplex. In such a case, the stabilizing effect of CL on cyt. *c*'s residence on the supercomplex also benefits lipid-mediated cyt. *c* associations to the supercomplex.

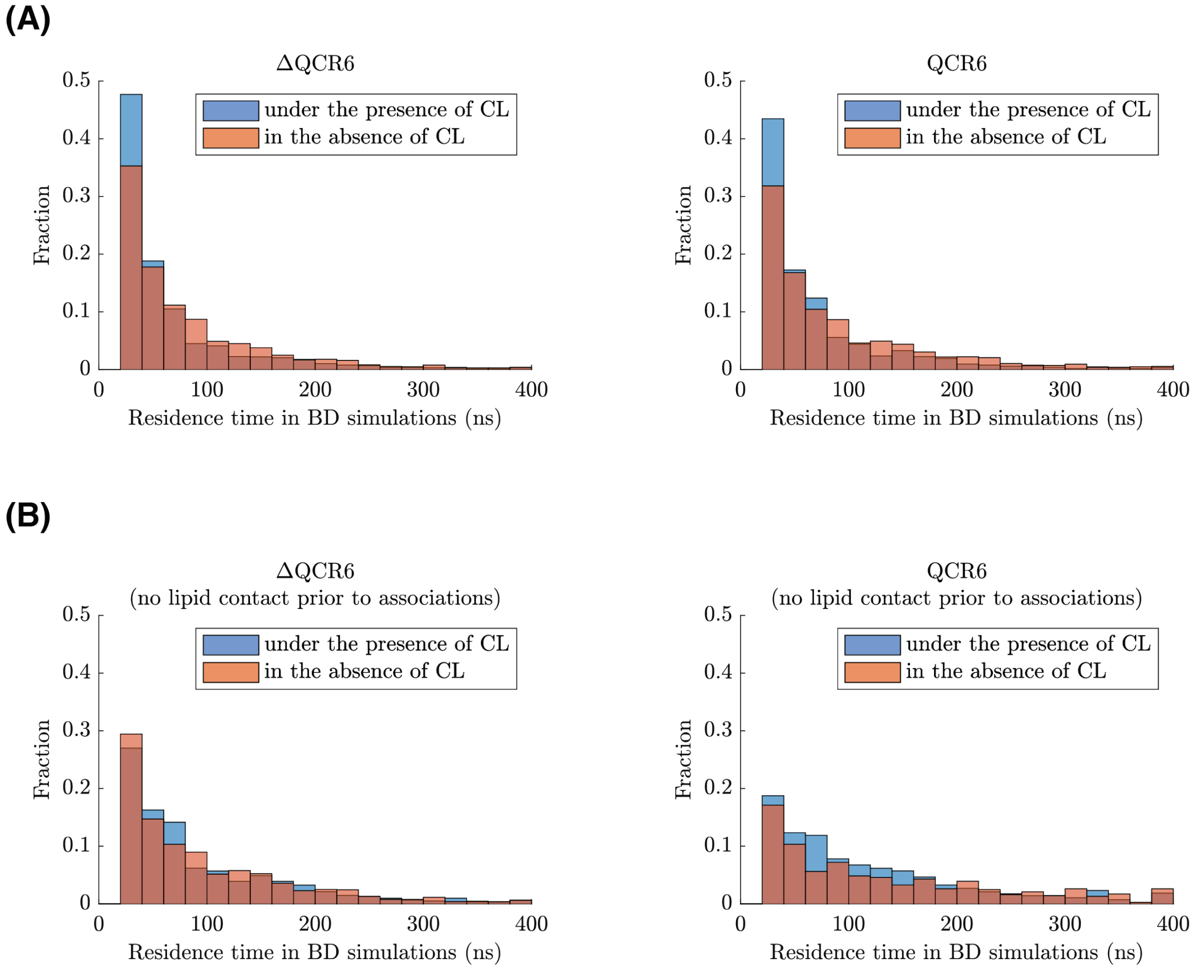


#### SI4.9: Impacts of the presence of CL on the residence time of cyt. *c* - supercomplex associations regarding the QCR6 Trunc variant.

The significance of the additional data set (left) with cyt. *c* associated with the supercomplex without any prior association with the membrane is explained in **Fig. SI3.6**.

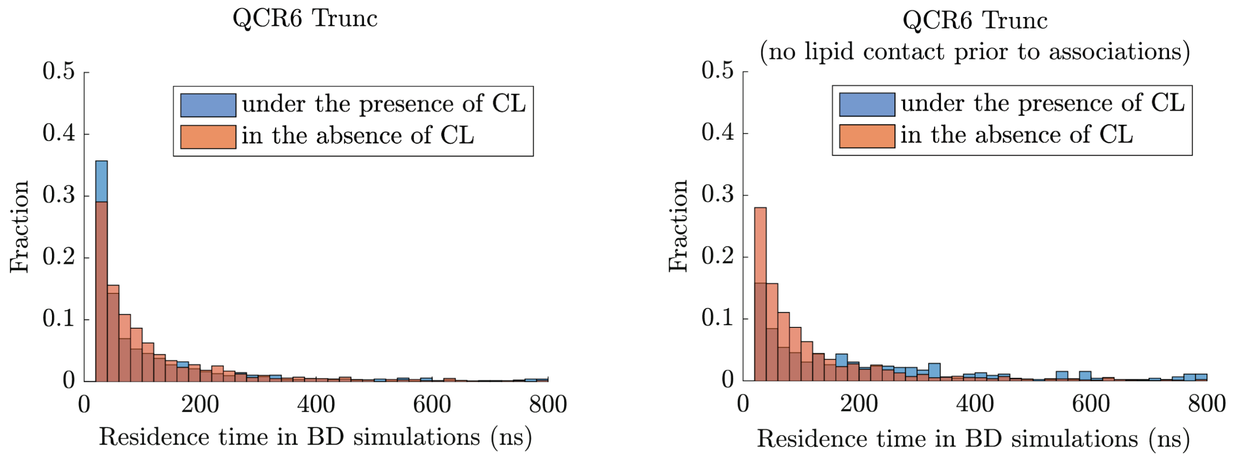
QCR6 Trunc
QCR6 Trunc
(no lipid contact prior to associations)
under the presence of CL
in the absence of CL
Fraction
Residence time in BD simulations (ns)
0
0.1
0.2
0.3
0.4
0.5
0
200
400
600
800

### SI figures for Figure 6

#### SI5.1: Mobility of the supercomplex upon removal of QCR6.

Cryo-EM densities for the wild type supercomplex and the mutant with the whole QCR6 removed, labeled as ΔQCR6, are shown in (A). In (B), multiple views of the ΔQCR6's cryo-EM density map and its corresponding resolution profiles are shown.

**(A)**

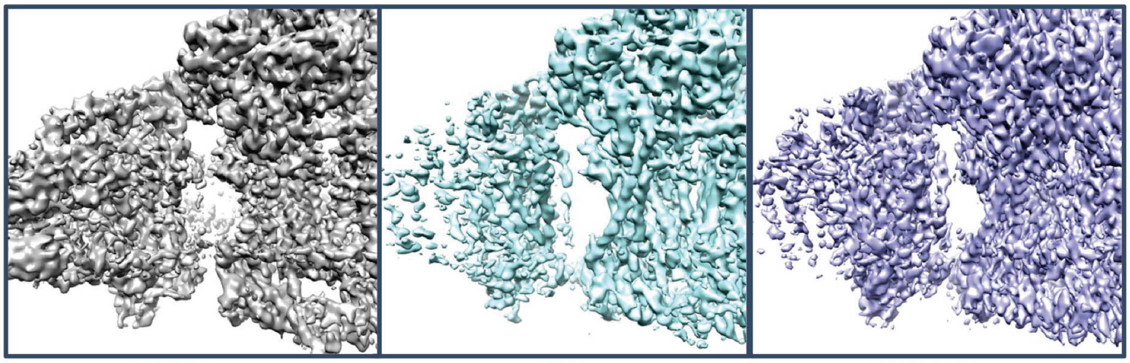

| ΔQCR6 | WT | WT full dataset |
|---|---|---|
| 81k particles | 81k particles | 493k particles |
| ~4.2Å resolution | ~4.5Å resolution | ~3.2Å resolution |

**(B)**

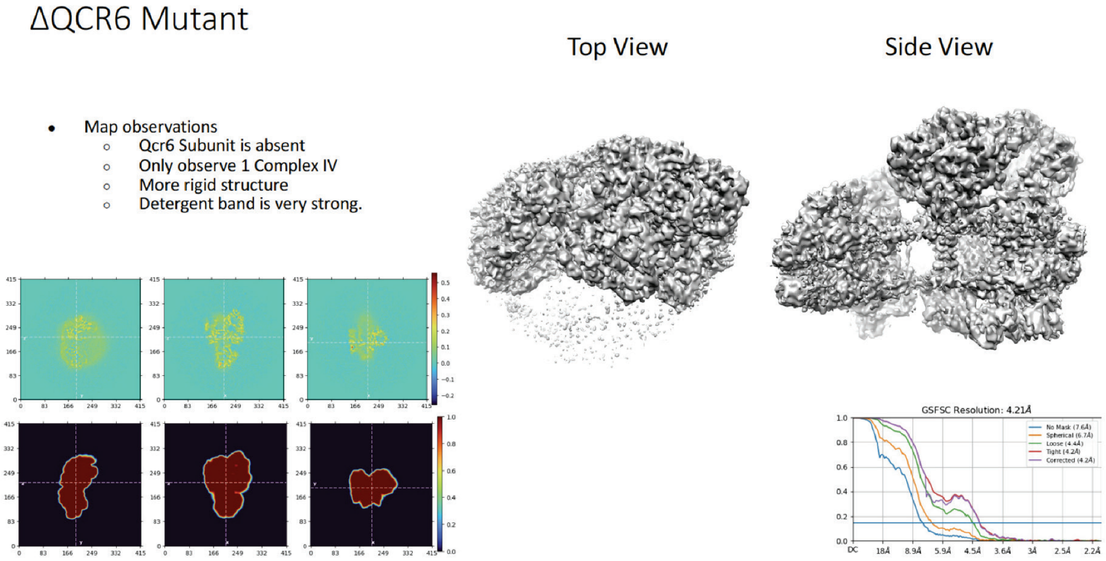

### SI figures for Figure 7

#### SI6.1: Hotspots for QCR6-cyt. *c* associations under the presence of CL, with conformations as being sampled by SMD.

QCR6's residues, which are with a percentage occupancy larger than 0.25, are highlighted in the molecular images as surface representations. The color of each surface representation follows the colorbar on the right and indicates the corresponding percentage occupancy.

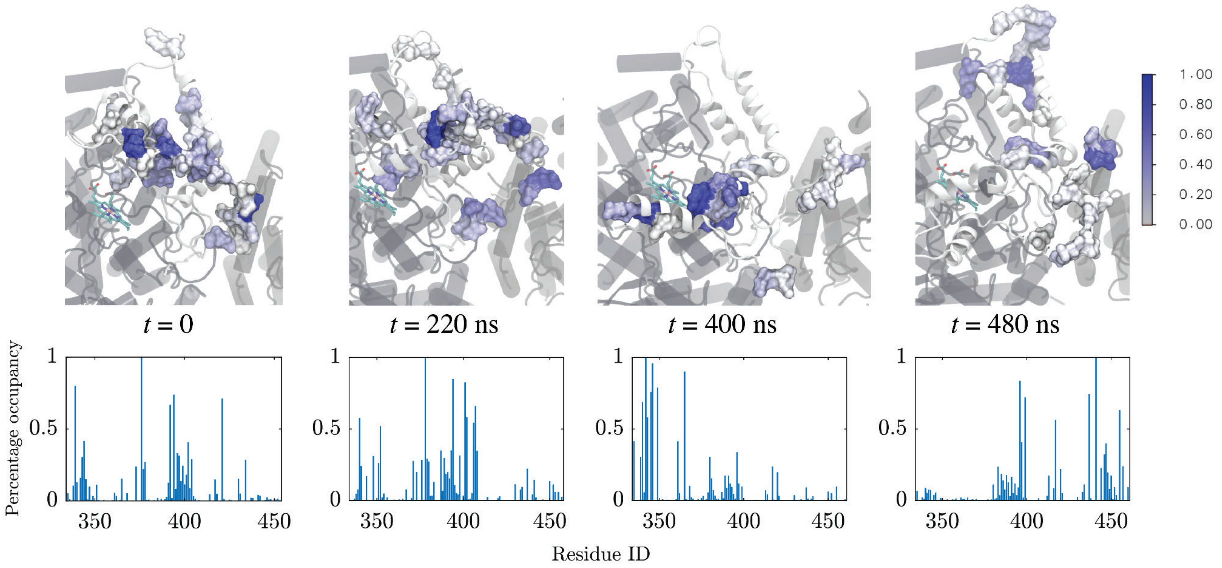


#### SI6.2 Impact of proposed mutations on cyt. *c*'s diffusion profile in the proximity of the supercomplex.

With WT denoting the wild type supercomplex, the electrostatic potential at -3 $k_BT$ around for the WT (blue, transparent, solid surface) and that for the mutant (red wireframe) are shown in (A). The mutant was created by implementing our design from **Fig. 7C** (including all selected mutations), which were expected to perturb the surface diffusion (2D diffusion) of cyt. *c* on the supercomplex. As being shown, our proposed mutations perturb the electronegativity of QCR6, inducing regions that are less electronegative. Here, a trajectory segment of cyt. *c* was considered experiencing 2D diffusion around QCR6 if the cyt. *c* diffused in a close proximity to the supercomplex's surface, namely within 6 Å heavy atom-to-heavy atom separation from either both QCR6 and CIII, or both QCR6 and CIV. Trajectory segments with cyt. *c* diffusing within 6 Å of heavy atom-to-heavy atom separation from QCR6, but not so with respect to CIII or CIV, were regarded as 3D diffusion around the supercomplex. The fraction of cyt. *c*'s trajectories demonstrating 2D diffusion around QCR6, shown in (B), was negligibly affected by our proposed mutations. However, the corresponding diffusion profiles, represented by occupancy maps for all diffusion trajectories around QCR6 in (C) with that for WT in blue and that for the mutant in red, are noticeably different. Compared with the profile of WT, the profile for the mutant is more spread out and extends away from the binding-interface for either CIII or CIV with cyt. *c*. This later geometric feature is demonstrated in (D) by distributions of the *z* coordinates of cyt. *c* contributing to each of the diffusion profiles in (C), with a smaller *z* coordinate indicating a closer proximity to the membrane (the membrane mid-plane is at $z = 0$Å).

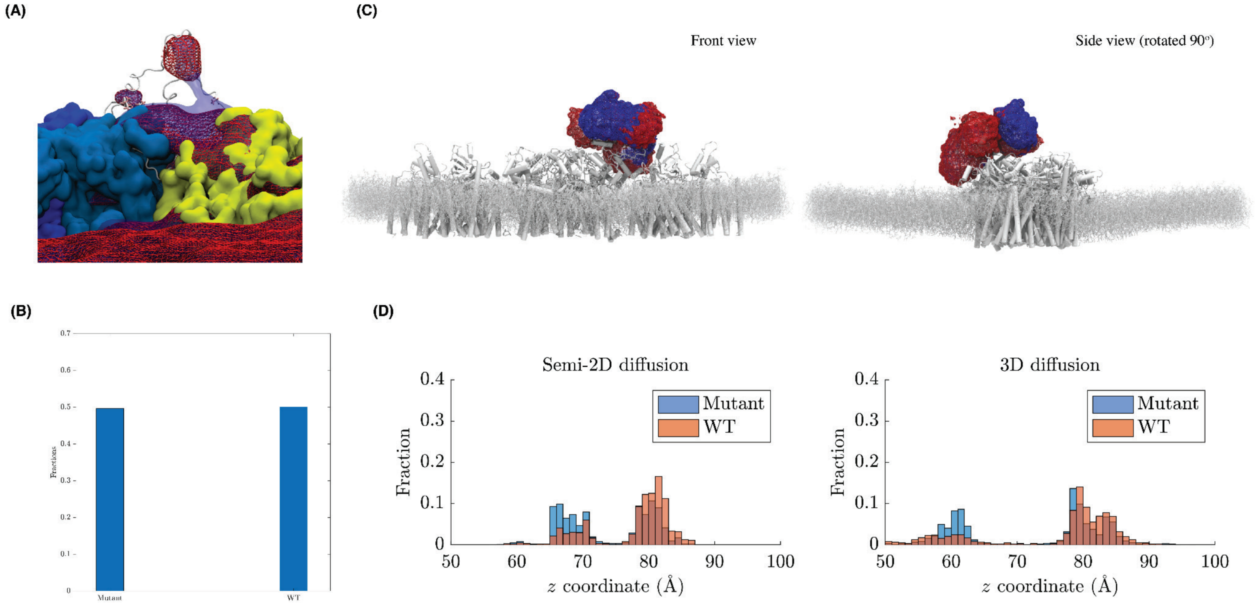


#### SI6.3 Tethers for membrane localization of charge carrier proteins from primitive heliobacterial species do not show QCR-like $CX_9C$ sequence signatures.

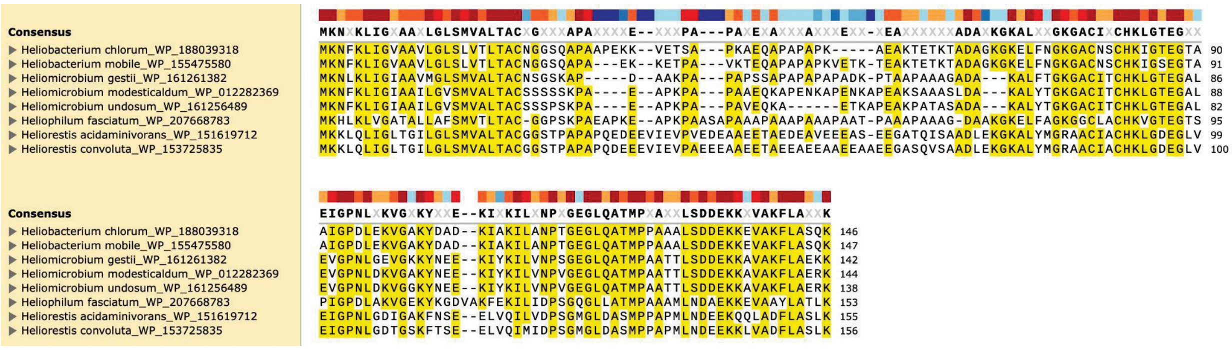

#### SI6.4 Cross-correlations (CC) between simulated cyt. *c* surface diffusion map and experimental data.

The distribution of CC coefficients, determined between each cyt. *c* pose (shown as protein backbones in dark blue), as being extracted from the sampled surface diffusion over SC, and the experimentally determined SC density map at the vicinity of cyt. *c* - CIII binding interface is shown below. Poses recorded with a poor cc (<= 0.1) are predominantly found either on top of CIII or CIV, reflecting the fact cyt. *c* tightly bound to either CIII or CIV was not included in the experimental data. Poses recorded with an average cc (> 0.1) are exclusively found in the region between CIII and CIV, highly overlapping with the location of Qcr6, suggesting the involvement of Qcr6 in the surface translocation of cyt. *c* over SC.

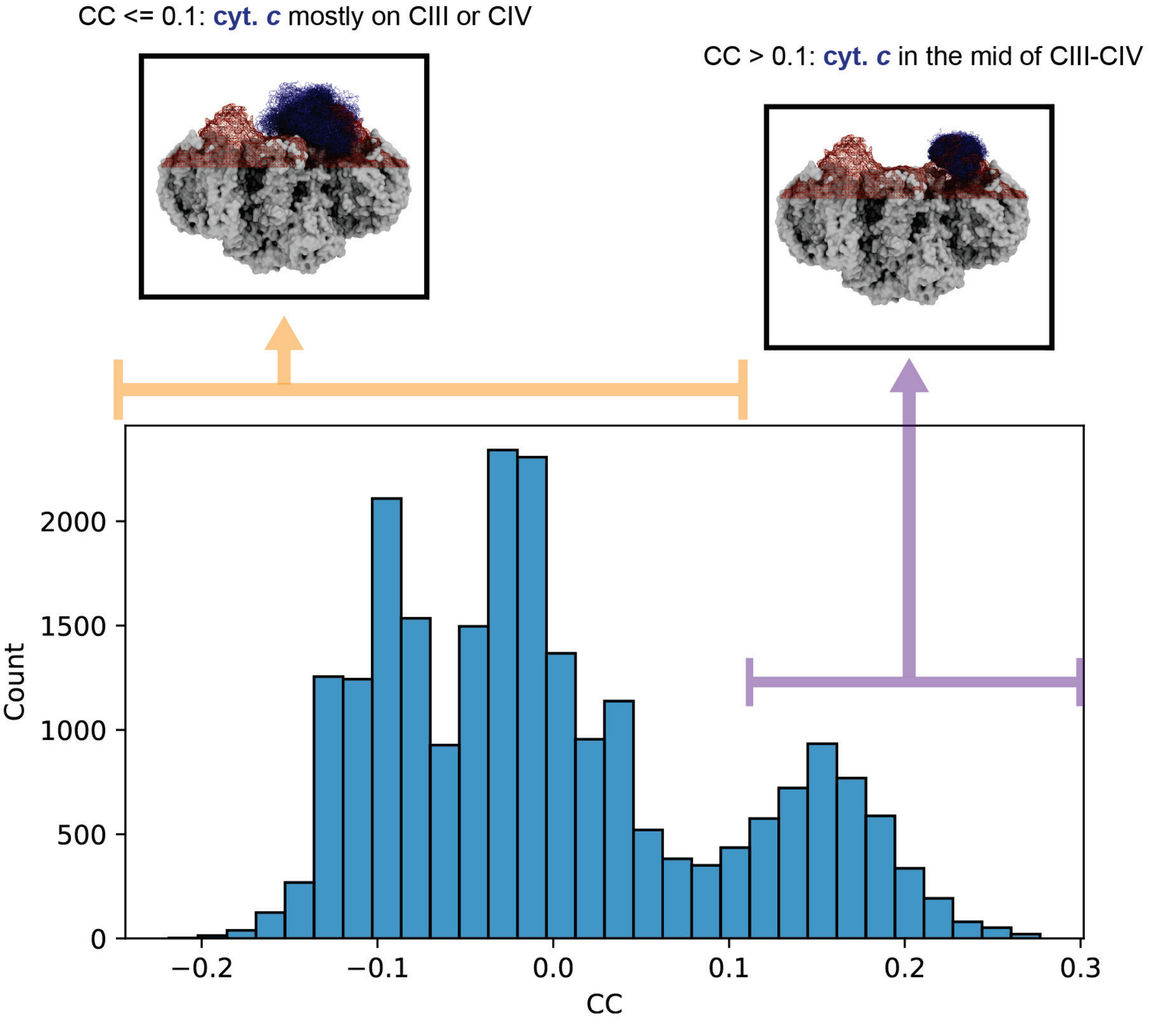

### SI figures for Methods

#### SI7.1 Workflow for modeling, simulations and analysis

CIII from the supercomplex model is docked using cyt.*c* following which, MELD simulations were performed to recover the structural ensemble of QCR6. The cyt.*c*-bound supercomplex model with complete QCR undergoes extended MD and BD simulations, kinetic parameters from which are fed into a Master equation model to determine relative ATP turnover by the WT and rearranged ΔQCR6 supercomplex.

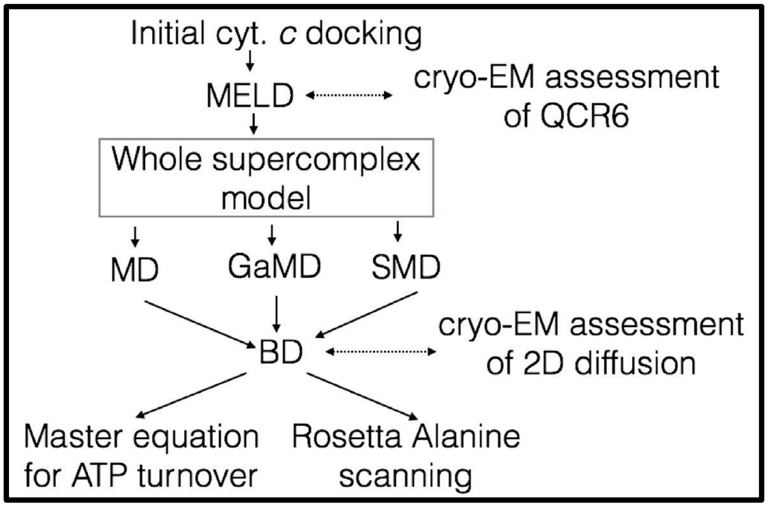

#### SI7.2 Modeling of QCR6 folds

The modeling of the disordered domain of QCR6 involves using part of the CIII and the whole QCR6. This combined system is shown in blue. Within this combined system, 2 lists of residues, one from CIII (orange) and one from QCR6 (green) were used to set restraints that guided the folding simulations of QCR6.

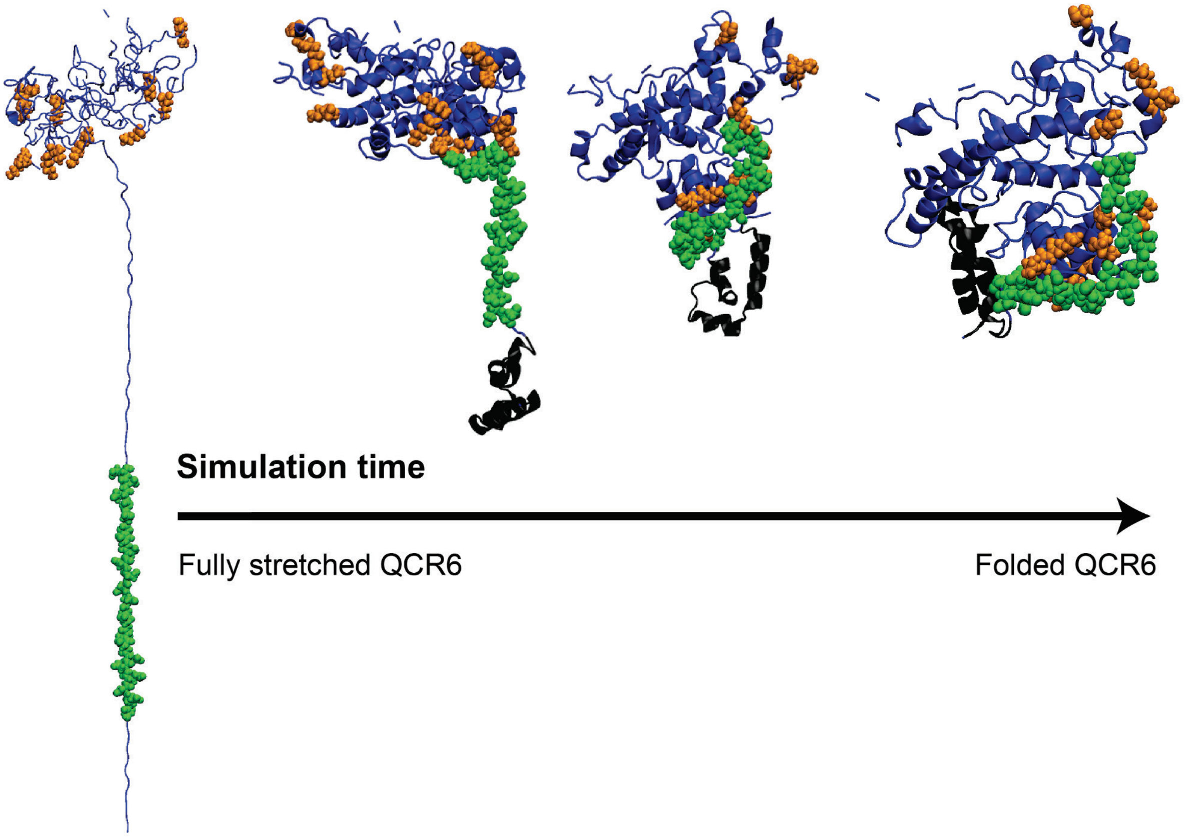

#### SI7.3: Spatial distributions of spaces with weak electropositivity and spaces with weak electronegativity.

Contour surfaces with weak electropositivity ($0.05k_BT$ per charge and $0.05k_BT$ per charge) for the wild type supercomplex are given in (A), with the color scheme of these surfaces given below the molecular image. Corresponding contour surfaces with weak electronegativity ($-0.05k_BT$ per charge and $-0.05k_BT$ per charge) are given in (B).

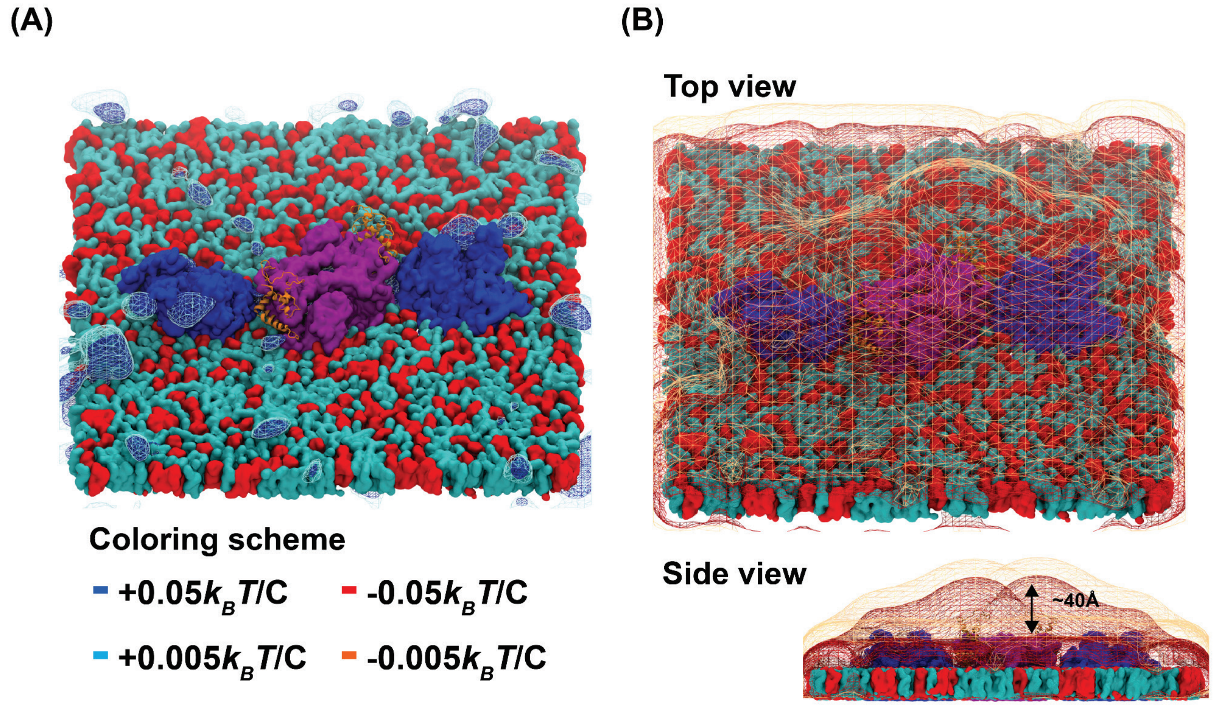

#### Simulation Table

Table 1: Table summarizing details of simulation involved in this study.

| Type of simulations | Systems simulated | Total simulation time |
|---|---|---|
| MELD | QCR6 + CIII | 500 ns |
| GaMD | QCR6 | 500 ns |
| MD | WT supercomplex embedded in PC-membrane<br><br>WT supercomplex embedded in PC & CL-membrane | 1 µs<br><br>1 µs |
| SMD | WT supercomplex embedded in PC-membrane | 3 x 500 ns |
| | WT supercomplex embedded in PC & CL-membrane | 3 x 500 ns |
| ARBD (14.08ms) | WT supercomplex embedded in PC-membrane | 0.64ms or 640 µs |
| | WT supercomplex embedded in PC & CL-membrane | 0.64 ms |
| | Mutant (ΔQCR6) supercomplex embedded in PC-membrane | 0.64 ms |
| | Mutant (ΔQCR6) supercomplex embedded in PC & CL-membrane | 0.64 ms |
| | Mutant ($QCR6_{Trunc}$) supercompelx embedded in PC-membrane | 0.64 ms |
| | Mutant ($QCR6_{Trunc}$) supercomplex embedded in PC & CL-membrane | 0.64ms |
| | WT supercomplex at 0 of SMD embedded in PC-membrane | 0.64ms |
| | WT supercomplex at 0 of SMD embedded in PC & CL-membrane | 0.64 ms |
| | WT supercomplex at 150 ns of SMD embedded in PC-membrane | 0.64 ms |
| | WT supercomplex at 150 ns of SMD embedded in PC & CL-membrane | 0.64 ms |

| | | |
|---|---|---|
| | WT supercomplex at 220 ns of SMD embedded in PC-membrane | 0.64 ms |
| | WT supercomplex at 220 ns of SMD embedded in PC & CL-membrane | 0.64 ms |
| | WT supercomplex at 400 ns of SMD embedded in PC-membrane | 0.64 ms |
| | WT supercomplex at 400 ns of SMD embedded in PC & CL-membrane | 0.64 ms |
| | WT supercomplex at 480 ns of SMD embedded in PC-membrane | 0.64 ms |
| | WT supercomplex at 480 ns of SMD embedded in PC & CL-membrane | 0.64 ms |
| | WT supercomplex of an ensemble average model over SMD electrostatics | 0.64 ms |
| | Mutant (Fig. 7) supercomplex embedded in PC & CL-membrane with QCR6's conformation at 0 of SMD for WT supercomplex embedded in PC & CL-membrane | 0.64 ms |
| | Mutant (Fig. 7) supercomplex embedded in PC & CL-membrane with QCR6's conformation at 150ns of SMD for WT supercomplex embedded in PC & CL-membrane | 0.64 ms |
| | Mutant (Fig. 7) supercomplex embedded in PC & CL-membrane with QCR6's conformation at 220 ns of SMD for WT supercomplex embedded in PC & CL-membrane | 0.64 ms |
| | Mutant (Fig. 7) supercomplex embedded in PC & CL-membrane with QCR6's conformation at 400 ns of SMD for WT supercomplex embedded in PC & CL-membrane | 0.64 ms |
| | Mutant (Fig. 7) supercomplex embedded in PC & CL-membrane with QCR6's conformation at 480 ns of SMD for WT supercomplex embedded in PC & CL-membrane | 0.64 ms |

### Analyses details

**Computations of binding propensity per unit area:** The per unit area binding propensity of cyt. *c* to a system *A*, ;labeled as $p_A$, is computed as,

$$p_A = \frac{Count_A}{SASA_A},$$

Where $Count_A$ is the number of associations between cyt. $c$ and $A$, and $SASA_A$ is the corresponding solvent accessible surface area (SASA). In this work, $A$ takes multiple forms, including membranes with various lipid compositions, the complete supercomplex, supercomplexes with different QCR6's conformations, and mutants of supercomplex.

For each of the different systems (*A*) employed, $Count_A$ was sampled from BD simulations. An association between cyt. *c* and *A* was considered to occur if any heavy atom of cyt. *c* came within 6 Å of a heavy atom of *A*. It is worth recalling that a positional fluctuation of 1.5 Å (see Method: Brownian Dynamics (BD)) had been imposed in our BD simulations. This positional fluctuation extended the spatial regime in which a cyt. *c* could sense forces, including the steric repulsive force, from a static binding target *A*, subsequently rendering a cyt. *c* harder to come into extremely close proximity with *A*. Thus, to compensate for this phenomenon, a 6 Å cutoff was used in place of the smaller, commonly used 4 Å cutoff in analyzing contact formations in MD simulations. Similarly, for each system *A*, the corresponding $SASA_A$ was computed using VMD [67] through its internal *measure sasa* function with a 6 Å cutoff.

**Per-residue percentage occupancy and percentage-occupancy weighted distributions:** For each set of BD simulations, the number of events ($N_i$) that a residue *i* makes contact with cyt. *c* was sampled. Assuming *i* is a member of domain ***I***, where ***I*** can be CIII or CIV, the percentage occupancy ($O_i$) of *i* is defined as,

$$O_i = \frac{N_i}{\max_{i \in I} N_i}.$$

Thus, $O_i$ measures how frequently *i* interacts (make contacts) with cyt. *c* with respect to the number of contacts achieved by the most frequent cyt. *c* - interacting residue from the same domain.

In **Fig. 4**, we are interested in the spatial distribution of cyt. *c* - interacting residues and their relevance to cyt. *c* - mediated electron transfer. To do so, for each cyt. *c* - interacting residue *i* in a domain ***I***, we determined the minimal separation ($d_i$) from *i* to the respective electron-transfer related cofactor. Then, we weighted the relevance of $d_i$ across the whole data set ($D = \{d_i\}_{i \in I}$) by the percentage occupancy of *i*, where a single data point $d_i$ is duplicated for $O_i \times \max_{i \in I} N_i$ times. After the weighting, a data point $d_i$ related to a residue *i* with a high $O_i$ will appear more frequently in *D*. This weighting scheme originates from the understanding that a residue *i* is spatially more relevant to electron transfer if it has both a small $d_i$ and a high $O_i$ simultaneously. The weighted data set *D* is then visualized in **Fig. 4** to show how the populations of cyt. *c* - interacting residues changes across the unfolding of Qcr6, as being characterized by their separations from electron-transfer linked cofactors,

**Cumulative density function for voxel values recorded in an electrostatic potential ($f^{elec}$):** The electrostatic potential for any system $A$ employed in this study was computed using Adaptive Poisson Boltzmann Solver (APBS) [68,69] with a resolution of 1Å in each spatial dimension. The resulting electrostatic potential $E_A(x, y, z)$, where $x, y, z$ span over the entire volume considered, was represented as a set of voxel values, $E_1, E_2, \cdots, E_N$ and was stored in the .dx file format. The index $i$ of each of these values$E_i$, can be converted back to the $x, y, z$ coordinates that it represents, and $N$ is the total number of grid points employed for the calculation.

A system's electrostatic field, once away from the source charges, decays quickly in a dielectric environment. For supercomplexes employed in our current study, spaces with electropositive potential below $0.05k_BT$ per charge are usually found outside the supercomplexes and are dispersed, behaving as fluctuations (**Fig. SI7.3A**). Spaces with the strength of electronegative potential below $0.05k_BT$ per charge are, on the other hand, space filling. Yet, these spaces are well separated from the surfaces of supercomplexes, with some of them located at a distance of around 40Å away from the supercomplex's surface (**Fig. SI7.3B**). Thus, voxel values lying between -0.05k$_B$T per charge and 0.05k$_B$T per charge primarily denote the non-atom-filling empty space involved in the electrostatic calculations for our systems. Also, it is worth noting that every atom in our systems is associated with a non-zero partial charge based on the Charmm36 force field [74]. These partial charges provide a quantum mechanical description for the classical behavior of atoms in molecular dynamics-like simulations and give non-zero electrostatic potential at the very close proximity of any atom. As a result, the number of voxel values lying between -$0.05k_BT$ per charge and $0.05k_BT$ per charge is more likely to represent the amount of void spaces involved in our calculations rather than the extent of the electroneutral region over the system's surface. Besides, a voxel value lying between -$0.05k_BT$ per charge and $0.05k_BT$ per charge will only modify the energy of a unit charge by less than $0.05k_BT$, which is 10% of the thermal energy associated with each degree of freedom for an atom. The impact of these voxel values on a charged object is thus relatively minor compared with the thermal fluctuations involved in Brownian dynamics. Therefore, with all these considerations, we discarded voxel values lying between -$0.05k_BT$ per charge and $0.05k_BT$ per charge when computing the cumulative distribution for voxel values of an electrostatic potential ($f^{elec}$).

After cleaning up our electrostatic data as mentioned above, we performed a histogram analysis on the remaining data. We computed the number of voxel values belonging to each bin of our histogram. The lower boundary value of a bin is -$20k_BT$ per charge and the upper boundary value is $20k_BT$ per charge. The bin width is $0.01k_BT$ per charge. The resulting histogram is then normalized so that the value for any bin $i$ ($B_i$) with a maximum/boundary value of $M_{B_i}$ represents the probability ($Pr\{E_j \leq M_{B_i}\}$) to find a voxel value less than $M_{B_i}$. Then, following the definition of cumulative density function, we employ the following approximation,

$$Pr\{E_j \leq M_{B_i}\} \approx \int_{b_i-0.005}^{b_i+0.005} f^{elec}(E)dE \Rightarrow f^{elec}(E = b_i) \approx \frac{Pr\{E_j \leq M_{B_i}\}}{0.01},$$

to obtain the CDF for voxel values of the electrostatic potential for our system, where $b_i$ is the mid-point value of bin $i$, and $b_i \pm 0.005$ gives the upper and lower boundary respectively. Thus, $M_{B_i} = b_i + 0.005$.

**Rate matrix computations and population transfer simulations:** To compute the rate matrix for any particular cyt. *c* - supercomplex system, the number of occurrences for a cyt. *c* transfer from one cyt. *c* bound state to another was first sampled from the corresponding BD simulations. There are 8 cyt. *c* bound states: 1) c3m1 (cyt. *c* associated with CIII of monomer 1 of the supercomplex); 2) c3m2 (cyt. *c* associated with CIII of monomer 2 of the supercomplex); 3) c4m1 (cyt. *c* associated with CIV of monomer 1 of the supercomplex); 4) c4m2 (cyt. *c* associated with CIV of monomer 2 of the supercomplex); 5) qcm1 (cyt. *c* associated with QCR6 of monomer 1 of the supercomplex); 6) qcm2 (cyt. *c* associated with QCR6 of monomer 2 of the supercomplex); 7) Membrane (cyt. *c* associated with the membrane); and 8) Bulk (cyt. *c* remaining in the bulk). These sampled numbers were then laid out in a matrix format with the columns representing the current cyt. *c* bound state and the row representing the next cyt. *c* bound state. An example of such matrices, termed here as occurrence matrices to facilitate disucssions, is given in **Fig. SI3.2** for the WT.

To derive the rate matrix, modifications were applied to the corresponding occurrence matrix, which derivations are outlined above. For any occurrence matrix $O$ with its elements $O_{ij}$, where $i, j$ are indices for the row the column respectively, each matrix element $R_{kl}$ for the corresponding rate matrix $R$ is given by

$$R_{kl} = \frac{O_{kl}}{\sum_{i=1}^{8} O_{ki}} .$$

The change on the population of a bound state $k$ at time $t$, labeled as $\rho_k(t)$, over a time interval $\Delta t$, which was used to sample the occurrence matrix, can be computed from the corresponding rate matrix as:

$$\dot{\rho}_k(t)\Delta t = \sum_{i=1}^{8} R_{ki}\rho_i(t) - \sum_{i=1}^{8} R_{ik}\rho_k(t).$$

The above equation does not bear an analytical solution. Thus, we integrate the above equation numerically as,

$$\rho_k(t+\Delta t) = \rho_k(t) + \dot{\rho}_k(t)\Delta t \ .$$

The above integration was performed for 500 steps with a time interval of 20 ns, a value used in parsing the BD simulations into MD-like trajectories. As we want to focus the study on population transfer from CIII to CIV, an initial population condition where cyt. *c* is equally likely to be found across all different components of CIII was employed. Vector elements of this initial population condition are:

$$\rho_{c3m1}(0) = \rho_{c3m2}(0) = \rho_{qcm1}(0) = \rho_{qcm2}(0) = 0.25,$$
$$\rho_{c4m1}(0) = \rho_{c4m2}(0) = \rho_{Membrane}(0) = \rho_{Bulk}(0) = 0.$$

After obtaining $\rho_k(t)$ for all 8 different cyt. *c* bound states based on the above initial condition, the results were concatenated to highlight population changes for bound states related to CIII, CIV, the membrane, and the bulk solvent. The concatenation was done as:

$$\rho_{CIII}(t) = 0.25 \times (\rho_{c3m1}(t) + \rho_{c3m2}(t) + \rho_{qcm1}(t) + \rho_{qcm2}(t))$$
$$\rho_{CIV}(t) = 0.5 \times (\rho_{c4m1}(t) + \rho_{c4m2}(t))$$
$$\rho_{Membrane}(t) = \rho_{Membrane}(t)$$
$$\rho_{Bulk}(t) = \rho_{Bulk}(t)$$

The result of the above concatenation concerning population transfer starting from CIII to CIV is given in **Fig. SI3.2**.

**Estimating the ATP production rate:** For a power input of $\lambda$, the rate of ATP output ($K_{ATP}$) for a linear electron transport chain is approximately given by [75]:

$$K_{ATP} = \frac{\lambda}{2}(1 + \frac{\lambda\tau}{2N})^{-1},$$

where $N$ is the number of electron acceptor complex per donor complex (which is 1 in our case), $\tau$ is the rate of electron turnover. Thus, at large values of $\lambda$ and constant donor-stoichiometry, $K_{ATP} \propto \tau^{-1}$, implying $\frac{K_{ATP,WT}}{K_{ATP,\Delta QCR6}} \propto \frac{\tau_{\Delta QCR6}}{\tau_{WT}}$, which we present in **Fig. 7A**.

**Discussions on membrane-mediated cyt. *c* diffusion to CIII:** When the anionic lipids are removed from the membrane model, surface coverage of the 50-80%-attenuated cyt. *c* population on the WT and ΔQCR6 supercomplex still increases by 2-fold (**Fig. SI4.6**), namely from 1.1 cyt.*c*/Å$^2$ to 2.1 cyt.*c*/Å$^2$ for the former and 0.7 cyt.*c*/Å$^2$ to 1.5 cyt.*c*/Å$^2$ for the later. These increments in cyt. *c* coverage does not necessarily indicate an enhanced ability of the supercomplex to recruit more cyt. *c*. Rather, it reveals a repartition of the surface cyt. *c* population towards the supercomplex instead of the membrane, only after a significant amount of the carriers are lost to the bulk.

Since our simulations were performed with a finite diffusion space, most cyt. *c* would be able to find an association spot on the supercomplex-membrane system given a long enough simulation time. The residence times of cyt. *c* on the supercomplex, associating without any lipid mediation, are also in line with this view: the cyt. *c* residence on the WT marginally decreases due to removal of the cardiolipins (**Fig. SI4.7B**) and that for ΔQCR6 is less dependent on the cardiolipin content as a significant fraction of the carriers remain in the bulk (**Fig. SI4.5**). In fact, upon the removal of anionic lipids, direct cyt. *c* - membrane interactions go down almost by 2.8-fold (**Fig. SI4.6**), and the passage time of the cyt. *c* from the bulk (150 Å away from the membrane) to the supercomplex also increases (**Fig. SI4.8**). Therefore, the membrane surface-mediated mechanism of cyt. *c* transfer, often referred to as 2D diffusion mechanism [1,4], wherein the carrier is first brought to the membrane vicinity and then interacts with CIII slows down when either the QCR6 or the cardiolipin are absent.

### Movie captions

#### Movie 1: MELD

The folding process of a fully stretched QCR6 (orange) to its interacting conformation with CIII (red) and CIV (blue) under the presence of a tightly bound cyt. *c* (green) on CIII was explored by MELD (Modeling employing limited data) sampling simulation.

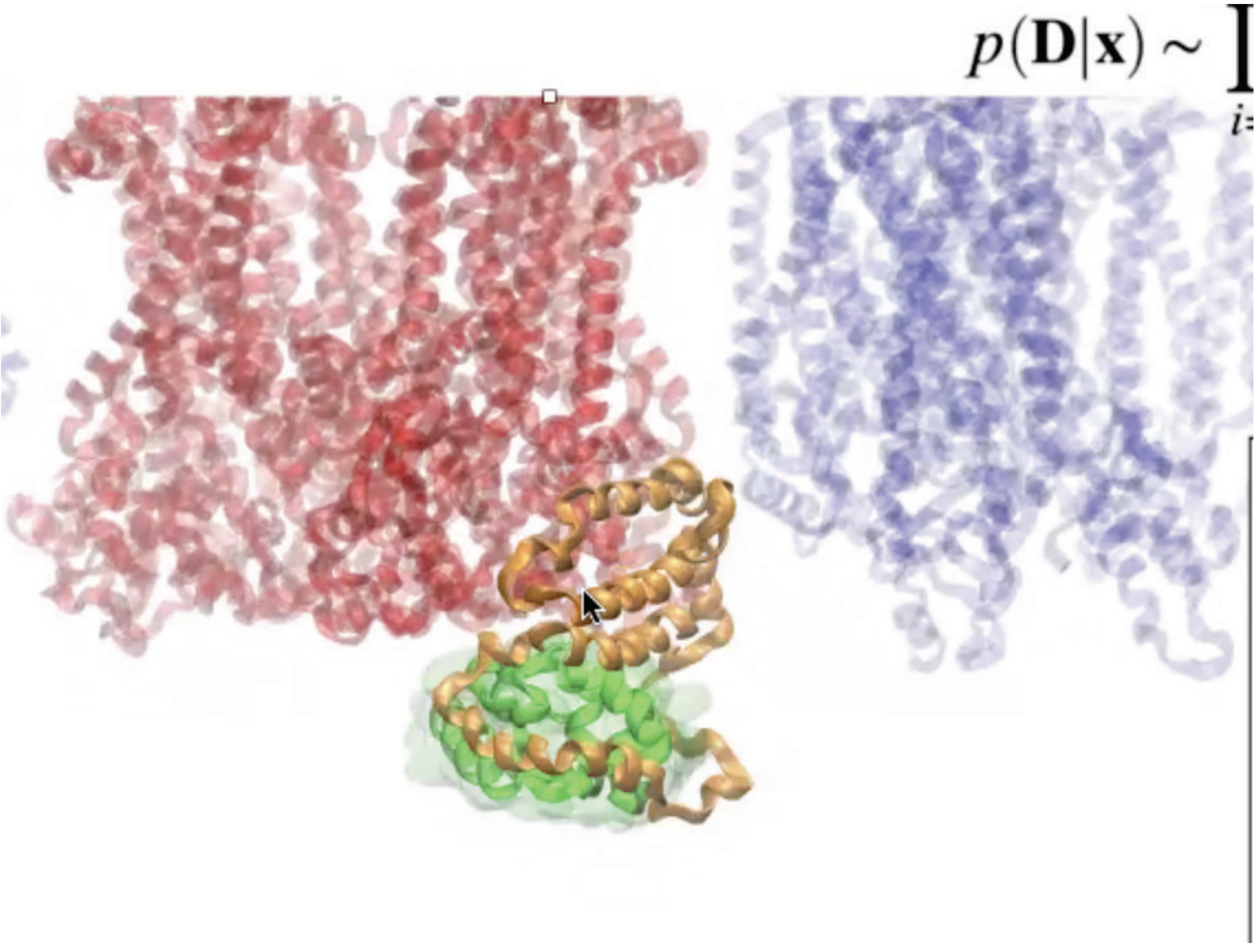


### Movie 2: SMD

Potential QCR6 conformations along the translocation of cyt. *c* from CIII to CIV were sampled by SMD (steered molecular dynamics) simulations. During this simulation, cyt. *c*'s motion was biased towards directional motion from CIII to CIV with the dynamics of QCR6 unrestricted.

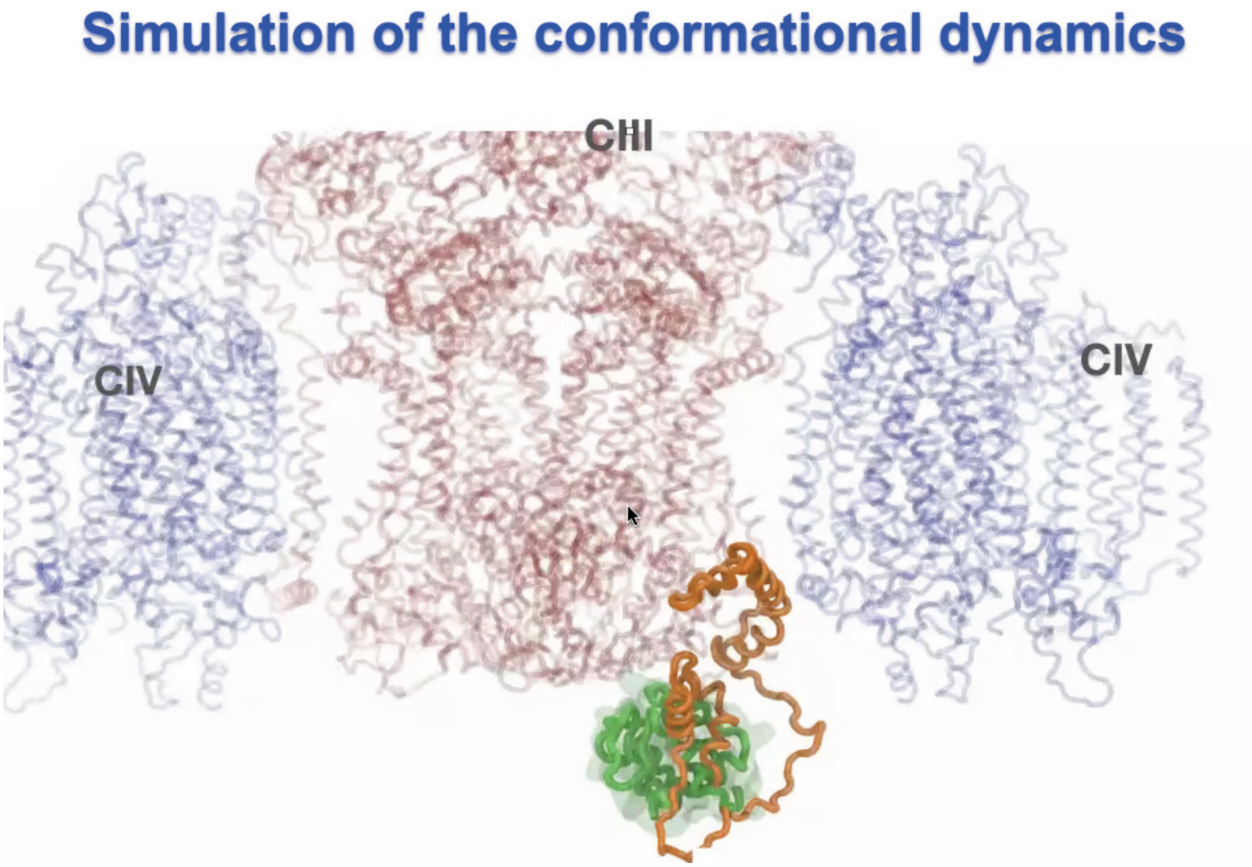


Movie 3: ARBD

The shift of popular association spots of cyt. *c* on the supercomplex (SC) (exhibited by the protein's occupancy map on SC) was sampled by ARBD (atomic-resolution brownian dynamics) simulations, with SC-QCR6 structures representing a directional translocation of cyt. *c* from CIII to CIV serving as the underlying inputs. These structures were sampled by SMD and were each explored by ARBD through a 0.64ms-long simulation.

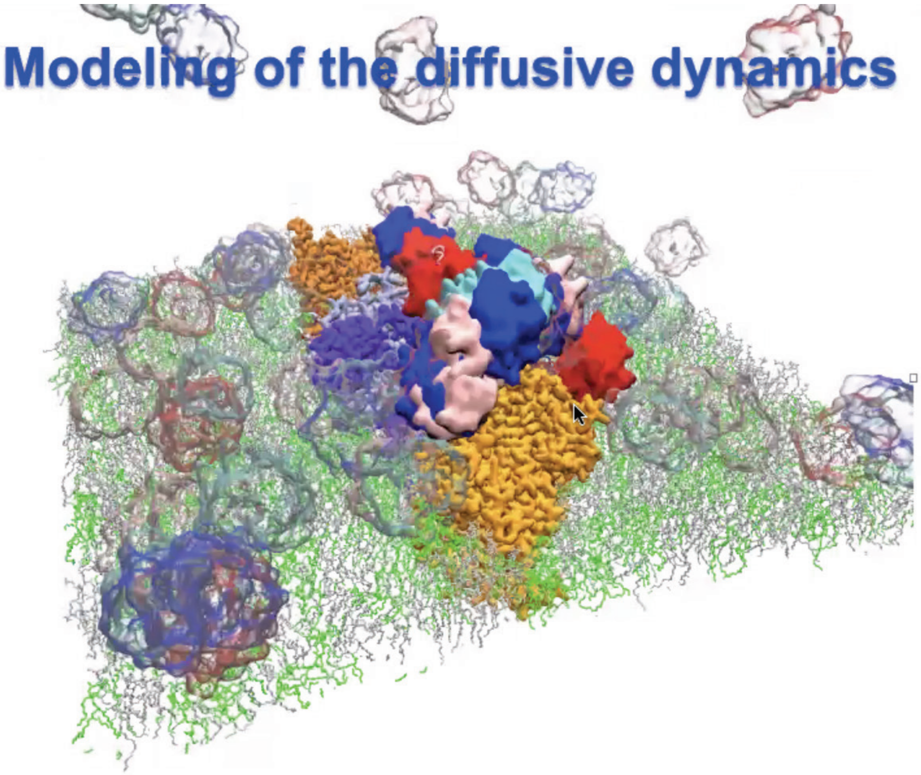
Modeling of the diffusive dynamics
